# A trove of antiviral TRIM family E3 ligases in reptiles

**DOI:** 10.1101/2025.06.23.661133

**Authors:** Ian N. Boys, Meghan R. Quinlan, Nels C. Elde

## Abstract

Scaled reptiles (squamates) naturally encounter orthoflaviviruses and in some cases robustly resist infection. To identify antiviral defenses encoded in reptile genomes, we screened a green iguana cDNA library and discovered a reptilian TRIM-family E3 ubiquitin ligase that reduces dengue virus replication ∼10,000-fold in human cells. Experimental evolution identified orthoflavivirus capsid as a key determinant of harbiTRIM-mediated restriction, revealing a convergence of TRIMs on viral capsids as targets during viral infections.

HarbingerTRIM is situated near an endogenized *Harbinger* transposon in various copy numbers in squamate genomes. Analysis of a sampling of reptile harbiTRIM variants revealed distinct antiviral properties, highlighting the immune diversity present in non-mammal animal genomes as a largely untapped reservoir of antiviral defenses.

## Introduction

Beyond mammals, relatively little is known about specific mechanisms underlying metazoan protein based antiviral immunity (Fig. 1A and fig. S1A). Scaled reptiles (squamates) are the largest order of terrestrial vertebrates, with over 12,000 extant species (*1*). This ancient lineage emerged 200–250 million years ago and comprises billions of years of evolution, including at host-pathogen interfaces (*2-4*). Studies have shown that some squamates such as iguanas resist infection by orthoflaviviruses (*5, 6*), a group of medically important mosquito-transmitted arboviruses that collectively infect over 400 million people annually (*7*). These viruses include emerging and re-emerging pathogens such as dengue, Zika, West Nile, and yellow fever viruses and are expected to dramatically increase in prevalence as global climate trends modify the ranges of the mosquitos that vector them (*7*). The mechanisms underlying orthoflavivirus resistance by squamates are unknown (Fig. 1A).

**Figure 1.**
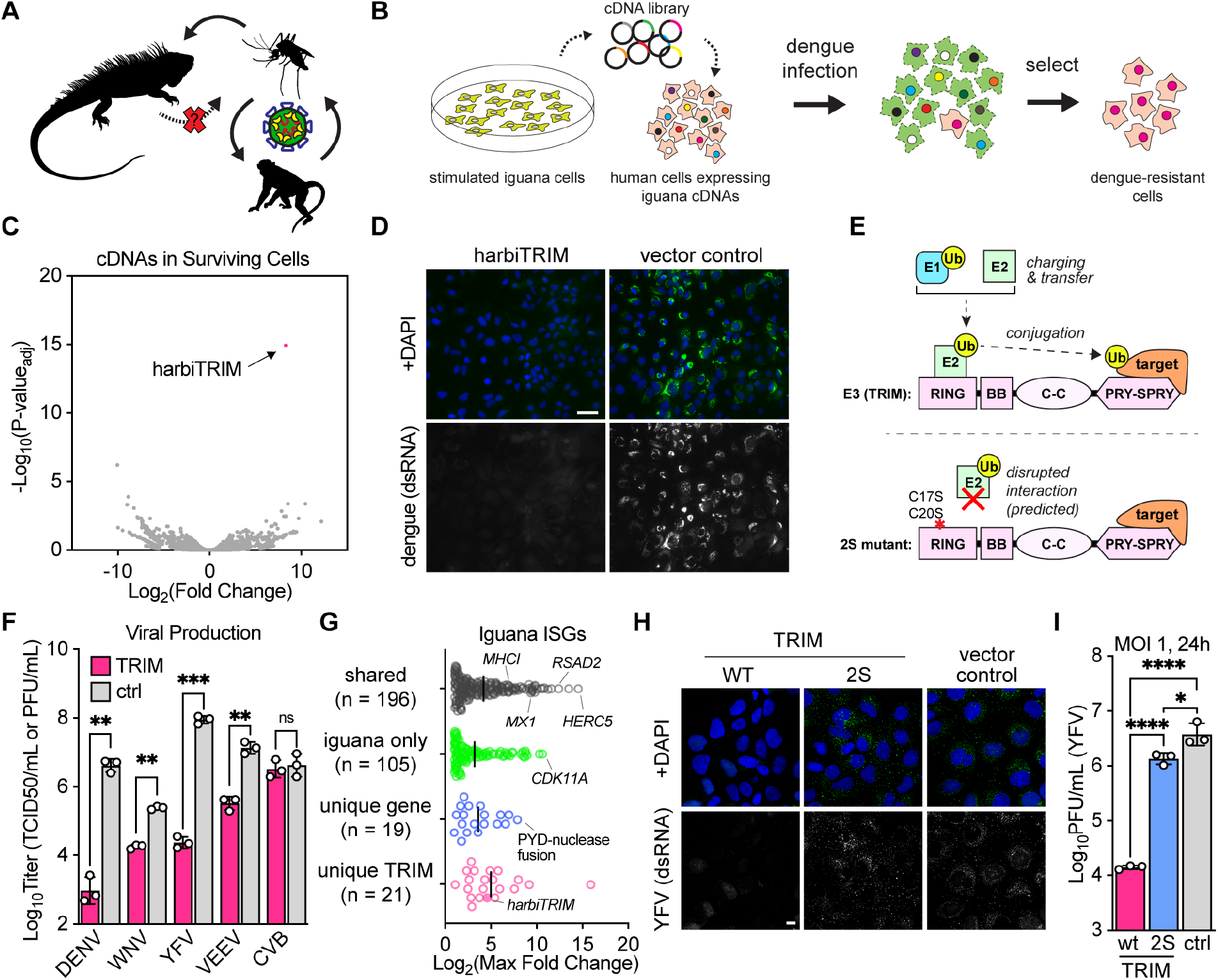
Identification of harbiTRIM as a flavivirus restriction factor in iguanas. (**A**) Iguanas and other lizards have been reported to resist infection by some vector-borne flaviviruses. (**B**) Restriction factor screen schematic. (**C**) Results of dengue restriction factor screen. (**D**) Huh7 cells stably expressing harbiTRIM or a vector control were infected with DENV at an MOI of 1 for 48 hours and stained for double-stranded RNA. Scale bar: 50 μm (**E**) Overview of the canonical ubiquitylation cascade, including a schematic of a class IV TRIM E3 ligase and a predicted loss-of-function mutant (2S). Class IV TRIMs contain a RING domain involved in ubiquitin transfer, B-box and coiled-coil domains, and a PRY-SPRY domain involved in substrate recognition. (**F**) Viral production from Huh7 cells stably expressing harbiTRIM or a vector control following single-cycle infections with DENV (48 hours), WNV and YFV (24 hours), VEEV and CVB (6 hours). Infections were performed at an MOI of 1 except CVB (MOI of 10) as Huh7 cells are less permissive to CVB. (**G**) Maximum transcript induction following treatment of IGH2 cells with iguana IFN for 6 or 24 hours. (**H**) Huh7 cells expressing harbiTRIM, the 2S mutant, or a vector control were infected with YFV at an MOI of 5 and stained for double-stranded RNA after 24 hours. Scale bar: 10 μm. (**I**) Viral production following infection of Huh7 cells stably expressing harbiTRIM, the 2S mutant, or a vector control with YFV at an MOI of 1 for 24 hours.

Our shared heritage of cell-autonomous innate immunity involving interferon signaling points to hundreds of candidate genes that might contribute to antiviral defenses in squamates. We used genetic screens of cDNA libraries enriched for interferon stimulated genes (ISGs), experimental evolution, and comparative analyses to discover a collection of antiviral TRIM-family E3 ubiquitin ligases that potently inhibit diverse pathogenic orthoflaviviruses. The newly identified TRIM from green iguana functions through an apparent capsid-directed process, suggesting that viral capsids represent a recurring target of antiviral TRIM proteins. Because orthoflavivirus entry requires capsid ubiquitylation, capsid may represent a vulnerable point for antiviral intervention. Sampling the family of related TRIMs among reptiles revealed distinct profiles of antiviral activity for multiple viruses. These divergent TRIMs comprise a versatile assortment of antiviral proteins and provide a new perspective for studying the evolutionary potential of antiviral TRIM proteins across vertebrates.

## Results

### harbingerTRIM is an orthoflavivirus restriction factor

To identify specific antiviral genes present in the iguana, we generated a cDNA library enriched for ISGs from iguana cells (table S1) and screened for cDNAs that protect human cells from infection by the orthoflavivirus dengue virus (Fig. 1B). Our screen identified a single cDNA that robustly protects cells from dengue virus: a TRIM-family E3 ligase proximal to an endogenized *Harbinger* transposon element, hence referred to as harbingerTRIM (or harbiTRIM for short, Fig. 1, C and D). TRIM E3 ligases are ubiquitin conjugating enzymes that provide substrate specificity as the terminal ligase in the ubiquitylation cascade, a regulatory process that is important in many aspects of cell biology, including immune defenses (*8*) (Fig. 1E).

We assessed the breadth of harbiTRIM antiviral activity against representative RNA viruses including the flaviviruses dengue (DENV), West Nile (WNV), and yellow fever (YFV) viruses, as well as the distantly related arbovirus Venezuelan equine encephalitis virus (VEEV, an alphavirus), and the picornavirus coxsackievirus B3 (CVB) by ectopic expression in human cells. HarbiTRIM restricted DENV and YFV by over 1,000-fold, WNV and VEEV by over 100-fold, but did not inhibit CVB (Fig. 1F). While inhibition of an alphavirus suggests that the antiviral activity of harbiTRIM extends beyond orthoflaviviruses, we focused on its ability to inhibit orthoflaviviruses, towards which it was broadly inhibitory.

Like many antivirals, we found that harbiTRIM is upregulated by interferon in iguana cells (Fig. 1G and fig. S1 B–D). Using publicly available data, we also determined that harbiTRIM is expressed in a broad range of tissues in vivo, any of which could be relevant for viral pathogenesis (*7*) (fig. S1E). Alongside harbiTRIM, numerous genes were upregulated by IFN in iguanas (Fig. 1G, fig. S1, C and H), including many that are shared with mammals. There are additional shared genes that display interferon regulation in iguanas but not mammals, and other genes, including an assortment of TRIMs, that are unique to reptiles, suggesting potential antiviral innovations in squamates in addition to harbiTRIM.

Ubiquitin is required for orthoflavivirus uncoating and genome release (*9, 10*), complicating efforts to investigate the antiviral phenotype of a TRIM E3 ligase that inhibits orthoflaviviruses. For example, treating cells with a proteasome inhibitor such as MG132 before or during viral entry blocks infection (*11*). To sidestep this issue, we generated a predicted E2-binding mutant based on conserved cysteine residues in the RING domain of harbiTRIM (*12*) (Fig. 1E). This mutant (2S), which expressed at equivalent or higher levels than wild-type harbiTRIM (fig S1, F and G), exhibited a nearly complete loss of antiviral activity towards YFV when ectopically expressed, suggesting that harbiTRIM requires an intact RING domain to inhibit orthoflaviviruses (Fig 1, H and I).

### Experimental evolution identifies capsid as a target of harbiTRIM

To identify the potential viral target of harbiTRIM, we performed serial infections of yellow fever virus in human cells overexpressing harbiTRIM. Viral titers increased after several passages, hinting at virus adaptation to evade harbiTRIM activity (Fig. 2A). Long-read sequencing revealed two nonsynonymous mutations at nearly 100% frequency in the capsid protein of this population, suggesting that it may be targeted by harbiTRIM (Fig. 2B and fig S2A). Sequencing of earlier passages of virus showed that these mutations independently arose in separate genomes (Fig. 2C and data S6). A genome containing both mutations – likely the result of a second point mutation on either background since recombination is exceedingly rare in orthoflaviviruses (*13*) – arose later, rapidly sweeping the population (Fig. 2C). We introduced each of these mutations into a clean genetic background and in combination to examine viral replication in the presence of harbiTRIM. Independently, each mutation modestly enhanced replication in harbiTRIM-expressing cells, while the virus containing both mutations replicated to over 50-fold higher levels than parental virus. This phenotype was specific to harbiTRIM, as all viruses replicated near parental levels in control cells (Fig. 2, D and E). Together, these results implicate capsid as a target for harbiTRIM. Capsid is a multi-function protein that protects the orthoflavivirus genome during infection, facilitates the loading of nascent genomes into virions during assembly, and is the first component of the viral polyprotein to be translated (Fig. 2F).

**Figure 2.**
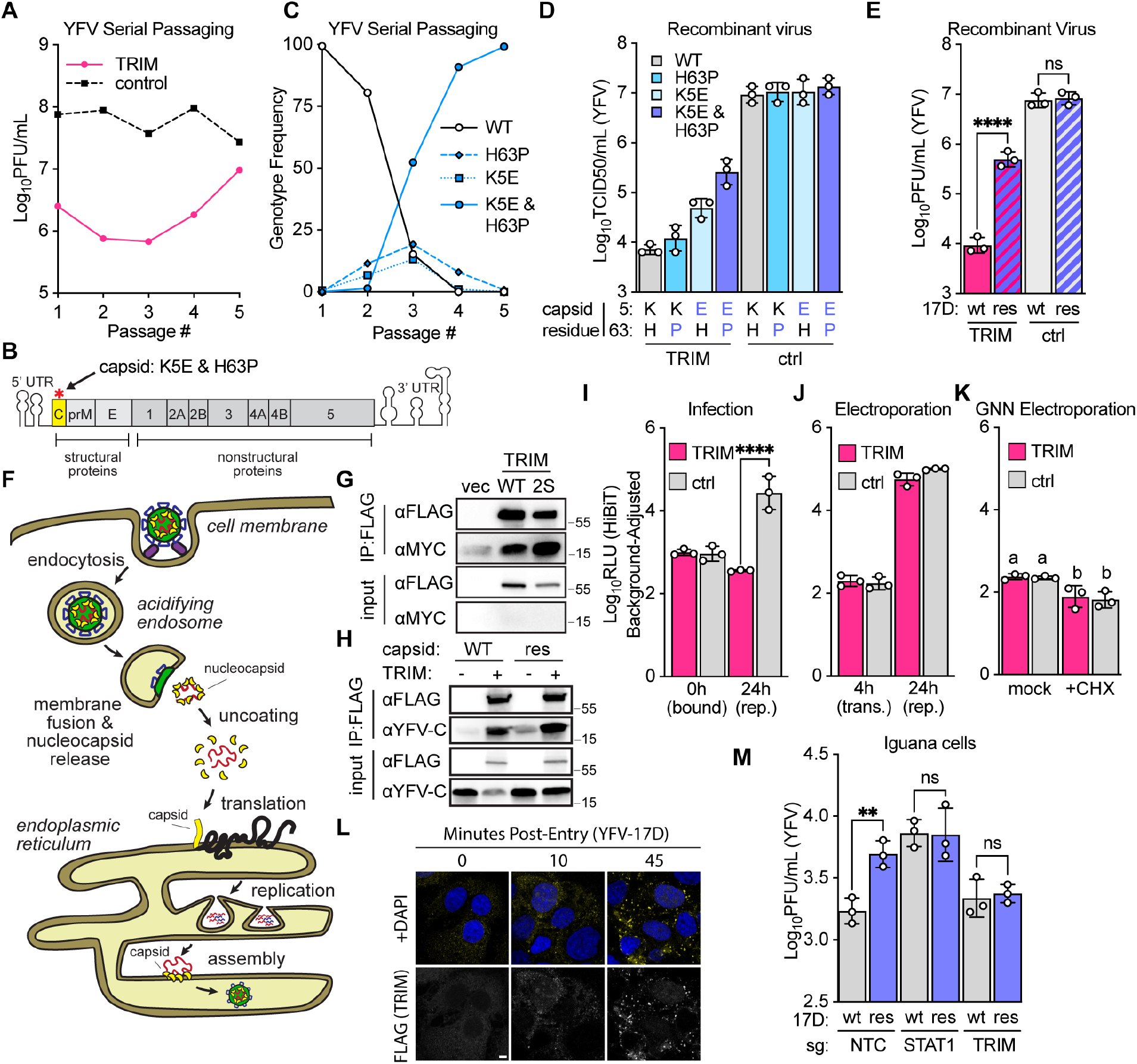
harbiTRIM restricts flavivirus infection at an early capsid-dependent step. (**A**) YFV-17D was serially passaged in Huh7 cells expressing harbiTRIM or a vector control (starting MOI of 100). (**B**) Schematic of the YFV-17D genome, indicating resistance mutations. (**C**) Viral genotype frequencies during passaging. (**D**) Viral production following infection of Huh7 cells expressing harbiTRIM or a vector control with HiBiT YFV-17D bearing the indicated mutations at an MOI of 1 for 24 hours. (**E**) Huh7 cells expressing harbiTRIM or a vector control were infected with YFV-17D bearing both mutations (res) or wild-type (wt) virus at an MOI of 1 for 24 hours. (**F**) Overview of early steps of flavivirus replication. (**G** to **H**) FLAG immunoprecipitation of 293T cells co-transfected with Myc-tagged (G) or untagged (H) YFV-17D capsid and FLAG-tagged wild-type harbiTRIM (wt), the 2S mutant, or a vector control. (**I**) Viral protein levels measured by luciferase assay during a synchronized infection of Huh7 cells expressing harbiTRIM or a vector control with YFV-17D-HiBiT at an MOI of 25. (**J** to **K**) Huh7 cells expressing harbiTRIM or a vector control were electroporated with YFV-17D-HiBiT RNA (J) or YFV-17D-HiBiT-GNN RNA (K), and viral protein levels were determined by luciferase assay. In (K), cells were cultured in the presence or absence of cycloheximide (100 μM). (**L**) Cells were synchronously infected with YFV-17D at an MOI of 50 and fixed for immunofluorescence at the indicated time points post-shift to 37°C. Scale bar: 5 μm (**M**) Bulk CRISPR-targeted iguana cells were infected for two days with the indicated viruses. harbiTRIM editing efficiency was 96.3–100% (Data S6).

To test for interactions between harbiTRIM and orthoflavivirus capsid, we expressed both in cells and found that they co-immunoprecipitate (Fig. 2, G and H and fig. S2, C and D). The RING domain mutant (2S), which loses antiviral activity (Fig 1, H and I), retains its ability to complex with capsid, indicating that association with capsid alone is insufficient for antiviral activity. Accordingly, we next characterized the interaction between harbiTRIM, capsid, and ubiquitin through ubiquitin pulldown assays. Although ubiquitylated harbiTRIM complexed with capsid and ubiquitin in lysates (fig. S2, G and H), we did not detect ubiquitylated capsid in these experiments, despite evidence that harbiTRIM itself was polyubiquitylated (fig. S2, E and F).

We next sought to determine how harbiTRIM targeting of capsid impacts orthoflavivirus infection. We focused on early steps of infection since dsRNA, a hallmark of viral replication complexes, is not observed during orthoflavivirus infection of harbiTRIM-expressing cells (Fig. 1, D and H). We generated a reporter yellow fever virus encoding a split luciferase (HiBiT) (*14*) as part of its capsid protein (fig. S2B). The engineered virus bound harbiTRIM-expressing cells at similar levels to control cells (Fig. 2I), but we observed a substantial reduction in luciferase activity in the presence of harbiTRIM during later stages of infection, consistent with our observations with wild-type virus (Fig. 1F and Fig. 2E).

To directly test late stages of replication, we bypassed viral entry by electroporating naked viral RNA into cells and observed no significant differences in viral protein translation or production during replication in knockout cells (Fig. 2M and fig. S2, I–K). The absence of a detectable knockout phenotype indicates that harbiTRIM is not a dominant determinant of YFV restriction in iguana cells under these conditions. However, the harbiTRIM resistant virus replicated to higher levels in iguana cells compared to wild-type virus (Fig. 2M), suggesting that escape of harbiTRIM-mediated restriction is advantageous during infection of iguana cells. This virus replicates at wild-type levels in harbiTRIM knockout iguana cells, consistent with this advantage being harbiTRIM-dependent. At later time points during infection, titers of both harbiTRIM-resistant and wild-type viruses declined in all IFN-competent cell populations, consistent with additional IFN-dependent antiviral mechanisms contributing to restriction of orthoflaviviruses in iguana cells (fig. S2J).

### A versatile collection of antiviral E3 ligases in squamate reptiles

Selective pressures from pathogens often drive the diversification of antiviral proteins. While the iguana genome encodes only one harbiTRIM, other squamates encode between zero to five paralogs (Fig. 3, A and B and fig. S3A) in a complex, repeat-rich genomic neighborhood anchored by the presence of an endogenized and transcriptionally active *Harbinger* transposable element (fig. S3, A–C). An analysis of this locus in thirteen representative squamate genomes revealed extensive variability, with multiple gains and losses of genes encoding PRY-SPRY domains including TRIMs, butyrophilins, and PRY-SPRY domain-only proteins reminiscent of thaicobrin, a component of snake venom.

**Figure 3.**
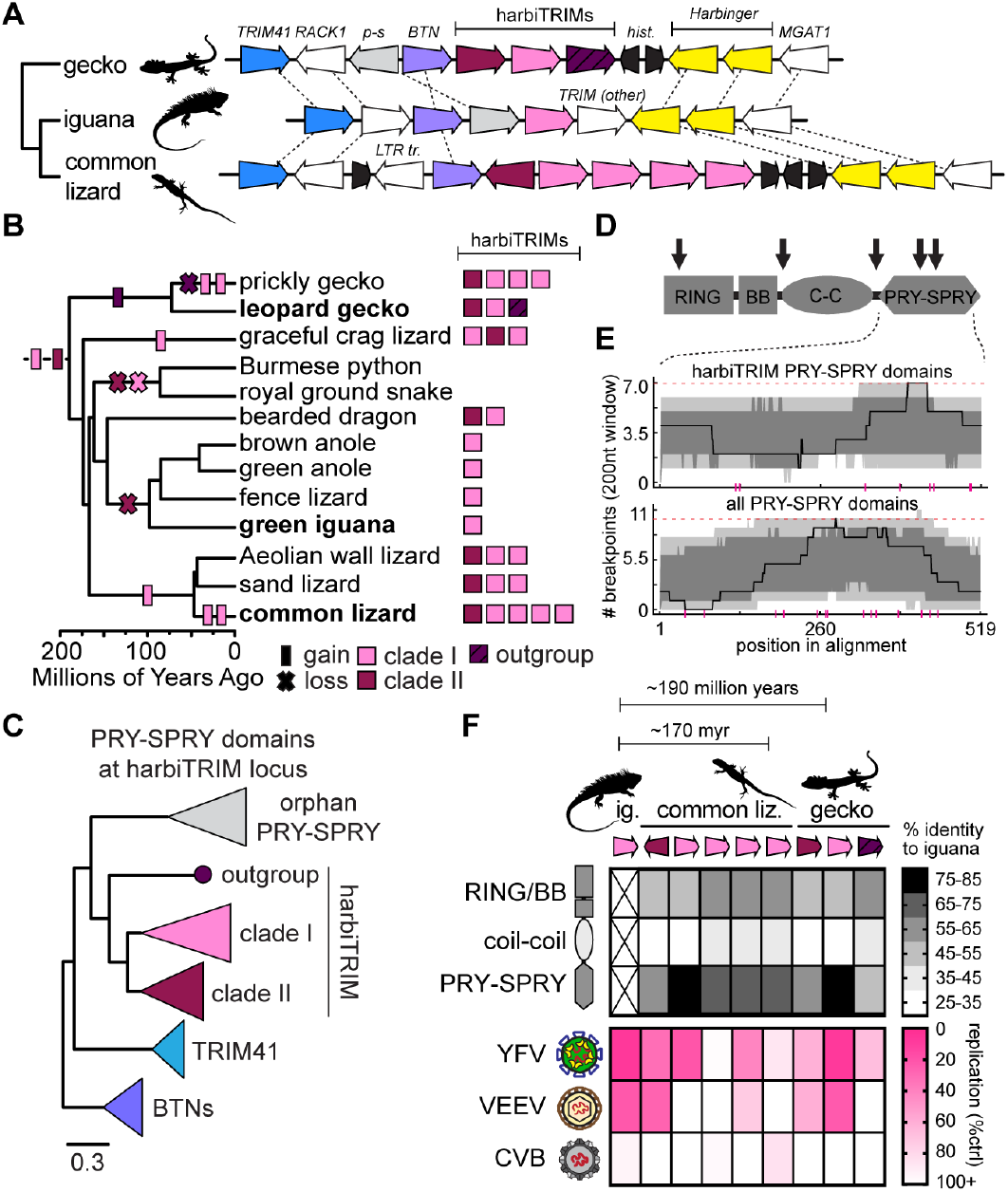
An expanded family of antiviral E3 ligases in squamate reptiles. (**A**) Schematic of the harbiTRIM locus in three squamate genomes. Butyrophilins (BTN), orphan PRY-SPRY domains (p-s), TRIMs, and other genes at the loci are indicated. (**B**) Summary of gains and losses of harbiTRIMs in squamates. **(C**) Maximum likelihood tree of PRY-SPRY domains from PRY-SPRY domain-containing proteins at the harbiTRIM locus. All bootstrap values for depicted nodes are > 88%. Scale bar: substitutions per residue. (**D**) Domain representation of iguana harbiTRIM with arrows indicating recombination breakpoints identified by GARD. (**E**) RDP4 analysis of recombination breakpoints and hotspots within PRY-SPRY domains of harbiTRIM homologs (top) and all PRY-SPRY domains at the harbiTRIM locus (bottom) in 13 squamate species. Light and dark grey shading indicate local 99% and 95% confidence intervals, respectively, and the dashed red line indicates the global 95% confidence threshold. Recombination breakpoints are indicated with purple tick marks. (**F**) Top heatmap: percent amino acid identity of indicated domains with iguana harbiTRIM. Bottom heatmap: Huh7 cells stably expressing the indicated harbiTRIM homologs were infected with the indicated viruses for 24 hours. Infection was quantified by luciferase assay (YFV) or GFP fluorescence (VEEV, CVB). Data plotted are relative to values obtained by infecting vector control-expressing cells.

Based on their PRY-SPRY domains, harbiTRIMs comprise three phylogenetically distinct clades, with clade I, to which iguana harbiTRIM belongs, being the most widespread (Fig. 3, A–C and fig. S3A). Clade I also exhibits the most genetic variability, with lineage- and species-specific expansions throughout squamates. The presence of tandem arrays of related TRIMs and proteins that contain similar domains prompted us to search for signatures of gene conversion. The Genetic Algorithm for Recombination Detection (GARD) (*16*), which detects alignment-wide recombination breakpoints, indicated five breakpoints, including two within the PRY-SPRY domain of harbiTRIMs (Fig. 3D). Further analysis by Recombination Detection Program 4 (RDP4) (*17*), which considers each possible triplet of sequences when inferring recombination events, found evidence of extensive recombination among PRY-SPRY domains present at this locus (Fig. 3E). Breakpoints are distributed throughout harbiTRIM and other PRY-SPRY domains, with only low-confidence “hotspots” of recombination, suggesting a history of widespread, recurrent recombination events (Fig. 3E). These patterns of extensive variation are consistent with a history of genetic conflict with viruses (*18*).

To test functional consequences of harbiTRIM variation, we synthesized all harbiTRIMs from common lizards (four from clade I, one from clade II) and leopard geckos (one each from clades I and II, and a highly diverged paralog). In experimental infections with the orthoflavivirus YFV, the alphavirus VEEV, and the picornavirus CVB, these TRIMs exhibit distinct patterns of antiviral activity (Fig. 3F and fig. S3, D and E). Some harbiTRIMs are similar to iguana harbiTRIM, inhibiting both orthoflaviviruses and alphaviruses, while others encode more specific activity. For example, the common lizard clade I harbiTRIM with the greatest amino acid identity to that of the iguana exhibits robust antiviral activity to YFV but, unlike the iguana harbiTRIM, does not inhibit VEEV. By contrast, the gecko ortholog of iguana harbiTRIM shared a similar profile of virus restriction, restricting both YFV and VEEV despite ∼200 million years of divergence. Other harbiTRIMs in the common lizard, however, inhibit VEEV and YFV to varying degrees, and both clade II harbiTRIMs from gecko and common lizard are of intermediate antiviral potency, inhibiting both YFV and VEEV. These observations are consistent with evolutionary transitions to distinct antiviral properties as TRIMs at this locus duplicated and diverged. These varied profiles also point to a vast and diversified genetic reservoir of antiviral activities for harbiTRIM variants restricting these and other viruses.

## Discussion

TRIM E3 ubiquitin ligases encoded by mammals are well-known restriction factors noted for recurrent bouts of rapid evolution (*19, 20*). Many studies on antiviral immunity in species diverged from mammals rely on detecting similar signals of rapid evolution to nominate TRIMs as antivirals (*21, 22*) and test candidates for inhibition of non-mammalian viruses. The discovery of harbingerTRIMs in squamates with a functional genetic screen reveals inhibition of orthoflaviruses and alphaviruses, showing that highly diverged immune gene repertoires can encode tools for combatting viruses relevant to human disease.

HarbingerTRIMs exhibit strong and distinct antiviral profiles, despite signals of gene conversion that may homogenize closely related PRY-SPRY domains (Fig. 3). Given high levels of recombination and the nearby enrichment of non-paralogous PRY-SPRY domains (fig. S3), putative antiviral TRIMs may sample diverged sequences, impacting the breadth of antiviral phenotypes. These exchanges could facilitate the spread of beneficial mutations arising within a single PRY-SPRY domain at the locus and/or quickly cull mutations incompatible with antiviral TRIM functions. These patterns differ from what has been observed for many other TRIM genes, which often exhibit signatures of diversifying, positive selection at individual residues that are important for antiviral specificity. Recurrent gene duplications and extensive genetic exchange offer a valuable comparison point for contrasting observations of other immune-involved TRIM loci, such as the young (∼100 myr old) TRIM6/34/5/22 cluster in placental mammals (*19*) and the more ancient (∼450 myr old) collection of TRIM genes situated near the major histocompatibility complex (MHC) in vertebrates (*23*).

The harbingerTRIM locus is of intermediate age (∼200 myr old) and exhibits substantial genetic volatility. This locus is also split across two separate scaffolds in the genome of a critical outgroup species to squamates, the Tuatara (*24*), where its evolutionary origins and linkage to the Harbinger transposon remain unresolved. Immunoregulatory TRIMs at the MHC locus generally exhibit lower sequence identity than harbiTRIMs and limited evidence of recombination, consistent with a history of ancient duplications and independent evolution (*23, 25*). TRIMs at the mammalian TRIM6/34/5/22 locus have undergone recent lineage-specific duplications and seem to evolve with lower levels of recombination in most lineages, but instead with enriched signals of pervasive diversifying selection (*19*). In contrast, an in-depth study (*26*) of the TRIM6/34/5/22 locus in rodents reported evolutionary patterns consistent with ongoing gene duplications and possible gene conversion. Given these findings and our observations with the harbingerTRIM locus, gene conversion and recombination in TRIM clusters may be a more impactful process in E3 ligase diversification than generally appreciated. These separate studies highlight the evolutionary flexibility of TRIMs in antiviral immunity, with a full complement of gene duplication, gene conversion, and diversifying, positive selection observed at different loci.

HarbiTRIM represents a distinct example of an E3 ligase restricting orthoflaviviruses by targeting capsid. A canonical example of a capsid-directed antiviral TRIM is TRIM5alpha, which recognizes HIV capsid (*27*), as does TRIMCyp, a unique TRIM-cyclophilin fusion protein (*28*). Our results suggest that harbingerTRIM promotes a ubiquitin-containing complex, without direct evidence that capsid itself is ubiquitylated (fig. S2, E and F). Similarly, TRIM5alpha does not measurably ubiquitylate HIV capsid, but instead undergoes autoubiquitylation as part of its antiviral mechanism (*29*). However, TRIM5alpha also forms an intricate lattice cage surrounding HIV capsid that disrupts viral uncoating (*30*), and whether harbiTRIM shares any aspect of this mechanism remains unknown. Other E3 ligases, such as TRIM41, recognize the nucleoproteins of viruses (*31, 32*). Together these observations reveal a growing collection of antiviral mechanisms targeting viral proteins that coat and protect viral nucleic acids.

Orthoflavivirus capsids require a form of non-degradative ubiquitylation for productive uncoating and genome release during infection (*9, 10*). A previous study showed that the loss of lysine residues in the capsid protein of dengue virus reduced infectivity in human cells (*9*). While it does not lead to attenuation in our experimental systems, a harbiTRIM resistance mutation for yellow fever virus involves loss of a capsid lysine residue (Figure 2). Further studies, particularly in vivo pathogenesis assays comparing replication of harbiTRIM-adapted and wild-type virus in mammal models, would be necessary to determine whether this presents an evolutionary constraint for cross-species adaptation of flaviviruses. Such a constraint might hinder orthoflaviviruses from adapting to infect squamate populations in the wild, as mechanisms to resist harbiTRIM may result in reduced infectivity in mammal hosts. The discovery of natural squamate-infecting orthoflaviviruses would provide a tool for clarifying this potential block to virus adaptation. A recent study surveying Australian skinks reported the presence of hepaciviruses (*33*), which are related to orthoflaviviruses, although it is not yet known whether hepaciviruses require ubiquitylation to facilitate genome uncoating.

The discovery of harbiTRIM antivirals in iguanas and other reptiles provides an accessible path to new antiviral discovery, even for species with limited experimental tools. Our study also highlights the impact of countless viral encounters with genetically diverse hosts over millions of years and the underlying links to viruses that infect humans today. Extending experimental biology into new species is critical given accelerating threats to biodiversity. For the field of infection biology, such an expansion will yield a deeper understanding of host-virus evolution and provide new opportunities for antiviral discovery.

## Supporting information

Supplemental Sequences Accesions

Supplemental RNAseq Data

Supplemental Alignments Trees

Supplemental Oligos

Supplemental Screen Results

Supplemental Genotyping Analysis

## Acknowledgments

We thank members of the Elde lab for helpful discussions. We thank Chris Faulk (University of Minnesota) for helpful discussions regarding genome assembly from long-read sequencing data. We additionally thank Jonathan Pruneda (OHSU) for helpful discussions concerning TRIM biology. Many of the computational analyses in this manuscript were performed with resources of the University of Utah Center for High Performance Computing (CHPC). We additionally thank Nitin Phadnis, Jonathan Pruneda, Mia Levine, Owen Pornillos, Lews Caro, and Patrick Mitchell for critical manuscript feedback.

## Author contributions

Conceptualization: I.N.B. and N.C.E.

Resources: I.N.B. and M.R.Q.

Investigation: I.N.B. and M.R.Q.

Visualization: I.N.B.

Funding acquisition: I.N.B. and N.C.E.

Supervision: N.C.E.

Writing – original draft: I.N.B.

Writing – review & editing: I.N.B. and N.C.E.

## Data and materials availability

Plasmids encoding including YFV-17D and VEEV-TC83 were obtained under MTA from Charles Rice (The Rockefeller University) and Bill Klimstra (University of Pittsburgh), respectively. RNA sequencing data have been deposited with the SRA (accession PRJNA1263791). The *Iguana iguana* genome assembly has been deposited with the NCBI (GCA_051529865.1). All data are available in the main text or the supplementary material.

## Supplementary Materials

Materials and Methods

Figs. S1 to S3

Tables S1 to S3

References (34-50)

Data S1 to S7

