## Supplemental Materials for "A trove of antiviral TRIM family E3 ligases in reptiles"

**The PDF file includes:**

Materials and Methods
Figs. S1 to S3
Tables S1 to S3
References (34-50)

**Other Supplementary Materials for this manuscript include the following:**

Data S1 to S7

### Materials and Methods

#### Cell culture

BHK-21J (ATCC), IGH2 (ATCC), Huh7 (a kind gift from Volker Thiel, University of Bern), HeLa (a kind gift from Adam Geballe, Fred Hutchinson Cancer Research Center), and 293T (a kind gift from Wes Sundquist, University of Utah) were cultured in DMEM supplemented with 10% FBS. Transduced cells were maintained in the presence of 4µg/mL puromycin. All cells were cultured at 37°C in 5% $CO_2$. All cells were cryopreserved in liquid phase liquid nitrogen.

#### Viruses

The infectious clone pACNR-FLYF-17Dx (kindly provided by Charles Rice, The Rockefeller University) was used to generate non-reporter YFV-17D. The infectious clone pWNV-MAD-IC (ATCC NR-49446) was used to generate non-reporter WNV. The infectious clone pVEEV-TC-83-Cap-eGFP-Tav (kindly provided by Bill Klimstra, University of Pittsburgh) was used to generate GFP reporter VEEV. The infectious clone pACNR-FLYF-17Dx-ΔC11-26-HiBiT (this study) was used to produce HiBiT-tagged YFV-17D. The infectious clone pCMV-DV2 (a kind gift from Matt Evans, Mount Sinai) was used to produce non-reporter DENV. The infectious clone pCMV-CVB-GFP (kindly provided by J. Lindsay Whitton, Scripps) was used to produce GFP reporter CVB3 (Woodruff strain).

For YFV, WNV, and VEEV, *in vitro* transcribed RNA was introduced into BHK-21J cells by electroporation (1µg RNA/$1\times10^6$ cells) and supernatants were collected once CPE was visible (typically 3 days for YFV and WNV and 2 days for VEEV), cleared by centrifugation at 2,000 xg for 10 minutes at 4°C, aliquotted, and stored at -80°C prior to use. CVB3 plasmid was introduced into HeLa cells by transfection (10µg plasmid/$4\times10^6$ cells). After one day, when cells were ~80% GFP-positive, cells were placed through two freeze-thaws, after which lysates were cleared and aliquotted as above. Dengue virus serotype 2 (DENV, DV2) was produced by transfecting 293T cells with plasmid and harvesting supernatant when cell death was apparent. This initial stock was amplified by one low-MOI passage in Huh7 cells, with supernatants from days 4 to 6 collected, cleared, aliquoted, and stored at -80°C prior to use. Viral stocks were re-titered annually to ensure stability. Viruses used in direct comparisons, such as recombinant viruses bearing mutations, were titered side-by-side in triplicate.

#### Lentiviral pseudoparticle production and transductions

All lentiviral pseudoparticles were generated by co-transfecting sub-confluent 293T cells with expression plasmids pSCRPSY (a kind gift of Paul Bieniasz, The Rockefeller University), pGag-Pol (J. Schoggins) and pVSV-glycoprotein (J. Schoggins) at a ratio of 25:5:1 using Lipofectamine 3000 (ThermoFisher). Two to six hours post-transfection, media was replaced with DMEM containing 3% FBS. Supernatants were collected at 48h, cleared by centrifugation, supplemented with 20mM HEPES, aliquotted, and stored at −80°C.

Cells were transduced by passive infection. Briefly, lentivirus was added to a minimum volume of transduction media (3% FBS, DMEM, 4µg/mL polybrene, 20mM HEPES) and added to cells.

Cells were allowed to rest with pseudoparticle-containing media for 1–2 hours before addition of complete medium. Cells were placed under puromycin selection 48 hours post-transduction.

Viral infections

Cells were seeded at 50,000–100,000 (24w plate, chambered cover glass for imaging),150,000–200,000 (12w plate), or 12,000,000 (15cm dish) cells per well, depending upon experiment endpoint, the day prior to infection. Virus was added to cells in a minimal volume of low-serum media and incubated for one hour (all viruses besides DENV) or two hours (DENV). After incubation, complete media was added to maintain cells until harvest. For infections to assess viral production by plaque assay or TCID50, inoculum was aspirated and cells were washed three times with PBS prior to addition of complete media. All multiplicities of infection (MOIs) are reported as plaque forming units (PFUs) (YFV-17D, DENV, WNV) or focus-forming units (FFUs) (CVB, VEEV, YFV-17D-HiBiT) per cell as determined at the time of plating.

For synchronized ("cold bind") experiments, media was changed to cold 1% FBS media and cells were chilled on ice in a cold room for 15–30 minutes prior to infection. Pre-chilled inoculum was added, and cells were incubated on ice in a cold room with intermittent rocking to allow virus to bind for one hour. Following incubation, inoculum was aspirated, cells were washed 3× with PBS on ice, and pre-warmed media was added to shift cells to 37°C to promote viral entry.

Electroporation assays

To directly introduce replication-competent viral RNA into cells, Huh7 cells were electroporated with either WT YF-17D RNA or YFV-17D-HiBIT RNA ($1\mu g/1\times10^6$ cells). Viral production was quantified by plaque assay or TCID50, and viral protein production was quantified by luciferase assay.

Generation of recombinant iguana interferon

A highly induced type I IFN transcript (see ORF sequence in (data S3)) was cloned from our cDNA library by PCR into a lentiviral vector. 293T cells were subsequently transduced and, following stable selection, conditioned media was obtained, cleared by centrifugation at 2,000 xg for 10 minutes at 4°C, aliquotted, and stored at -80°C prior to use. IFN-containing supernatant was tested for specificity and potency on iguana cells and human cells (293Ts, on which there was no transcriptional response for the ISG MX1) by qPCR, relative to control supernatant.

RT-qPCR

For gene expression assays, one-step reverse transcriptase pPCR was performed using the SuperScript III Platinum SYBR One-Step qPCR Kit (Invitrogen), with 50ng of RNA per reaction on a QuantStudio 3 Thermal Cycler.

Co-immunoprecipitations

Cells were washed with PBS and lysed on ice in RSB100T (50mM TRIS-HCl 7.5, 100mM NaCl, 2.5mM $MgCl_2$, 1mM $CaCl_2$, 1% Triton-X100, 1x cOmplete protease inhibitor cocktail (Roche)) for 15–20 minutes, after which nuclei were pelleted by a rapid spin at ~20,000×g. Lysates were subsequently incubated with beads for one hour, after which beads were captured on a magnetic separator and washed three times in RSB100T. Protein was eluted by boiling in 1x SDS buffer.

<u>Co-transfection assays</u>

For capsid degradation and co-immunoprecipitation assays, 293T cells were transfected with a 9:1 ratio of plasmids encoding harbiTRIM and flavivirus capsid, respectively, using Lipofectamine 3000. After 24 hours, cells were lysed in Laemmli buffer for western blotting. For co-immunoprecipitations, MG132 (1µM) was added two hours prior to harvest to enhance signal.

<u>Plaque assays</u>

For YFV, WNV, and DENV plaque assays, BHK-21J cells were seeded at 400,000 cells per well on 6 well plates one day prior to infection. Supernatants were serially diluted in DMEM supplemented with 1% FBS and 250ul of relevant dilutions were applied to BHKs. Cells were incubated with intermittent rocking for one hour, after which wells were overlaid with overlay media (1% Avicel, DMEM, 4% FBS, 100U/mL penicillin/100mg/mL streptomycin, 10mM HEPES, 0.1%$NaHCO_3$). After three (YFV, WNV) or five (DENV) days, wells were fixed with formaldehyde and plaques were visualized by staining with crystal violet.

<u>TCID50 assays</u>

Cells (for VEEV, YFV-17D-HiBiT: BHK-21J, for CVB: HeLa) were seeded at 20,000 cells per well on 96-well plates. Viral stocks were serially diluted in DMEM supplemented with 1% FBS, and cells were infected with 50µl of inoculum for 2 hours, after which 100µl of DMEM supplemented with 10% FBS was added per well. Wells were scored for GFP positivity three days post-infection. YFV-17D-HiBiT TCID50s were assayed by luciferase assay. TCID50 values were calculated by the method of Spearman and Kärber.

<u>Serial passaging and viral sequencing</u>

For the initial passage, Huh7 cells stably expressing harbiTRIM or a vector control were seeded at 250,000 cells per well on a 6w plate and infected with YFV-17D at an MOI of 100 for 48 hours. For subsequent passages, 500µL of virus-containing supernatant was directly transferred to freshly plated cells, and the remaining 1.5mL of supernatant was aliquot and stored at -80°C for titering and amplification. Cell monolayers were harvested with Trizol and banked at -80°C for sequencing. Working escape mutant stocks were amplified by infecting BHK cells at an MOI of 1 PFU per cell and harvested following one life cycle (24h) to minimize chances of accumulating additional mutations or maintaining bystander genomes by trans-complementation.

To identify point mutations, RNA was isolated from cells infected with passage five virus. RNA was reverse-transcribed using SuperScript III (Invitrogen) per manufacturer protocol using random hexamers. 18 tiling PCRs spanning the YFV genome (data S4) were performed using this cDNA as a template and PCR products were sequenced by Nanopore sequencing (Oxford Nanopore). Mutations were identified by comparing sequencing data to a reference genome. Genotyping was performed using raw fastq data using the “grep” command in bash to search for the sequences identified in the resistance mutants (see (data S6) for additional details).

Luciferase assays

To detect luciferase-tagged YFV-17D, the Nano-Glo® HiBiT Lytic Detection System (Promega) was used per manufacturer protocol and luminescence was read using a BioTek Synergy HT plate reader. We determined that signal hit maximum intensity for HiBiT-tagged capsid protein after 30 minutes of incubation, with five minutes of shaking on a plate shaker to assist in cell lysis. All luciferase values were determined relative to a luciferase-minus set of samples to subtract background within each experiment.

GFP fluorescence quantification

A BioTek H1 plate reader was used to quantify total-well GFP fluorescence for VEEV and CVB, relative to background fluorescence as determined per cell line alongside each experiment to control for potential autofluorescence and bleed-through from tagRFP. Clear bottom, opaque-walled plates were used for all GFP experiments.

Nanopore sequencing

For genome sequencing, high molecular weight DNA was obtained from low-passage IGH2 cells using a NanoBind CBB kit (PacBio). Sequencing libraries were prepped using the Ligation Sequencing Kit V14 (Oxford Nanopore) and sequenced using a MinIon Mk1C (Oxford Nanopore).

For transcriptome sequencing, the Direct cDNA Sequencing V14 kit (Oxford Nanopore) was used per manufacturer recommendations, with total RNA as input. Libraries were sequenced using a MinIon Mk1C (Oxford Nanopore).

Genome assembly

We assembled the *Iguana iguana* genome by broadly following the method published by Christopher Faulk (*34*). First, raw nanopore reads were re-basecalled using Dorado (Oxford Nanopore). Short reads (reads below 1,500bp) were removed using samtools (*35*). An initial assembly was generated using Flye (v2.9.3) (*36*) (mode: nano-hq, two iterations). Medaka (Oxford Nanopore) (v1.11.3) was used to polish the genome (41x coverage, BUSCO of 95.2%). A nextdenovo assembly was also generated and found to be of lower quality (24x coverage, BUSCO of 82.5%) than that of the flye/medaka pipeline. BUSCO (v5.3.2) (*37*) was used to determine assembly quality. The genome was scaffolded to the *Cyclura pinguis* genome (rCycPin1), which diverged from the iguana approximately 36 million years ago, using ragtag (*38*). Repeats were identified de novo using RepeatModeler (*39*) from RepeatMasker using default settings. Quast (*40*) was used to calculate assembly statistics.

Transcriptome assembly

RNA-Bloom (*41*) (v2.0.1) was used to generate a transcriptome using a total of 14 million ONT reads, with approximately half from untreated and half from poly(I:C)-treated IGH2 cells. For differential gene expression analysis, "SuperTranscripts" (data S7) were generated to avoid multi-mapping issues from splice variants using the pipeline published by Davidson, Hawkins, and Oshlack (*42*). Briefly, Corset was used to cluster transcripts which were then used by Lace to generate SuperTranscripts for each cluster.

Illumina RNA sequencing

Illumina RNA sequencing was performed by Azenta. For the cDNA library screen, RNA-sequencing was performed at ~50 million reads per sample with 2x150bp reads. For the iguana IFN RNA seq experiment, a read depth of ~20 million per sample was used.

cDNA library construction

IGH2 cells were seeded at 7.25E6 cells per p150 across 7 dishes. The following day, cells were transfected with high molecular weight poly(I:C) (InvivoGen) (35μg per dish using FuGene HD at a 4:1 ratio). After 6 hours, RNA was isolated with TRIzol. mRNA was purified with a PolyA Purist mRNA purification kit (ThermoFisher). The CloneMiner II cDNA library construction kit (ThermoFisher) was used to generate the Gateway-compatible cDNA library from 1μg of mRNA. Library size was calculated by plating dilutions of library-containing bacteria on agarose plates for CFU counts. The library was qualified by restriction digest to ensure appropriate average insert size. The cDNA library was transferred into an expression library (pSCRPSY) by LR reaction. Libraries were amplified using SeaPrep soft agar (Lonza) per manufacturer recommendations. Briefly, 2X LB/SeaPrep agar was autoclaved and chilled to 25°C in a water bath, after which antibiotics and library-transformed bacteria were added. Inoculated LB was aliquotted into 50mL conical vials and submerged in wet ice for one hour, after which conicals were incubated at 37C for 48h. Conicals were centrifuged, agar was decanted, and pellets were pooled and midiprepped (Zymo).

Restriction factor screen

Huh7 cells were transduced at an MOI of 2 with lentiviral pseudoparticles containing the iguana cDNA library. The library was introduced at approximately 10,000x transcriptome coverage and cells were expanded prior to infection. Library-transduced Huh7 cells (at a transcriptome coverage of 350x per replicate) were infected with DENV at an MOI of 0.02. RNA was harvested from surviving cells at 12 days. RNA from surviving cells was compared to RNA from uninfected, library-transduced cells (150x coverage per replicate) to identify enriched cDNAs. Screens were performed in biological triplicate. RNA-sequencing was performed at ~50 million reads per sample with 2x150bp reads by Azenta. Salmon (*43*) was used to map and quantify Illumina reads to our *Iguana iguana* SuperTranscript assembly. Differential expression analysis was conducted using DESeq2 (*44*) in an R (v 4.2.2) environment in R Studio (v 2022.12.0.0+353).

Annotation of iguana ISGs

NCBI BLAST as implemented on the University of Utah CHPC was used to blast all SuperTranscripts by blastn and dc-megablast. Transcripts with either poor E values or alignment scores were subsequently manually checked by blastx. ISGs were identified as being shared with mammals if they were shown to be upregulated in more than one mammalian species in the Orthologous Clusters of Interferon-Stimulated Genes database (*45*).

Phylogenetic analyses

Multiple sequence alignments were generated using Muscle as implemented in AliView (*46*) and manually inspected for likely sequencing errors. Phylogeny was inferred using maximum likelihood trees generated by IQ-TREE 2 (*47*). TimeTree 5 (*48*) was used to infer species trees. Trees and alignments are available in (data S5).

Synteny analysis

The genomes listed in (table S3) were manually inspected in the NCBI genome viewer to assess synteny at the harbiTRIM locus. harbiTRIM pry-spry domains were used to determine which harbiTRIM cluster particular homologs belonged to by generating maximum likelihood trees using IQ-TREE 2(*47*). For the *Iguana iguana* locus, RNA-seq reads (PRJNA490698, SRR1693196, and those generated in the present study) were mapped to our draft genome assembly using STAR (*49*) to identify expressed genes, which were then identified and characterized by BLAST.

Immunofluorescence

Cells were fixed with 4% PFA in PBS. Cells were washed with PBS, then permeabilized with 0.2% Triton X-100 in PBS. Cells were blocked with 5% goat serum in PBS for at least 30 minutes. Primary antibody was added in blocking solution and incubated for 1–2 hours. Cells were washed 3× with PBS, after which secondary antibody was added in PBS containing 3% BSA and incubated for 30 minutes. Cells were washed 3× with PBS and then mounted using Fluormount-G (Southern Biotech). Imaging was performed on a Zeiss LSM980 laser-scanning confocal microscope. Images were processed in ImageJ. When made, linear adjustments were applied evenly across all samples within each experiment. Primary antibodies used were: anti-FLAG (F1804) and anti-dsRNA (J2; Thermo 10010500). Secondary antibodies used were: anti-Ms-647 (A-21236), anti-Ms-647 (A-21244), anti-Ms-488 (A-11001), and anti-Rb-488 (A-32731).

Western blotting

Unless otherwise noted, cells were lysed directly in 1× SDS loading buffer (10% glycerol, 5% BME, 62.5mM TRIS-HCl pH 6.8, 2% SDS, and BPB), boiled, and sonicated (Qsonica Q500). Samples were run on “Any kD” acrylamide gels (Bio-Rad) and transferred to PVDF membranes using a Trans-Blot Turbo (Bio-Rad). Blots were blocked in 5% dry milk/TBS-T for 30 minutes to an hour at RT. Primary antibodies were diluted in 5% dry milk/TBS-T and added for 1 to 2 hours at RT or overnight at 4°C. Blots were washed four times in TBS-T before addition of HRP-conjugated secondary antibody in 5% milk for thirty minutes. Blots were washed four times in TBS-T prior to detection with Clarity ECL (Bio-Rad) substrate and imaging on an Azure Q500 (Azure). For some experiments, infrared secondaries (Azure spectra dyes) were used. Primary antibodies used were: Rb-anti-YFV-C (GTX134022), Rb-anti-DV2-C (GTX639659) Ms-anti-HiBiT (Promega N720A), Rb-anti-STAT1 (CST 9172S), Ms-anti-ACTB (Sigma A1978), Ms-anti-FLAG (Sigma F1804), Rb-anti-FLAG (Sigma F7425), Rb-anti-MYC (Sigma SAB4300319), Ms-anti-HA (Thermo 5B1310), and a Ms IgG isotype control (Thermo 31903). Secondary antibodies used were: anti-Ms (Sigma AP130P) and anti-Rb (Sigma AP132P).

Cloning

Primers used for cloning are listed in (data s4).

The following were cloned from the *Iguana iguana* cDNA library by PCR, digest, and ligation into pSCRPSY-MCS: pSCRPSY_harbiTRIM, pSCRPSY_igigIFNkappa.

pSCRPSY_harbiTRIM was used as a template to clone C-terminal 1x FLAG and 3x FLAG-tagged constructs by PCR. The 3x FLAG-tagged harbiTRIM was used to clone the harbiTRIM_2S loss of function mutant.

YFV and DENV capsids were cloned from infectious clone plasmid by PCR into the pEF backbone. N-terminal myc tags were added by PCR.

The following harbiTRIM homologs were codon-optimized and synthesized by Genscript: XP_054832434.1, XP_054832433.1, XP_034962875.1, XP_060126838.1, XP_060126836.1, XP_034962878.2, XP_054834655.1, XP_034960918.2.

pACNR-FLYF-17Dx_dC11-26-HiBiT was constructed by Gibson assembly. A nonessential alpha-helix was selected for tag insertion based on a previous study (*50*).

HarbiTRIM resistant virus was engineered into the pACNR-FLYF-17Dx and pACNR-FLYF-17Dx_dC11-26-HiBiT backgrounds by Gibson assembly.

CRISPR of IGH2 cells

Pre-complexed GFP-tagged Cas9 (Genscript Z03393) and pooled CRISPR guide RNAs (50pmol Cas9:100pmol RNA oligos:1×$10^6$ cells) were electroporated using a Neon electroporator (Thermo) (1.2kV, 20μm width, 3x pulses) and placed in complete media. The next day, GFP+ cells (~3,000 cells per condition) were sorted and collected in a single well on a 48w plate. Once cells were confluent, an aliquot was taken for clonal derivation by limiting dilution while the remainder was maintained as a bulk population. Guide RNA sequences are listed in (data S4). Clones were genotyped by PCR using the primers included in (data S4) with ICE (Synthego) and CRISPRLungo (data S6).

Quantification and statistical analysis

Unless otherwise indicated, individual data points represent independent biological replicates. Bars and lines with error bars indicate the mean ± SD. Experiments were performed with three independent biological replicates unless otherwise noted. Statistical analyses were performed using GraphPad Prism unless otherwise noted. Unless otherwise indicated, all comparisons are relative to control (ctrl), as labeled. For data with two groups, two-tailed t tests were used. For data with more than two groups, ANOVA were used, and appropriate adjustments were made for multiple hypothesis testing (Šídák's multiple comparisons test for most data, Fisher's LSD test for multiple pairwise comparisons in Fig. 1F). All luciferase and fluorometry data are background-adjusted to blank wells. Unless otherwise specified, P values are denoted as follows: n.s. not significant, *P<0.05, **P<0.01, ***P<0.001, ****P<0.0001.

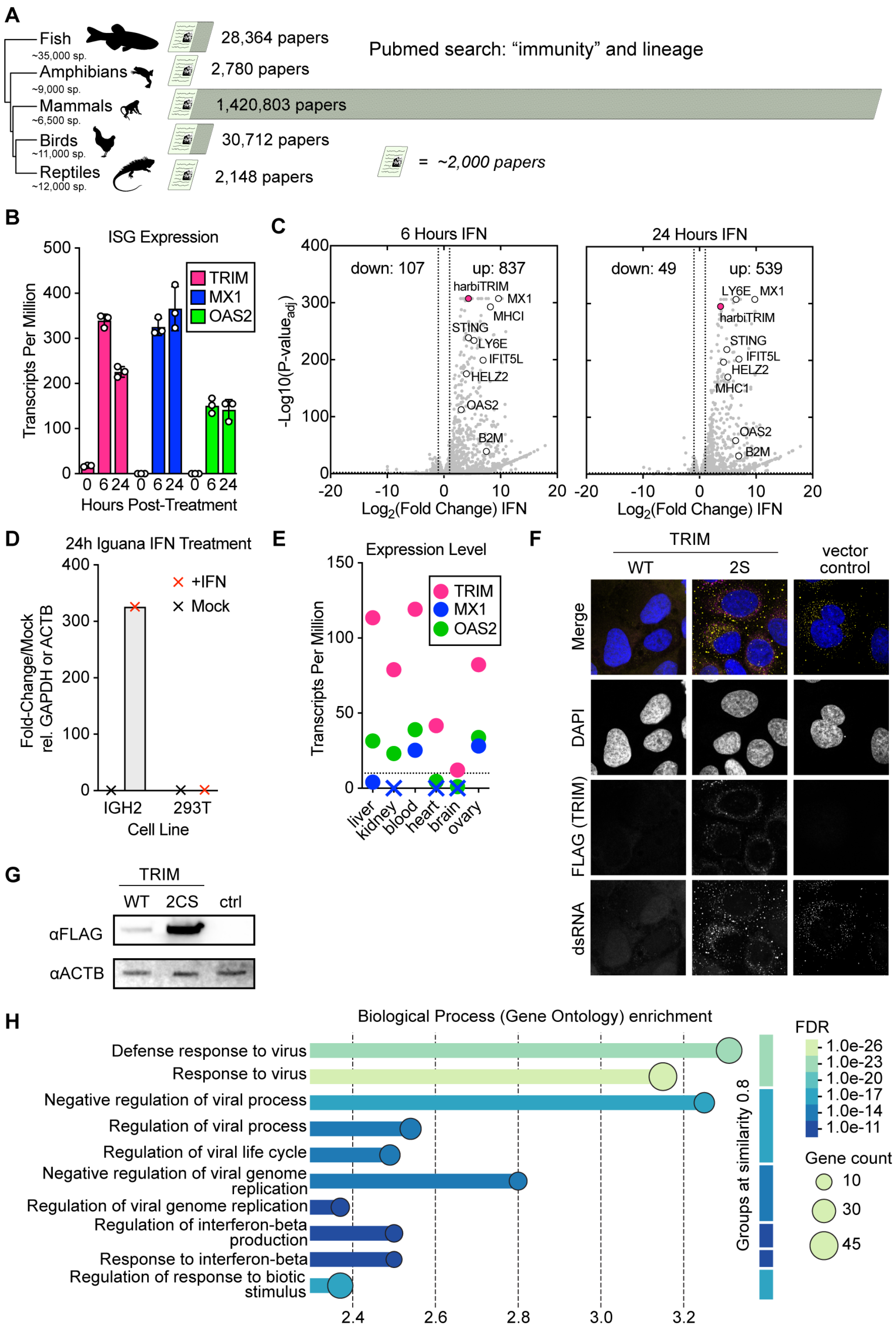


**fig. S1.**

(**A**) PubMed search results for “immunity” and either fish, amphibians, mammals, birds, or reptiles on 5/12/2025. (**B**) Transcripts per million from RNA seq data from IGH2 cells treated with IFN for 0, 6, and 24 hours. harbiTRIM and two canonical ISGs are shown. (**C**) IGH2 cells were treated with recombinant iguana IFN for 6 or 24 hours and transcripts were quantified by RNA sequencing. Select upregulated genes are indicated. (**D**) IGH2 and 293T cells were treated with iguana IFN containing supernatant at a 1:100 dilution for 24 hours. qPCR was used to assess MX1 induction in both species backgrounds. (**E**) Publicly available RNA seq data (PRJNA490698, SRR1693196) were mapped to the iguana transcriptome to enable tissue-specific quantification of harbiTRIM and select ISGs. X: not detected. Dotted line: TPM of 10, indicative of low expression. (**F**) Cropped section of Figure 1H, with FLAG staining included. (**G**) Western blot of harbiTRIM-3xFLAG and harbiTRIM-2S-3xFLAG expressing stable cell lines demonstrating relative TRIM expression levels. (**H**) 342 non-redundant significantly upregulated transcripts were imported into the STRING database for gene ontology analysis.

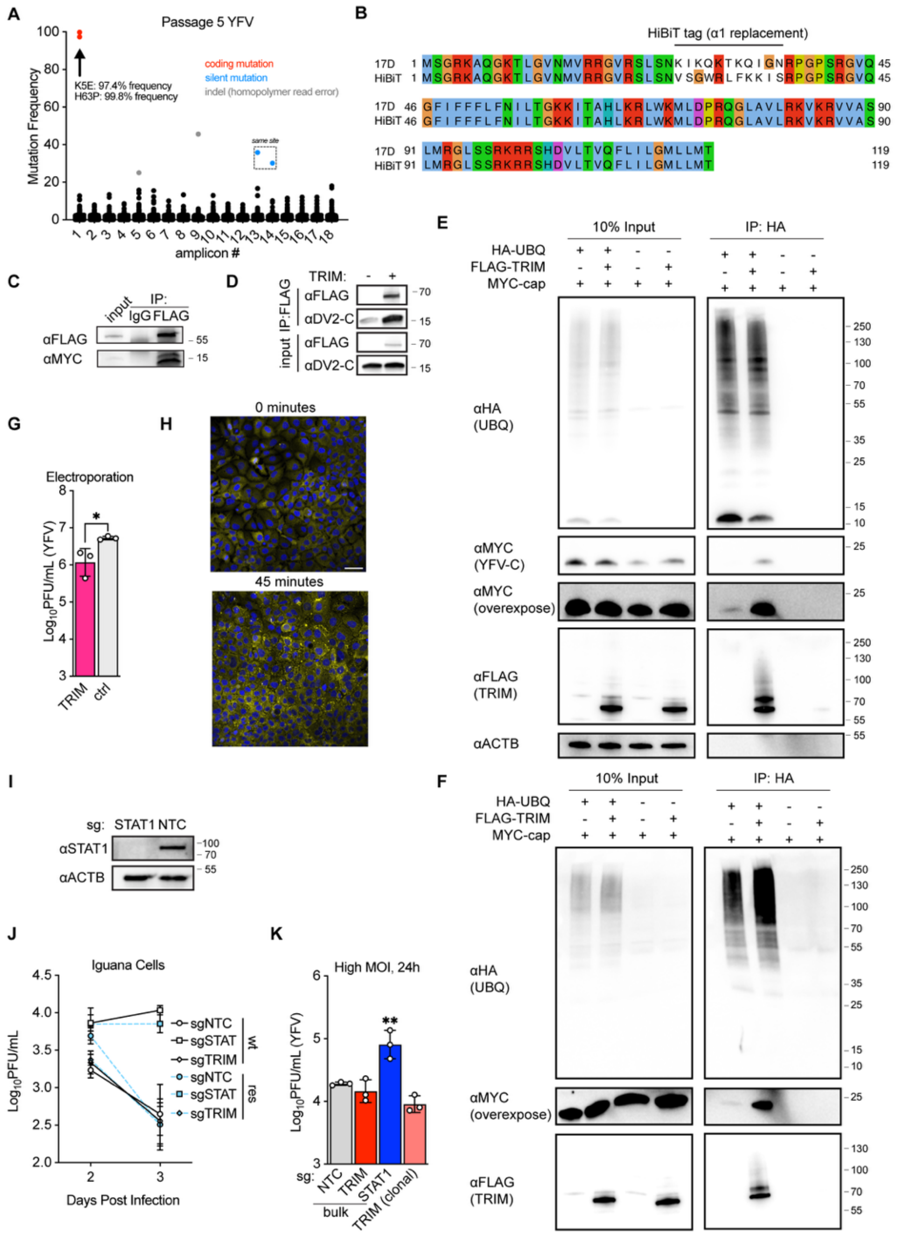

**fig. S2.**

(**A**) Amplicon sequencing results of passage five harbiTRIM resistant virus. (**B**) Alignment of YFV-17D capsid and YFV-17D-HiBiT capsid. (**C**) 293T cells were co-transfected with plasmids encoding yellow fever virus capsid and harbiTRIM. 24 hours post-transfection, cells were lysed and immunoprecipitated with either anti-FLAG antibody or an IgG control for western blotting. (**D**) 293T cells were co-transfected with plasmids encoding dengue capsid and either harbiTRIM or a vector control. 24 hours post-transfection, cells were lysed and FLAG-tagged harbiTRIM was immunoprecipitated for western blotting. (**E**) 293T cells were co-transfected with the indicated plasmids. 24 hours post-transfection, cells were lysed and HA-tagged ubiquitin was immunoprecipitated for western blotting. (**F**) 293T cells were co-transfected with the indicated plasmids. 24 hours post-transfection, cells were treated with MG132 for 4 hours, lysed in the presence of the deubiquitylase inhibitor N-Ethylmaleimide and HA-tagged ubiquitin was immunoprecipitated for western blotting. Lack of free ubiquitin (compared to fig. S2E) indicates successful deubiquitylase inhibition. (**G**) Huh7 cells expressing harbiTRIM or a vector control were electroporated with YFV-17D RNA and supernatants were collected 24 hours post electroporation. (**H**) Cells were synchronously infected with YFV-17D at an MOI of 50. Cells were fixed for immunofluorescence at the indicated time points post-shift to 37°C. Scale bar: 50 μm (**I**) Western blot demonstrating loss of STAT1 in STAT1-targeted IGH2 cells. (**J**) Bulk populations of CRISPR-targeted IGH2 cells were infected with YFV-17D at an MOI of 1 for 72 hours. 48h data are the same date represented in Figure 2M. (**K**) The indicated IGH2 cell lines were infected with YFV-17D at an MOI of 10 for 24h.

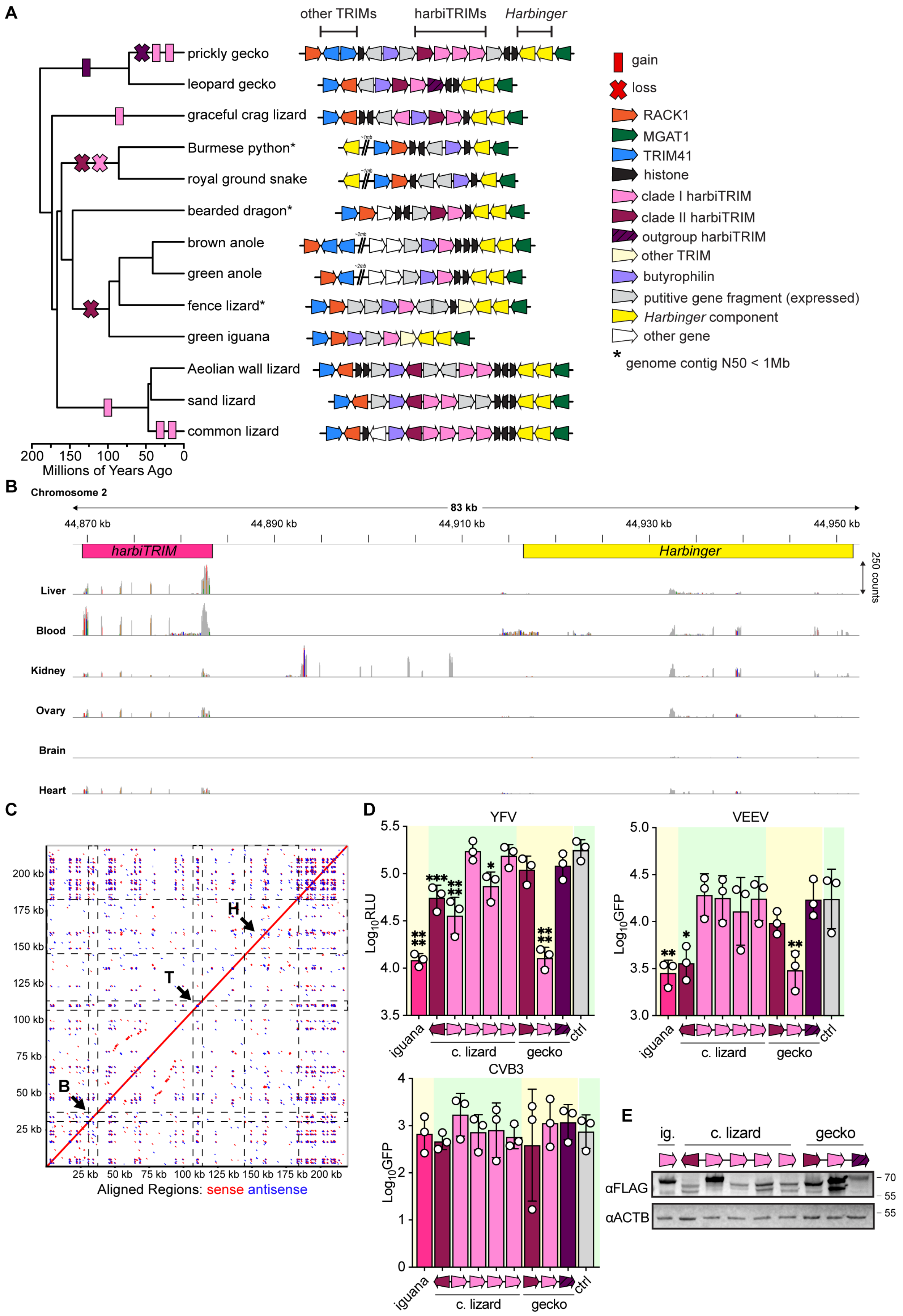
A
other TRIMs
harbiTRIMs
Harbinger
prickly gecko
leopard gecko
graceful crag lizard
Burmese python*
royal ground snake
bearded dragon*
brown anole
green anole
fence lizard*
green iguana
Aeolian wall lizard
sand lizard
common lizard
200 150 100 50 0
Millions of Years Ago
gain
loss
RACK1
MGAT1
TRIM41
histone
clade I harbiTRIM
clade II harbiTRIM
outgroup harbiTRIM
other TRIM
butyrophilin
putitive gene fragment (expressed)
Harbinger component
other gene
* genome contig N50 < 1Mb
B
Chromosome 2
83 kb
44,870 kb
44,890 kb
44,910 kb
44,930 kb
44,950 kb
harbiTRIM
Harbinger
250 counts
Liver
Blood
Kidney
Ovary
Brain
Heart
C
Aligned Regions: sense antisense
D
YFV
VEEV
CVB3
Log10RLU
Log10GFP
iguana
c. lizard
gecko
ctrl
E
ig.
c. lizard
gecko
αFLAG
αACTB
70
55
55

**fig. S3.**

(**A**) Synteny analysis of the harbiTRIM locus in 13 squamates. (**B**) IGV overview of RNA-seq reads mapping to the *harbiTRIM* locus in the green iguana. Peaks represent mapped reads, ranging from 0-250 reads per base on a linear scale. The two-gene *Harbinger* transposon and *harbiTRIM* are highlighted. This represents the only intact, transcriptionally active *Harbinger* transposon in the iguana genome. (**C**) DNA dot plot of the harbiTRIM locus in the common green iguana. Axes show position in nucleotide bases. Labels are B: butyrophilin, T: harbiTRIM, and H: *Harbinger* transposon, as labeled in Figure 3A and supplemental figure 3a. (**D**) Huh7 cells stably expressing the indicated harbiTRIM homologs were infected with the indicated viruses for 24 hours. Infection was quantified by luciferase assay (YFV) or GFP fluorescence (VEEV, CVB). All statistical comparisons are made relative to vector control. HarbiTRIM homologs are represented in the same order as in the native locus, as depicted in Figure 3A and supplemental figure 3a. (**E**) Alongside two screen replicates, counted cells were collected for western blotting. Representative blot of n = 2 biological replicates.

**table S1.**

| | |
|---|---|
| Primary clones (CFU) | $6.3x10^6$ |
| Estimated coverage | 350x |
| Average insert (n=24 tested) | 1.1kb |
| Maximum insert (n=24 tested) | 2.4kb |

cDNA library quality control metrics.

**table S2.**

| | |
|---|---|
| n. reads used in assembly | 22,625,639 |
| Average read quality | 28.4 |
| Longest read | 1,012,048 base pairs |
| Assembly size | 2.01 GB |
| Average coverage | 41x |
| Contig N50 | 12.74 MB |
| Contig L50 | 44 |
| Scaffold N50 | 294.5 MB |
| Scaffold L50 | 3 |
| BUSCO | 95.2% [S:93.7%,D:1.5%],F:1.0%,M:3.8% |
| Repeat content | 43.2% |

Genome assembly statistics.

**table S3.**

| Scientific name | Common name | Genome | Coverage | Sequencing platform | Genome Size (Gb) | Scaffold N50 (Mb) | Contig N50 (Mb) |
|---|---|---|---|---|---|---|---|
| Zootoca vivipara | common lizard | GCF 963506605.1 | 34x | PacBio,Arima2 | 1.4 | 96.1 | 3.4 |
| Heteronotia binoei | prickly gecko | GCF 032191835.1 | 21x | PacBio Sequel | 2.6 | 150.6 | 14.4 |
| Anolis sagrei | brown anole | GCF 037176765.1 | 34.9x | PacBio Sequel II HiFi; Arima Hi-C v2 | 2 | 249.2 | 43.1 |
| Eublepharis macularius | leopard gecko | GCF 028583425.1 | 30x | PacBio Sequel | 2.2 | 145.6 | 80.1 |
| Pogona vitticeps | central bearded dragon | GCF 900067755.1 | 85x | Unspecified shotgun WGS | 1.8 | 2.3 | 0.0333 |
| Sceloporus undulatus | fence lizard | GCF 019175285.1 | 4859x | Illumina HiSeq; PacBio | 1.9 | 275.6 | 0.0755 |
| Lacerta agilis | sand lizard | GCF 009819535.1 | 63.4x | PacBio Sequel I CLR; Illumina NovaSeq; Arima Hi-C; Bionano DLS | 1.4 | 86.6 | 6.6 |
| Podarcis raffonei | Aeolian wall lizard | GCF 027172205.1 | 31x | PacBio Sequel II HiFi; Bionano DLS; Arima Hi-C v2 | 1.5 | 93.6 | 61.4 |
| Anolis carolinensis | green anole | GCF 035594765.1 | 35x | PacBio Sequel | 1.9 | 289 | 26.1 |
| Hemicordylus capensis | graceful crag lizard | GCF 027244095.1 | 43x | PacBio Sequel IIe; Dovetail Omni-C | 2.3 | 359.6 | 160.7 |
| Iguana iguana | common green iguana | GCA 051529865.1 | 41x | Oxford Nanopore | 2 | 294.5 | 12.74 |
| Python bivittatus | Burmese python | GCF 000186305.1 | 20x | Illumina; 454 | 1.4 | 0.214 | 0.0107 |
| Erythrolam-prus reginae | royal ground snake | GCF 031021105.1 | 34.5x | PacBio Sequel II HiFi; Arima Hi-C v2 | 2 | 259.1 | 3.1 |

Genomes used for synteny analysis.

**Data S1. (separate file)**
Screen results

**Data S2. (separate file)**
Iguana interferon RNA sequencing results

**Data S3. (separate file)**
Accession numbers and sequences

**Data S4. (separate file)**
Oligos used in study

**Data S5. (separate file)**
Alignments and phylogenetic trees

**Data S6. (separate file)**
Iguana cell harbiTRIM CRISPR validation and yellow fever virus genotyping

**Data S7. (separate file)**
Iguana supertranscripts
