## Supplemental Alignments Trees for "A trove of antiviral TRIM family E3 ligases in reptiles"

**PRY-SPRY TREE**

#NEXUS

begin taxa;

dimensions ntax=54;

taxlabels

'aeolian_BTN_XP_053234972.1'

'aeolian_hTRIM_XP_053234980.1'

'aeolian_left1_XP_053234931.1'

'aeolian_left4_XP_053234941.1'

'aeolian_leftOfBTN_psOnly_XP_053235017.1'

'anole_BTN_XP_067319452.1'

'anole_hTRIM_XP_067319465.1'

'anole_thaicobrin_XP_060615369.2'

'beardie_hTRIM_XP_020633187.1'

'beardie_ligase_left_1_XP_020633194.1'

'beardie_thaicobrin_right_of_RACK1_XP_020633192.1'

'beardie_trim41_left_of_RACK1_XP_020633185.1'

'cliff_BTN_XP_053158942.1'

'cliff_hTRIM_XP_053151972.1'

'cliff_left1_XP_053143803.1'

'cliff_leftOfBTN_1_XP_053158941.1'

'cliff_left_of_BTN_2_psonly_XP_053149380.1'

'common_lizard_BTN_XP_060126844.1'

'common_lizard_hTRIM_XP_060126836.1'

'common_lizard_ligase_left1_XP_034960918.2'

'common_lizard_ligase_left2_XP_034962875.1'

'common_lizard_ligase_right1_XP_034962878.2'

'common_lizard_tdhTRIMdup_XP_060126838.1'

'fence_BTN_XP_042310934.1'

'fence_LeftOfHarbi1_XP_042310940.1'

'fence_LeftOfHarbi3_fragment_XP_042311000.1'

'fence_leftofBTN_1_psOnly_XP_042311001.1'

'fence_leftofBTN_2_psOnly_XP_042303361.1'

'fence_leftofHarbi4_XP_042310941.1'

'gecko_btn_XP_060095246.1'

'gecko_hTRIM_XP_060095237.1'

'gecko_ligase_T41_left_of_BTN_1_XP_060095233.1'

'gecko_ligase_T41_left_of_BTN_2_XP_060095232.1'

'gecko_ligase_left1_XP_060094749.1'

'gecko_ligase_right_1_XP_060095238.1'

'gecko_ligase_right_2_XP_060095236.1'

'gecko_thaicobrin_left_of_BTN_XP_060095244.1'

'greenAnole_BTN_XP_008103839.1'

'greenAnole_hTRIM_XP_008103837.1'

'greenAnole_leftofBTN_psOnly_XP_062825617.1'

iguana_BTN_pryspry_only

iguana_hTRIM

iguana_ligase_right1_pryspry_only

'lepgecko_BTN_XP_054832433.1'

'lepgecko_FusedRightOfBTN_XP_054832433.1'

'lepgecko_hTRIM_XP_054834655.1'

'lepgecko_psOnly_leftofBTN_XP_054832432.1'

'lepgecko_right1_XP_054832434.1'

'sand_BTN_XP_032997396.1'

'sand_hTRIM_XP_032997387.1'

'sand_left1_XP_032997388.1'

'sand_left2_XP_032997424.1'

'sand_leftofBTN_psOnly_XP_032997411.1'

'sand_trim41_XP_032997386.1'

;

end;

begin trees;

tree tree_1 = [&R] (iguana_ligase_right1_pryspry_only:0.141799,'fence_LeftOfHarbi1_XP_042310940.1':0.171895,((((((('aeolian_leftOfBTN_psOnly_XP_053235017.1':0.041054,'sand_leftofBTN_psOnly_XP_032997411.1':0.080592)[&label=100]:0.426816,('cliff_left_of_BTN_2_psonly_XP_053149380.1':0.182359,('lepgecko_psOnly_leftofBTN_XP_054832432.1':0.172368,'gecko_thaicobrin_left_of_BTN_XP_060095244.1':0.193341)[&label=100]:0.211258)[&label=89]:0.077525)[&label=68]:0.04481,'fence_leftofBTN_2_psOnly_XP_042303361.1':0.382191)[&label=60]:0.02823,((('anole_thaicobrin_XP_060615369.2':0.066543,'greenAnole_leftofBTN_psOnly_XP_062825617.1':0.085662)[&label=100]:0.193959,'fence_leftofBTN_1_psOnly_XP_042311001.1':0.232742)[&label=93]:0.047918,'beardie_thaicobrin_right_of_RACK1_XP_020633192.1':0.206497)[&label=96]:0.094588)[&label=100]:0.564092,('lepgecko_right1_XP_054832434.1':0.687296,(('cliff_leftOfBTN_1_XP_053158941.1':0.635104,(('beardie_hTRIM_XP_020633187.1':0.227716,(('fence_leftofHarbi4_XP_042310941.1':0.251245,('anole_hTRIM_XP_067319465.1':0.052979,'greenAnole_hTRIM_XP_008103837.1':0.017144)[&label=100]:0.085222)[&label=99]:0.049289,iguana_hTRIM:0.046372)[&label=100]:0.064513)[&label=94]:0.03687,((('gecko_hTRIM_XP_060095237.1':0.196546,('lepgecko_hTRIM_XP_054834655.1':0.066671,('gecko_ligase_right_1_XP_060095238.1':0.051874,'gecko_ligase_right_2_XP_060095236.1':0.035594)[&label=100]:0.101126)[&label=62]:0.018964)[&label=75]:0.049469,'cliff_hTRIM_XP_053151972.1':0.140491)[&label=70]:0.06253,((((('aeolian_hTRIM_XP_053234980.1':0.112729,('common_lizard_hTRIM_XP_060126836.1':0.007782,'common_lizard_ligase_right1_XP_034962878.2':0.021825)[&label=91]:0.01524)[&label=100]:0.093584,'common_lizard_tdhTRIMdup_XP_060126838.1':0.080332)[&label=93]:0.049164,'sand_hTRIM_XP_032997387.1':0.025332)[&label=74]:0.037889,'sand_left1_XP_032997388.1':0.022579)[&label=97]:0.04593,('common_lizard_ligase_left1_XP_034960918.2':0.04654,'aeolian_left1_XP_053234931.1':0.036934)[&label=68]:0.005896)[&label=100]:0.110221)[&label=63]:0.030889)[&label=98]:0.182045)[&label=86]:0.100667,(('lepgecko_FusedRightOfBTN_XP_054832433.1':0.172338,'gecko_ligase_left1_XP_060094749.1':0.178863)[&label=100]:0.146886,((('beardie_ligase_left_1_XP_020633194.1':0.220019,'fence_LeftOfHarbi3_fragment_XP_042311000.1':0.166547)[&label=93]:0.086843,('sand_left2_XP_032997424.1':0.043363,('common_lizard_ligase_left2_XP_034962875.1':0.022997,'aeolian_left4_XP_053234941.1':0.006756)[&label=98]:0.025081)[&label=100]:0.120388)[&label=100]:0.087232,'cliff_left1_XP_053143803.1':0.196703)[&label=97]:0.071582)[&label=95]:0.094495)[&label=97]:0.133662)[&label=89]:0.13388)[&label=95]:0.125373,((('gecko_ligase_T41_left_of_BTN_1_XP_060095233.1':0.085787,'gecko_ligase_T41_left_of_BTN_2_XP_060095232.1':0.002865)[&label=100]:0.1032,'sand_trim41_XP_032997386.1':0.036318)[&label=97]:0.047661,'beardie_trim41_left_of_RACK1_XP_020633185.1':0.0443)[&label=100]:0.574822)[&label=93]:0.082046,((((('common_lizard_BTN_XP_060126844.1':0.057908,'sand_BTN_XP_032997396.1':0.043538)[&label=92]:0.023925,'aeolian_BTN_XP_053234972.1':0.033419)[&label=100]:0.142646,('gecko_btn_XP_060095246.1':0.102981,'lepgecko_BTN_XP_054832433.1':0.053069)[&label=99]:0.036389)[&label=89]:0.032769,'cliff_BTN_XP_053158942.1':0.13734)[&label=67]:0.031882,((('anole_BTN_XP_067319452.1':0.076001,'greenAnole_BTN_XP_008103839.1':0.032019)[&label=100]:0.104289,iguana_BTN_pryspry_only:0.062693)[&label=53]:0.014351,'fence_BTN_XP_042310934.1':0.066442)[&label=76]:0.029234)[&label=99]:0.295926)[&label=100]:0.985775);

end;

begin figtree;

set appearance.backgroundColorAttribute="Default";

set appearance.backgroundColour=#ffffff;

set appearance.branchColorAttribute="User selection";

set appearance.branchColorGradient=false;

set appearance.branchLineWidth=1.0;

set appearance.branchMinLineWidth=0.0;

set appearance.branchWidthAttribute="Fixed";

set appearance.foregroundColour=#000000;

set appearance.hilightingGradient=false;

set appearance.selectionColour=#2d3680;

set branchLabels.colorAttribute="User selection";

set branchLabels.displayAttribute="Branch times";

set branchLabels.fontName="sansserif";

set branchLabels.fontSize=8;

set branchLabels.fontStyle=0;

set branchLabels.isShown=false;

set branchLabels.significantDigits=4;

set layout.expansion=0;

set layout.layoutType="RECTILINEAR";

set layout.zoom=0;

set legend.attribute="label";

set legend.fontSize=10.0;

set legend.isShown=false;

set legend.significantDigits=4;

set nodeBars.barWidth=4.0;

set nodeBars.displayAttribute=null;

set nodeBars.isShown=false;

set nodeLabels.colorAttribute="User selection";

set nodeLabels.displayAttribute="Node ages";

set nodeLabels.fontName="sansserif";

set nodeLabels.fontSize=8;

set nodeLabels.fontStyle=0;

set nodeLabels.isShown=false;

set nodeLabels.significantDigits=4;

set nodeShapeExternal.colourAttribute="User selection";

set nodeShapeExternal.isShown=false;

set nodeShapeExternal.minSize=10.0;

set nodeShapeExternal.scaleType=Width;

set nodeShapeExternal.shapeType=Circle;

set nodeShapeExternal.size=4.0;

set nodeShapeExternal.sizeAttribute="Fixed";

set nodeShapeInternal.colourAttribute="User selection";

set nodeShapeInternal.isShown=false;

set nodeShapeInternal.minSize=10.0;

set nodeShapeInternal.scaleType=Width;

set nodeShapeInternal.shapeType=Circle;

set nodeShapeInternal.size=4.0;

set nodeShapeInternal.sizeAttribute="Fixed";

set polarLayout.alignTipLabels=false;

set polarLayout.angularRange=0;

set polarLayout.rootAngle=0;

set polarLayout.rootLength=100;

set polarLayout.showRoot=true;

set radialLayout.spread=0.0;

set rectilinearLayout.alignTipLabels=false;

set rectilinearLayout.curvature=0;

set rectilinearLayout.rootLength=100;

set scale.offsetAge=0.0;

set scale.rootAge=1.0;

set scale.scaleFactor=1.0;

set scale.scaleRoot=false;

set scaleAxis.automaticScale=true;

set scaleAxis.fontSize=8.0;

set scaleAxis.isShown=false;

set scaleAxis.lineWidth=1.0;

set scaleAxis.majorTicks=1.0;

set scaleAxis.minorTicks=0.5;

set scaleAxis.origin=0.0;

set scaleAxis.reverseAxis=false;

set scaleAxis.showGrid=true;

set scaleBar.automaticScale=true;

set scaleBar.fontSize=10.0;

set scaleBar.isShown=true;

set scaleBar.lineWidth=1.0;

set scaleBar.scaleRange=0.0;

set tipLabels.colorAttribute="User selection";

set tipLabels.displayAttribute="Names";

set tipLabels.fontName="sansserif";

set tipLabels.fontSize=8;

set tipLabels.fontStyle=0;

set tipLabels.isShown=true;

set tipLabels.significantDigits=4;

set trees.order=false;

set trees.orderType="increasing";

set trees.rooting=false;

set trees.rootingType="User Selection";

set trees.transform=false;

set trees.transformType="cladogram";

end;

All CDS, all domains aligned

>iguana_BTN_psOnly

------------------------------------------------------------------------------------------------------------------------------------------------------------------------------------------------------------------------------------------------------------------------------------------------------------------------------------------------------------------------------------------------------------------------------------------------------------------------------------------------------------------------------------------------------------------------------------------------------------------------------------------------------------------------------------------------------------------------------------------------------------------------------------------------------------------------------------------------------------------------------------------------------------------------------------------------------------------------------------------------------------------------------------------------------------------------------------------------------------------------------------------------------------------------------------------------------------------------------------------------------------------------------------------------------------------------------------------------------------------------------------------------------------------------------------------------------------------------------------------------------------------------------------------------------------------------------------------------------------------------------------------------------------------------------------------------------------------------------------------------------------------------------------------------------------------------------------------------------------------------------------------------------------------------------------------------------------------------------------------------------------------------aatgtaacactggatccagatacagcacaccatgacttgatcctgtctgcagaccagaagaatgtcatacaa---ggcttcctgtggcacaggctccctgaccatcctcacagatttgatacagagcgctgcgtgttggggaaggagggtttttcctcagggaggcattactgggaagtggaagtaggcgaagaagggtattgggccgtgggggtggctagggattctgtcaggaggaaagggagGttt---------agccttgatcctgatgaaggaatctgggctgtggaaaagtactgggtccaaactgtccagtaccaagctcttacc---atcccacagacccctctgccActgcgcaagagacccaggagacttgggatatacttggattatgaaatggaagaggtggcttttcatgatgttattcacaagatgcacatcttcacattcacctcagcagatttcagtggagagacaatttacccttttctgcatgtaggg---gttgggtgctggcttacactc------------------------tgcccctga------------------------------------------

>iguana_hTRIM

---------------------------------------------------------------------------------------------------------------------------------------------------------------------------------ATGGCTGCCGGCGGGAATCCCTCCGTGAAGCTCCAGGACGAAGCGACCTGCGCCATCTGCCTGGATTACTTTAAGGACCCGGTGATGATCGTCGTCTGCGGGCACAACTTTTGCCGAGCCTGCATCGGCCAATACCAGGAGGCC------------------------------------------------------------------------------------------------------------------------------------------------------------------------------------------------------------------------------------------------------------------------------------------------------------------------------------------------------------------------------------------------------------------------------------------------------------------------------------GGGTCGGGCCACGATCAGTCCCGGTGTCCCCAGTGCAGAATCCATTTCCTTTCGGGCAGCCTCCGGCCCAACAGGGACCTGGGGAATCTGGTGGACGCGATCCGGCAGCTCCGCCTG------------------------------------------------------------------------AGCGACCCCGTTTCGGGGTCGAAGAGATCGAACCCGAGCTGGAAGGTGTGCGAGAAGCACGCCGAAGCCGCCAAGCTTTTCTGCCAGGAGGACAAAGCCTTCCTCTGCCTGGTCTGCAGAGAGTCCCGAGCTCATAAGGCACACAGCGTCTTCCCCGTAGAGGAGGCGGTTCAGGACTACCAGGATCAAATGCAAAGTTTCCTGACAAGACTGACAAAAGAGAGAGAGGAAATACTGAAG---GCAAACTTGTATAATCAGAAAGAAGTTCCGCAGCTGAAGGTTGATTTGGGATCT---------------------------------GCCAGGCAGCGCATTGCATCCAGGTTTCCAAAG---------------------GAAGAAGCCAGGGCCTTGCTGCCCCTGCTAGATGGATGTGAAGCAAAGATGAAGACAGTGATCGACATGGCAAATGACTTCTCAGTGCGTGATTCCTGTCTCACCAAGCTGATTAGTGAGATAGAAGAGAAGCGTCTGCAGCCCTCCTTTGAATTCTTGCAGGACATTGGTGAT---TTGCTGAGCAGGTGTGAGAGAGAATTAGGAAAACCC---------------AGACCTGATCTGAAAGGGATAGTTGGATACATCACTGAAAATATGAAGTGGGTTCACACCAAGCTGGATCAATACCCA---------------------------------------------------------------------------------------------------------------------------------------------------------------------------------------------------------------------------------------------------GAGAAGACATCAAATCCAAAATGGATAAAAGAAAATGTGATGCTGGATCCAGAAACAGCCCATCCTCGATACATTGTGTCTAAAGATCGGAAGAGTGTACACTGG---GGGGACAGCCGTAGGGACCTGCCTTACAACCCCAAAAGATTTGAGTATGCTCGCTGTGTGCTTGGAAAGAAGGGATTCACCTCAGGAAAGCATTATTGGACTGTGGATGTAGAGGATGGAGAATACTGGGCAGTGGGTGTGGCCTTAGAATCTGTGGACAGGGATGGGGAACTT---------GATTTTGAGCCTGATGAAGGGATCTGGGCTCTGGCACGTTAT---------GATGATCAGTACAAAGCTCTTACT---TCACCTCCAACTCTTCTTGACCCAGACAATGATCCGGTAGAGATCCAGATTTCTCTGAACTATGATGCTGGAACAGTGGCTTTTTATGAT---GAAGACAAGACCCGTCTCTTTATTTTCCGGTCCATTGATTTTGAAGGGGAGAAAGTGTTCCCTTTTTTCCGGATTTCTGATTCATCGACTTCTCTACGGCTG------------------------TGTTAA---------------------------------------------

>XM_060759386.2

------------------------------------------------------------------------------------------------------------------------------------------------------------------------------------------------------------------------------------------------------------------------------------------------------------------------------------------------------------------------------------------------------------------------------------------------------------------------------------------------------------------------------------------------------------------------------------------------------------------------------------------------------------------------------------------------------------------------------------------------------------------------------------------------------------------------------------------------------------------------------------------------------------------------------------------------------------------------------------------------------------------------------------------------------------ATGAAGGTGAGAGGTGTCCCCCAGACATTTGGAGGACTCTGCTGCGTTCTGTGGCTCTCATTGTGTTTTCTTACCGAATTCGAAGGTGGCGGGAAGGTCCTGGCTTCAACACCA---------------------------------------------------------------------------------------------------------------------------------------------------------------------------------------------------------------------------------------------------------------------------------------------------------------------------------------------------------------------------------------------------------------------------------------------------------------------------------------------------------------------------------------------------------------------------------------------------------------------------------------------------------------------------------------------------------------------------------------------------------------------------------------------------------------------------------------GCCAATGTACTTTTTGATCCAATCACTGCACACCCAAATCTCCGTGTGTCTCCAAACAAGAAGACTGTGACGTGG---CTACCTGAACCCCAAAATGTTCCTGACAATCCAGAGAGATTCAATGCCAGCTATTGTTTGCTGGGTACCCCAAGATTCACCAAGGGGAAGCACTACTGGGAAGTGGAGTATGGAGGCCAGAGGGAGTGGGCAGCAGGGGTGGCCCGGGCATCTGTGCAAAGGAAAGGGTATGTC---------AGCATGACACCAGAGGAAGGGATTTGGCAGGTGGGCCTTTGGTGGCTTCGTCGCGGGGAAACTGATTCCCACCAA------------------------CCTCCCAAGCACCCTGCCAAGATCAAGATATCTCTGGATTATGAGCAGGGGACAGTGACTTTTAACCTGGAGAATAAGGTCACTGTA------------CAGAAAGCCTCTTTTAAAGGGGAGATGATTCGCCCCTTCTTCTATGTAGGC---GCAACTGTTTCACTCAAAGTG------------------------------TGA------------------------------------------

>XM_067463364.1

---------------------------------------------------------------------------------------------------------------------------------------------------------------------------------ATGGCTGTCGGGGGCCACCCTTCCGCCCGGCTCCAGGAGGATGCCACTTGCTCCATCTGCTTGGACTTCTTCCAGGACCCGGTGATGATCATCGACTGCGGGCACAACTATTGCCGGGCGTGCATCTCCCAGTGCCAGGGGCAG------------------------------------------------------------------------------------------------------------------------------------------------------------------------------------------------------------------------------------------------------------------------------------------------------------------------------------------------------------------------------------------------------------------------------------------------------------------------------------GGC---------------TCCCGTTGCCCCCGTTGCAGGATCCCTTTCCCTTCGGAAAACCTCCGGCCCAACAGGGATCTCCGGAATCTGGTGGAGGCGATCCAGCAGCTCAGCCTG---CAGCCGCTT---------------------------------------------------------------GAGAAGACGACCCGGCCAAAGCCGGCGGGAGGCGGCCCGAAGGTGTGCGAGAAGCACCAGGAAGCCGCCAAGCTCTTCTGCCGGGAAGACCAAGCCTTCCTCTGCCTGGTCTGCAGGGAGTCCCGCGCCCACAGAGCCCACACCGCCCTCCCCATCGAGGAGGCAGCCCAGGAGTACCGGGATCAAATCCAGACTTTTTTGAAAAAACTGGCAAAAGAAAGAGATGAAATGCAGAGA---GTTAACTTGTCCAATAAGCACAATTATCAACAAATGAAGAGTAATTTGGAATCT---------------------------------GCCAAGCAGCGCATTGGATCCAATTATCCTAAA---------------------AAAGAAGCCCAGACCTTGATGCCCCTGCTAAACACGTGTGAAGCCAATCTGAAGATACTTAGTGAGGCACCCAATGATTTA------------TCCTGTCTTAATAAGCTGATTAGTGAAATAGAAAAGAAGTCCCAGCAGTCATCCTTTGGATTCTTGCAGGGCATTGGTGAT---TTACTGAACAGATGTGAGCGAGAAACA---AGAAGACCA------------AGACATGATCTGAGAGCGATCATTGGATCCTTACATGAAAAACTATGTGTGGTTAACACCAAGTTGAATCAATGTGCA---------------------------------------------------------------------------------------------------------------------------------------------------------------------------------------------------------------------------------------------------GAAAACACACCAAAGAAACCATGGATAAAAGAAAATGTGACGTTGGATGCAAAAACAGCCCATCCTCGGTTCTTTGTGTCTGATGATCGGAAAAGCGTACACTGG---GCGAAGACCCGCCAGGATCTGCCATACAGCTCCAAGAGATTTGAGTTTGCCCGCTGTGTGCTTGGGAAGAAAGGATTCACCTCTGGAAAACATTATTGGATTGTGGATGTAGAGGATGGTGACAACTGGGCGGTGGGTGTAGCCCAAGAGTCTGTGGACAGGGGAGGGAAACTT---------GATTTCGAGCCTGATGAGGGGATCTGGGCTATGGCACGTTAC---------GATGATCAGTATAAGGCTCTTACT---TCACCCCCTACCCTTCTGGATCTAGACTATGATCCAGTAGAGATCCAGGTTTCTTTGAACTATGATGCTGGAATAGTGCATTTCTATGAT---GAAGATAAGAAACCCCTCTTTACTTTCCGATCATTCGATTTTGAAGATGAGAAGGTCTTTCCTTTTTTCCGAATTGGTAATTCATCCACATCTTTGTACCTG------------------------CTCTAA---------------------------------------------

>XM_067463351.1

------------------------------------------------------------------------------------------------------------------------------------------------------------------------------------------ATGAAATTTCCTGTCTTCCCTTATGCCTCGGAAATGAGCTGCTTTCTGTTCCTTTTCTTTGTTATTTCACCTATTAATAACGTGAGCCCTGTACCATTTACCGTGACTGGGCCACTCCACCCAGTCATTGCC---------------------------------------------------------------------------------------------------------------------------------------------------------------------------------------------------------------------------------------------------------------------------------------------------------------------------------------------------------------------------------------------------------------------------------------------------------------------------------------TCTTTGAGTGAGGATATTGTGCTGCGCTGTCATCTGTCCCCCAGAATGAGTGCAGAAAACATGGAAATAAAATGGTTTCGCTCCCAGAATTCTTCATACGTGCACCTTTATCACAGT------------------------------------------------------------------------------------------GGCAAAGAGCATTTGGAAAAGCAGCAGCCAGAATACCAAGGAAGAACAGAATTGTTGACAGAGGGCATTGGGGATGGTAAGATCAGCCTGAGGATTTCTAATGTCAGCTTATTTGATGAAGGACAGTATCACTGCTCTGTGAAGAAC------AGGAGCTTCCATCAGGAAGTCACACTGGATATAAAAGTGGCTGCATCAGGCTCTACTCCTCTCATTTCC---ATTGAACGTTATCAGGAGAGAGGGATCTACTTGGTTTGCCGATCCAGGGGTTGG---------------------------------TACCCGGAGCCTGATATCTTTTGGAGAGATCCCAGTGGGAGGCATCTGTCCTTTTTGGCTGAGGAAACATCCCAGAAGAACGATGGGCTGTTTGAAGTACAGAAGGACATTCTCCTAACAGAAGGTTCAAACCAGTCCTTGACTTGTGTGGTCCGGAACAGGCTCCTCAACCAAGAGAAAGAATCAACCATTCACTTTGCAGATAACTTCTTCCCAAAGATTGACCCCTGGATAGTTGGTCTATGTGTATCCATCCTGGTTCCTGTTGGCTTACTCGTTGCTATTCTGTACCTGATTAACAGAAACAGAAAGCTCATCACGGAACTGAGTTGG------------------------------------------------------------------------------------------------------------------------------------------------------------------------------------------------------------------------------------------------------------------------------AGGAATATGGTGGTGCCAATA---GAGAAAGCCAATGTAACACTGGATCCAGACACAGCAAACAAGGAACTGATCCTGTCTGCAGACCAAAAGAATGTTATACAA---GGCTTCATGTGGCACAACCTACCTGACCATCCTTGGAGATTTAATAAGGAGCGCTGCATTTTGGGAAAGGAGGGCTTTTCCTCAGGGAGGCATTACTGGGAAGTGGAAGTTGGGGAAGAAGGGTATTGGGCCGTGGGGGTGGCTAGGCAATCTGTCAGGAGGAGAGGAAAGCTC---------AGCCTTGATCCGAATGAAGGCATCTGGGCTGTGGAGAAGTCCAGGGTCCAAACTGTCCAGTACCAGGCTCTTACC---GTCCCACTGACTCCTCTGCTGCTGCGCAAGAGACCCAGGAAAGTTGGGATTTATTTGGACTATGAGATGGGAAAGGTGGCCTTTCATGATGTCAATCACAAGATGCACATCTTCACATTCTTTGAAGCAGATTTCAACAGGGAAACAGTTTTCCCTTTCCTGCATGTGGGA---ATAGGATGCTGGCTTACAGTC------------------------TGCCCTTGA------------------------------------------

>XM_060270853.1

---------------------------------------------------------------------------------------------------------------------------------------------------------------------------------ATGGCCGCCGCCGGAGACCCCTCCATAACTCTCCAGGACGAAGTGACTTGCTCCATCTGCTTGGATTACTTCCGCGACCCGGTGATGATCATCGACTGCGGGCACAGCTTTTGCCGAGCCTGCATCTCCCAGTGCCCGGAG---------------------------------------------------------------------------------------------------------------------------------------------------------------------------------------------------------------------------------------------------------------------------------------------------------------------------------------------------------------------------------------------------------------------------------------------------------------------------------------GGGCCGGGCCGCGACCTCTCCTCTTGCCCCCAGTGCAGGATAGCTTTTCCCCGGGGCAACCTCCGGCCCAACAGGCACTTGGCGAATATGGCCGAAGCGATACAGCGCCTCCGCCTG---------------------------------------------------------------------CAGTGCCTAAGCCTAGCCGAAGGGGAGCCCAAGAAGGAAGCGCCGAAGTTGTGCGGGAAGCACTACGAAGCCCTCCGGCTCTTCTGCCAAGAGGACCGCGCCTTCATCTGCCTGGTCTGCAGGGAATCCCGGGCGCACAAAGCGCACGCCACCCTCCCCATCGACGAGGCTGCCCAGGATGCTAAGGATCAGCTCCAAAGTTGCCTAACAACCTTGAAGAAGGAGAGAGATGAAATTTTGAAGAGGATTGAAGGGTGCAGTGAGAGA---------GAAATGAAGAATCGTGTGGGAAGC---------------------------------ATCAGGGATCAAGTTGCGGGAAACTGGGAA------------------------------------------------CTGCTACTTGCGTGTGCAGAGATCATGGAGTCAATCGACAAAATAGCCAGTGAATTCTCAGAGCGCAGCTCTCATCTCGCCAAACTGATTAAAGAGGTAGAGGAGAAGTGCCTGAAGTCTGCCTTTGAACTCCTGCAGGATATGGATCCC---TTGCTGAGCAGATGCAACATAGAAATATCTGGGAAACCC------------AAAGCTGATCTGAATAAGAAAGTTGAGTTGCTTAATGAAAAACTACAGCTGGTGCAGACCAAGTTGCGTCAGCTTTCA------------------------------------------------------------------------------------------------------------------------------------------------------------------GCAGGTGTGCCACAGCAGCTCAATCAGCACAGATATACTTCTGATTTTCACTATCCAGTATGTGACGCAGACTATGTTGATGCCAACACGGTGAAATCAAAATGGAAAAGAGAAAACGTGGTGTTTGATCCGGAAACAGCCCATCCCCGATATATTGTGTCTTCTGATGGAAAATCAGTATGGTGG---GGAAAGGTCCGCCAGGACTACCCCTACAGTTCAATTAGGTTTGAATACGCCCGCTGTCTACTTGGAATGTGGGGATTCAGCTCAGGGAAACATTATTGGACGGTGGATGTTGAGGGTGTCAACCATTGGGCTGTGGGCGTAGCCAGAGAGTCTGTGGAGAGGGATAGAGAAATT---------AATTTTGAACCTGATGAAGGGATCTGGGCAGTGGGCTTTAGC------CGCAATGATCAGTTCAAAGCTCTGACT---TCACCACCTACCTATTTGGACCCAGAAGAGGGCCCAACACAAATCCAAGTTTCGTTGAACTACGAAGCAGGGACTGTGGCTTTTTATGATGCTGAAGATAAGACCCGCCTCTTTATTTTCGAGTCAATTGATTTTGAAGGGGAGGAAGTTTTCCCCTTTTTCCGGATTGTAGATCCATCAGTTTGCCTCCAGCTG------------------------TGCTACTAA------------------------------------------

>XM_035105027.2

------------------------------------------------------------------------------------------------------------------------------------------------------------------------------------ATGGCTACCGCGGATCCTGCAAAACACCTTCAGGATGAAGTCACTTGTTCCATTTGTCTGGATTATTTCAAGGATCCAGTAATGATCACGGAGTGCCAGCACGACTTTTGCCGATCCTGCATCACTAAGTATTGG------------------------------------------------------------------------------------------------------------------------------------------------------------------------------------------------------------------------------------------------------------------------------------------------------------------------------------------------------------------------------------------------------------------------------------------------------------------------------------------AAAAGATCGGGAGCTTCCATTTGCTGCCCGGATTGCAGAAGAAAAGCTTCCTGGCAAAGCCTAAAACCCAACCGGCGCCTGGCGAACATGGTGGAAGCAGCTAAGCAGCTCCGCCTG---------------------------------------------------------------------------------------CAGCTGGAGCAAAGTCCAGGAAGGGAGAAAATGTGCAAGGAGCACAGGAAGCCTCTGAGTCTCTTCTGCAAAACAGAAAATACCCTTATTTGTATGTTCTGCGAGAGATCCAAGGCTCACAGAAACCACCGTGTTATTTCTGCAGAAACAGCTGCTGCAGATTACAAGGGTATACTACTTGGACATCTAAAGGAACTGAAAAAAGAAAGAAACAAGATTCTGTCG---GTTAAATCAAATGGTGAGAAGCCATGCCAGGAGCTATTGAAAGAGGCAGAAGCC---------------------------------AAGAGGGAGGAGATTGTGTCTGAATTCCAACACCAGCGACAGTTTTTGGAGGAACAAGAGCAACGCCAGCTGGCGAGGTTGGAAGACATAAAAAGAGAGGTGGAGAAGAGAAGAAAAGATTATGCAAACAAATTTGAGGATGAAATATCCTACATCAGCGATCTGATTGGTGAGATAGAAAAGAAGGGCAATCAATCAGCAGATGAATTCTTGGAGGGTATCGGAAGC---ATACTGGACAGATGTGAGACAGAGAAGTTTCAGTTTCCAGCTACACCTGCTTCTACAGACATGAAGGAAAGGCTCAGGCAATTCTCTCAAGAAAGTGCATCTTTGACGAGTTCTCTGAGGAGAGTCAGA------------------------------------------------------------------------------------------------------------------------------------------------------------------------------------------------------------------------GACTGTTCTGAGCCCAACTGTTCTCAGTCCAACAAGGTCAACTCAAAATGGAAAAGAGAAAATGTGGTGTTGGATCTGGAAACAGCCCATCCCCGGTATGTTGTGTCTTCTGATGGAAAATCAGTATGGTGG---GGAGATGTCCGTCAGGACTACCCCTACAATCCAAAGAGATTTCAATACGCTCGCTGTGTGCTTGGAACGCAGGGATTCAGCTCTGGGAAACATTATTGGACGGTGGATGTCGAAGATGTTGACAATTGGGCTGTAGGCGTAGCCAGAGAGTCTGTGGAGAGGGATAGAGAAATT---------GATTTTGAACCTGATGAAGGGATCTGGGCGTTGGGCTTTAGT---------TATGATCAGTACAAAGCTCTGACT---TCACCTCCTACCATTTTGGAACCAGAAGAGGATCCAATAGAGATCCAAGTTTCTTTGAACTATGAAGCAGGGACTGTGGCTTTTTATGATGCTGAAGATAAGACCCGCCTCTTTATTTTCAAGTCAATTGATTTTGAAGGGGAGGAAGTTTTCCCCTTTTTCCGGATTGTAGATTCATCAACTCCCCTCCAGCTG------------------------TGCTCTTAA------------------------------------------

>XM_035106984.2

------------------------------------------------------------------------------------------------------------------------------------------------------------------------------------ATGGCTGCGCCGAACCCAGAGAAAAGCCTCCAGGATGAAGCCACCTGTTCCATCTGCCTGGATTATTTCCAGAACCCGATGATGGTCATCGACTGTGGGCACAACTTCTGCCGGGACTGCATTGCCCAGTGCTGTGAA---------------------------------------------------------------------------------------------------------------------------------------------------------------------------------------------------------------------------------------------------------------------------------------------------------------------------------------------------------------------------------------------------------------------------------------------------------------------------------------GGGTCCATTGCCAACGTGTTCTCCTGCCCGCAGTGCAGAAAACCCTTCCCTTGGCAGAACCTGCGGCCAAACAGGCACCTGTGGAATATCGTTGAGCTGGCCATGCAGTTCAGCATG---------------------------------------------------------------------------------------CGGCGAGCCAAGGAGTCGGGAGAACAGAAGCTCTGCAAGAAACACCAGGAGCCCCTGAAGCTCTTCTGCGAATACGATCAAACGTCCATCTGCGTGGTTTGTGACCGGTCCAAGCTGCATAAATATCACAATGTGATTCCTATAGAGGAGGCTGCTCAGGTTTACCAGGAGAAAATACACCACTGCTTGCAAACTTTGAAAGAAGAGAGAGACAAGATTTTATCA---TTTAAATTAGATGCAGAGAAGCCAAGCCAGGATCTGCTGGCAGAAACCGAAGCC---------------------------------GAGAGGCAGAAGCTTATGTTGGAATTTCAGCAGCTCCACCAGTTTCTAGAGAAAGAAGAACAACTCTTGTTAACCCAGCTGCAGAAGCTGGAGATAGAGATAGAGAAGAAGAAGGAAGAGCATGCAGCTAAATTTTCCGAGGAGATTTCCCATATCAGCACCCTAATTGGCGAGCTGGAGCAGAAGGGTCAGGAGCCGCCAGGTGGGCTGATGCAGGATATTGGAAGT---ATCTTCAACAGGTGCAAGAGGGGCAAATTTCAGTTTCCTACTCCACCTGATTCTTCGGAGCTCAGTACAATGGTTCAGGAATTCTCCAAAGAGAGTGCTTCACTGCAGGGCAAACTGAGGAAATTCAGA---------------------------------------------------------------------------------------------------------------------------------------------------------------------------------------------------------------------------------------------------GAGACTTTGGATGAACAAAAGTGGATAAATGAGAACGTGAAACTGGACCCCATGACGGCTCACCCTCGATTCATTGTGTTTGATGGCCAGAAAGCTGTTGCATGG---GGATCTGTCCGCCAGGAGCTGCCCTACAGCACCGGGCGATTTGACCCCACCCGCTGCGTGCTTGGCTGTCAGGGATTCACGTCCGGGAAGCACCATTGGACAGTAGATGTGAGCAAGGGAACTTTCTGGGCTGTGGGTGTAGCCAAAGAATCTGTGAAGAGGAAAGGACAGCTG---------AATATGGTCCCTGAGGAGGGAATCATGGCTGTGGGTTTTAAC---------AACGGCCAGTACAAAACCTTAACG---TCGCCCCCGACTGTTCTGAACCCAAGTAAACACCCCCAAAAGATTCAAATCCGTTTGGACTATGAAGGAGGAACGGTGGGGTTTTTTGATGCGGAAGAAAAGAAGCGCCTTGTTCTCTTCAGTGTCAAATTT------GTGTCCAAAATATTTCCTTTCTTCCGAGTTGGGGACATGAACACTGTGCTCCGCCTG------------------------TGTTAA---------------------------------------------

>XM_060270861.1

------------------------------------------------------------------------------------------------------------------------------------------------------------------------------------------------------------------------------------------------------------------------------------------------------------------------------------------------------------------------------------------------------------------------------------------------------------------------------------------------------------------------------------------------------------------------------------------------------------------------------------------------------------------------------------------------------------------------------------------------------------------------------------------------------------------------------------------ATGTCGTTTTCTCTGGTCTCTCGGATGAGCTTCCGGAACTTTCCAACGAGAAGCAGTGTT------------------------------------------------------------------------------------------------------------------------------------------------------------------------CAGCCGACGCTCAAAGGATTCTATTGTATCCTGTGGCTAATGCTGTGCTTTCTCGCGGAGCAGAGAGGTGGTGGGAAAGTCCTGGCTTCACTACCA------------------------------------AAGGGATGCTACCGA------GCAACAGGCCCTGCTCCCCTTATTTCC---ATCAAAGGCTATCAGGACCAAGGGATCCGAATAGTTTGCCGATCCAGTGGTTGG---------------------------------TACCCAAAGCCTGAGATCCTATGGAGAGATCCCAATGGAAGGCATTTGCCTTCACTGGGACAAAAAATTTCCATGGAGGGCAATGGGTTGTTTGAGGCGCAGAACGACATCATCCTAACAGGAAGTTCAAATCATAGCCTAACCTGTGTGGTCCGGAACAGCTTCCTCAACCAGGAAAAGGAATCAACTATTCATATAGCAGATGACTTTTTCCCAAAGATCTCTCAGTGGATAGCTGGCTTGTGTGTCTCCCTCATGGCTCTGCTCGGCTTCATTCTCCTCCTCCTTTACCTGTTTAATATAAACAGAAAGCTTTATGCAGAACTGAGTTGG------------------------------------------------------------------------------------------------------------------------------------------------------------------------------------------------------------------------------------------------------------------------------AGGAACTTGGTGATGCCAATC---AAAAAAGCAAATGTAACCCTGGATCCGGACACAGCGCACGACGAGATCATCCTGTCTGCGGACCAGAAGAACGTCATACAA---GGCTTCACGTGGCACACCCTGCCTGACAGCCCCCTGAGATTCAGCGTTGAGCGCTGCATTTTGGGAAGCGAGGGCTTTTCCTCAGGAAGGCACTACTGGGAAGTCGAGGTGGGCGAGGAGGGGTACTGGGCCGTGGGGTTGGCAAGGGGTTCCGTGAGGAGGAAGGAGTGCCTC---------AGCCTTGACCCCAGCGAAGGGATCTGGGCCATGGAGAAGTGC---------CGGATCCAGTACCAAGCTCTCACC---GTCCCCGAGACCCCCCTGTCTTTGAGGAAGAGACCCAGGAAACTTGGAGTTTACTTGGACTACGAGATGGGGCAAGTGGCGTTTCATGACGTCAACCACAAGACACCCATCTTCACTTTCCCCTCCGCTGCTTTCAACGGGGAGGAGGTCTTCCCTTTCCTGCAAGTCGGG---GTGGGATGCTGGCTTCGAGTG------------------------TGCCCCTGA------------------------------------------

>XM_060270855.1

---------------------------------------------------------------------------------------------------------------------------------------------------------------------------------ATGGCCGCTGCCGGAGACCCCTCCATAACTCTCCAGGACGAAGCGACTTGCTCCATCTGCTTGGATTACTTCCGCGACCCGGTGATGATCATCGACTGCGGGCACAGCTTTTGCCGACCCTGCATCGCCCAGTGCCCGGAG---------------------------------------------------------------------------------------------------------------------------------------------------------------------------------------------------------------------------------------------------------------------------------------------------------------------------------------------------------------------------------------------------------------------------------------------------------------------------------------GGGCCGGGCCGCGACCTCTCCTCTTGCCCCCAGTGCAGGATAGCTTTTCCCCGGGGCAACCTCCAGCCCAACAGGCACTTGGCGAATATGGCCGAAGCGATACAGCGCCTCCGCCTG------CAGCGC---------------------------------------------------------------CTCTCCCTGGCCGAAGGGGAGCCCAAGAAGGAAGCGCCGAAGTTGTGCGGGAAGCACTACGAAGCCCTCCGGCTCTTCTGCCAAGAGGACCGCGCCTTCATTTGCCTGGTCTGCAGGGAATCCCGGGCGCACAAAGCGCACGCCGTCCTCCCCATCGACGAGGCTGCCCAGGATGCTAAGGATCAGCTCCAAAGTTGGTTAACAACCTTGAAGAAGGAGAGAGATGATATTATGAAG---CAGAAAAGTTACACTGAGAAAAAAAATCAGGAATCAAAGAAAGGTGTGGAAAAC---------------------------------ATCAGGAATCAAATTGTGGGAAAGCGGGAACTGCTA------------------------------------------------------------GAAGCACTTGAAGAAATCGACAAAATAGCCAGAGAATTCTCAAACCGCAGTTCTCATCTCGCCGAGCTGATTAAAGAGGTAGAGGAGAAGTGTCTGCAGTCTGCCTTTGAACTCCTGCAGGATATGGATCCC---TTGCTGAGCAGATGCGACATAGAAATATCTGGGGAACCC------------AAAGCTGATCTGAATAAGAAAGTTAAGTTGCGTGATGAAAAACTACAGCTGGTGCAGACCAAGTTGCGTCAGCTTTCA---------------------------------------------------------------------------------------------------------------------------------------------------------------------------------------------------------------------------------------------GACCGTTCTCATTCCAACACGGCGAAATGGATAAAGGAAAACGTGGTGTTTGATCCGGAAACAGCCCATCCCCGATATATTGTGTCTTCAGATAGAAAATCAGTATGGTGG---GGAAAGGTCCGTCAGGACTACCCCTACAGTTCAAATAGGTTTGAATACACTCGCTGTGTACTAGGAATGCGAGGATTCTACTCTGGGAAACATTATTGGACGGTGGATGTTGAGGATGTCGACCATTGGGCTGTGGGCGTAGCCAAAGAGTCTGTGGAGAGGGATAGAAAAATT---------GATCTTGAACCTGATGAGGGGATCTGGGCAGTGGGGTTTAGC---------AATGATCAGTTGAAGGCTCTGACT---TCACCTCCTACCCCTTTGGAAACAGAAGATGTCCCAACACAGATCCAAGTTAAGTTGAACTATGAAGCAGGGTCTGTGGCTTTTTATGATGCTGAAGATAAGACCCGCCTCTTTATTTTCCGGTCAATTGATTTTGAAGGGGAGGTAGTTTTTCCTTTTTTCCGGATCGTCAATTCATCAACTTGCCTCCAGCTG------------------------TACTCTTAA------------------------------------------

>XM_035106987.2

---------------------------------------------------------------------------------------------------------------------------------------------------------------------------------ATGGCCGCTGCCGGAGACCCCTCCATAACTCTCCAGGACGAAGCGACTTGCTCCATCTGCTTGGATTACTTCCGCGACCCGGTGATGATCATCGACTGCGGGCACAGCTTTTGCCGACCCTGCATCGCCCAGTGCCCGGAG---------------------------------------------------------------------------------------------------------------------------------------------------------------------------------------------------------------------------------------------------------------------------------------------------------------------------------------------------------------------------------------------------------------------------------------------------------------------------------------GGGCCGGGCCGCGACCTCTCCTCTTGCCCCCAGTGCAGGATAGCTTTTCCCCGGGGCAACCTCCGGCCCAACAGGCACTTGGCGAATATGGCCGAAGCGATACAGCGCCTCCGCCAG---------------------------------------------------------------------CAGCGCCTAAGCCTAGCCCAAGGGGAGTCCAAGAAGAAAGCGCCGAAGTTGTGCGGGAAGCACTACGAAGCCCTCCGGCTCTTCTGCCAAGAGGACCGCGCCTTCATCTGCCTGGTCTGCAGGGAATCCCGGGCGCACAAAGCGCACGCCACCCTCCCCATCAACGAGGCTTCCCAGGATGCTAAGGATCAGCTCCAAAGTTGCCTAACAACCTTGAAGGAGGAGAGAGATGAAATTTTGAAGGGGATTGAAGTGTGCAGTCAGAGAGAAATTCAGGAAATGAAGAATCGTCTGGGAAGC---------------------------------ATCAGGGATCAAGTTGCGGGAAACTGGGAA------------------------------------------CTACAAGTTGTGGGAAACTGTGCAGAGCTCATGCAGTCAATCGACAAAATAGCCAGTGAATTCTCAGAGCGCAGCTCTCATCTCGCCAAACTGATTAAAGAGGTAGAGGAGAAGTGCCTGAAGTCTGCCTTTGAACTCCTGCAGGATATGGATCCC---TTGCTGAGCAGATGCAACATAGAAATATCTGGGAAACCC------------AAAGCTGATCTGAATAAGAAAGTTGAGTTGCTTAATGAAAAACTACAGCTGGTGCAGCGTCAGCTTTCAGAACATCCA------------------------------------------------------------------------------------------------------------------------------------------------------------------------------------------------------------------AAACCGGCCAGTCTGAGAGCAGACTATGTTCATGCCAACACGGTGAAATCAAAATGGAAAAGAAAAAACGTGGTGTTTGATCCGGAAACAGCCCATCCCCGATATATTGTGTCTTCAGATGGAAAATCAGTATGGTGG---GGAAATGTACGCCAGGACTACCCCTACAGTTCAATTAGGTTTGAATACGCCCGCTGTCTACTTGGAATGCGGGGATTCAGCTCAGGGAAACATTATTGGACAGTGGATGTTGAGGGTGTCAATCATTGGGCTGTGGGCGTAGCCAGAGAGTCTGTGGAGAGGGATAGAGAAATT---------AATTTTGAACCTGATGAAGGGATCTGGGCAGTGGGCTTTCGC------CGCAATGATCAGTTCAAAGCTCTGACT---TCACCACCTACCTATTTGGACCCAGAAGAGGGCCCAACACAAATCCAAGTTTCGTTGAACTACGAAGCAGGGACTGTGGCTTTTTATGATGCTGAAGATAAGACCCGCCTCTTTATTTTCGAGTCAATTGATTTTGAAGAGGAGGAAGTTTTCCCCTTTTTCCGGATTGTAGATCCATCAGCTTGCCTCCAGCTG------------------------TGCTACTAA------------------------------------------

>XM_060239254.1

------------------------------------------------------------------------------------------------------------------------------------------------------------------------------------ATGGCCTCAGCAAATCCAGTGAAAGACCTTCAGGATGAAGTCACTTGTTCCATCTGTCTGGATTTCTTTCAAGACCCCGTGATGATCGTAGAATGTGGGCACAATTTCTGCGGCGCCTGCATCACCAAGCACTGG------------------------------------------------------------------------------------------------------------------------------------------------------------------------------------------------------------------------------------------------------------------------------------------------------------------------------------------------------------------------------------------------------------------------------------------------------------------------------------------CAAGGCTCTGATCGTAGCGATCTTTGCCCCGAGTGCAGGCGTTTGTTTTCCTACCAGAGTCTTAAGCCGAACAGACAACTGGGGAAGACTGCAGAAAAAGTGAAACAGGTCAGCGCG---------------------------------------------------------------------------------CAGCTGAAAAGCAACAAAGCAGACACGCTGGAGAGAATGTGCAAGCAGCACCAGAAAGCTCTGACTCTCTTCTGCAAAACTGAAGAGCGCCTCATTTGCCAGGTCTGCGAGAAAACCAAGGCTCACAGAAGTCATGCAGTTGTTTATGCAGGAGAAGCGGCTGCAGATTACAAGGAGGTACTGCTGAGGCATCTAGCAGATCTGAAGCAAGAAAGAAACAGGATTCTCTCC---ATTCAATCAAATGGAGAAAAGCCTAGTCAGAACTTATTGAAAGAGGTGCAAGCT---------------------------------ATGAAGCTGGAGATTGTATCTGAATTCCATGAACAGCAGCAGTTTTTGGAACAACAAAAGCAGCTCCAGCTGTCCAAGCTGAGAGACCTAGAAAAGGAGATTGAGAAGAGAAGAGCTGACTATGCAAGCAAGTTTGTGGATGAAATATCTTACATCACTGATCTGATTGGTGATGTGGAAAAGAAGGCTAAGCAACCAGCAAATGAGCTCCTGCAGGATATTAGATAT---GAGTTGGACAGATATGAGAAGGAGGAATTTAAGTGGCCAGTTGTACCTCATTCTTCAGATGTGCAGAAAAAGCTCAATACATTTTCTCAAGACTGCATGTATCTGCAACGTATTCTAAGGAAATTCAGA---------------------------------------------------------------------------------------------------------------------------------------------------------------------------------------------------------------------------------------------------GAGACTCCACTGAAACCAACATGGATCAAAGAAAATGTGTTGCTAGATCGAGAAACAGCCCATCCCCGATATGTTGTGTCTGAAGATTGGAAAAGTGTAAGATGG---GGGGACATCCGTCAGGAATTCCCCTACAACCCAAAGAGATTTCAATATATTCGTGGTGTTCTTGGATGTGAAGGATTCACTTCTGGAAAACATTATTGGACTGTGGATGTAGGAAATGGAGGTTGTTGGGCTGTGGGAGTGGCCAGAGAATCTGTGGAGCGGGAGGTGCAAATT---------GATTTTCAACCAGATGAGGGAATCTGGGCGATTGCACATTCC---------AGTGATCAGTACAAAGCTCTAACC---TCACCTCCTACTCTGCTGGATGTAGAAGATGAGCCAACACAGATCAAGGTTTGTTTGAACTATGAAAAGGGGACAGTTGCTTTTTATGATGTTTACGATGATACCCGCCTCTTCAAATTCTTTTCAGTTAATTTTGAAGAGGAGGAAGTTTTTCCTTTTTTCCGGATTGTTGATTCATCCACAGTTCTGGAATTG------------------------TGTTCTTAA------------------------------------------

>XM_060238766.1

------------------------------------------------------------------------------------------------------------------------------------------------------------------------------------ATGGCTGCTCCAGATCCAGAAAAAAATCTCCAGAATGAAGCCACGTGTTCCATCTGCCTGGATTATTTTCAGGATCCGGTGATGGTCATAGACTGTGGGCATAATTTCTGTCGAAACTGCATCGCACAGTGCCACAAT---------------------------------------------------------------------------------------------------------------------------------------------------------------------------------------------------------------------------------------------------------------------------------------------------------------------------------------------------------------------------------------------------------------------------------------------------------------------------------------GGGTCCATTTTCAGACTCCTCATCTGCCCACAGTGCAGAAAGCCCTTTCCTTTGAAAAACCTTCACCCCAACAGGCATCTGTGGAACATTGTGGAACTTGCCAAACAGTTCAGCAAG---------------------------------------------------------------------------------------AGACGGGCAAATGAAACTGGCGACCAGAAGCTGTGTGAGAAACACCTGGAGCCGCTAAAACTGTTCTGTGAACATGACCAGACTTCCATTTGTGTGGTGTGTGACAGGTCCGTGGCACATAAGAAACACAATGTGGTTCCCATAGAAGAGGCTGCTCAGGTTTACAAGGATAAATTGCATCACTATTTGCAGATGCTGAAAGAAGAAAAAGGAAAGATTTTGTCA---CTTAAATTGGATGTAGAGAAGCCAAGCCAAGATCTGTTGAAAGACACAGCTGCT---------------------------------GAGAGGCAGAAATTATTATCAGAGTTTCAACAGCTTCATGAGTTTCTTGAGAAACACAAGCAGCATTTATTGTCCCAAGTGGATGAGGTGGAAAAAGAAGTTGAGAGAAAAAAGATGGGAGATACAGCAAAATTTTCTGATGAGATTTCCCGTCTTGGGGCCCTAATCCAGGAGCTTGAGGAGAAGTGCCAGGAGCACCCAGGTGGGTTCCTCCAGGATATTGGAAGT---ACTCTTAACAGATGTAAGGTGAACAAATTTCAGCCTCCTGCTGTACTTGATTGTTCTGACCTTAAGAGAAGACTCCATGTATTATCTAAGGAAAGTGAATCACTGCAGGAGGAATTGAATAAATTCAAA---------------------------------------------------------------------------------------------------------------------------------------------------------------------------------------------------------------------------------------------------GCAATACTGTCTGAACCAATATGGATTGAAGAGTATGTGGTGCTGGACCAAGGTACCGCTCATCCTCGATTTGTTGTGTCTGATAATCGGAGAATGGTAACATGG---GGACCTTTTCGGCAGGAAGTGCCCTACAGCCCTGAGAGATTTGACCCAGCCCGCTGCATGCTTGGTTCTCATGGATTCTCATCAGGGAAACATCACTGGACAGTGAATGTGGAGGATGGAACTTTCTGGGCTGTAGGTGTGGCCCAAGAGTCTGTAAAAAGAAAAGGACAGTTC---------CATTTTGTGCCTCAGGAAGGGATTTGGGCTGTGGGCCTTCAT---------AATGGTCAGTACAGAGCTCTGACT---TCACCTCCTGACCTTTTGCTTCTAAATAGCCCCCTTAACAAGATCCAGGTTTGTCTGGATTATGAAGGAGGGACTGTAGCTTTCTGTGATGCTGAACAACAGCGTCAGTTCTATTGCTTT---CCAGCAAACTTCGAGGGAGAAAAACTCTTTCCTTTCTTCCGTGTTGGGGACATGAAGACTACCCTTTTCCTT------------------------TGCTAA---------------------------------------------

>XM_060238767.1

------------------------------------------------------------------------------------------------------------------------------------------------------------------------------------ATGGCTACCAAGAGCCCTACAAAGAGTTTCCAAGATGAAGCTACTTGTTCCATCTGCTTGGAATATTTTAGAGAGCCTGTGACCCTC---AACTGTGGACACAATTTCTGCCAAGCCTGCATAGCCCAGTACTGG------------------------------------------------------------------------------------------------------------------------------------------------------------------------------------------------------------------------------------------------------------------------------------------------------------------------------------------------------------------------------------------------------------------------------------------------------------------------------------------GAAACTCTGGAGGCTTCCACGACCTGTCCTCAGTGCCGAGAAACGACTCAGCAAAGAAGCTTCAGGCCAAACAAGCAGATGGCGAACCTTGTCAAGATTGTCAAGCAGATGAAGGCT---------------------------------------------------------------------------------------GAGCAGGTGATGCCAGCAGGGGAGGAGAGAGCATGTGAAAGGCATCAGGAGGGTCTGAAACTCTTCTGTGAGGAAGACCAAACTCCCATTTGTCTCATCTGCCATCTGTCCCGGGAACACAGAGACCACACTGTCTTTCCCATAGAGGAAGCTGCTCAGGATTATAAGGAAAAAATTCAAATTCACTTGAGTATCCTAAAGGAAGAGAAAAGAGAACAGGAGAAA---TTTCATACAGCTGGGAAGCGCAAATGCCAGAAGAATATTAAAAAGATAGAGGCT---------------------------------GAGAAACAGAAGGCTGTGTCTGTATTTCAAGAGATGAGATGCTTTCTAGAGGAACAAGAACAGCTCATCCTTGCCCCACTGGAGCAGTTGAAGGAAAAGTTTCTGAAAATGAGAGATGAATTGCTCACAAAACTCTCCACAGAGGTCTCCTGTCTTGCTACACTAATTACTGAAATGGAGAAGAAGTGTCAGCAACCAGCCAGTGAATTCTTGGAAGACATCAGAAGC---ACCTTGAACAAGTGTCAGCAAAAAGAAGTTCAGAAGCCA---------------------------------------------------------------------------------------------------------------------------------------------------------------------------------------------------------------------------------------------------------------------------------------------------------------------------------------------------------------------------------------------------------------------------------------------------------GCAGATATTCCTCCTGAGCTAGAAAAGAGTCTTGTTAACCTGAGTGAACAGAATGCTGTTCTA---ACCACGGCTTTTAAAAAATGTAAAGATCTTCTGTCATCTGAACTGAAG---------------------------------------------------------------------AAATATGATCCTGATGTT------------------------------------------------------------------------------------------------------------------------TATGAAATGGGTAAGATT------------------------------------------------------------AAACATAAAGTTTATGGCTAT------------------------------------------------------------CTATCCTAA------------------------------------------

>XM_060239263.1

------------------------------------------------------------------------------------------------------------------------------------------------------------------------------------------------------------------ATGAGTTCCTTCGTATTCATTCTCATTATTTCTTTCATTGTTTTCCCTGTGAACAACATGGGCCCAGTCCAATTGACAGCGATTGAACCACTTGTCTCAGTTACTGCT---------------------------------------------------------------------------------------------------------------------------------------------------------------------------------------------------------------------------------------------------------------------------------------------------------------------------------------------------------------------------------------------------------------------------------------------------------------------------------------ATTTTGGGTGAGGATGTTGTGCTACACTATCATCTGTCCCCCAGGACAAGTGCTCAGGACATGGAGATAAAATGGTTCCGTTCTCAGAACTCTTCATATGTACACCATTATCACAAT------------------------------------------------------------------------------------------GGCAAAGATTATTCTGAAAGGCAGCAGCTAAAATACCAAGGAAGGACTGAGTTTCTGAGAGATGGCATTGATGCGGGAAAGATTGACTTGAGAATCTTTAATGTCAGCCTTTTTGATGAAGGACAGTACCATTGTTCTCTGAAAAATGGGACT------TTCTATCAGGAAGCCTCATGGGATGTGAAAGTAACTGCATCAGGCTCTACTCCTTTCATTTCC---ATCAAAGGTTATAAAAAATTTGGGATCCGAGCTGTTTGTCAGTCCAGCGGCTGG---------------------------------TACCCAAAACCTGAGATGCTGTGGAAAGATGCCAGTGGGAAACGTCTTTCTTCTTTTTTGGAAAGAAATGTCCAGAAGGACAATGGGCTCTTTGAAGTGCAGAATGAGATCATTGTCACTCAAAGTTCAGATCAGAACTTAACCTGTGTGGTCAGGAACATTCTTCTTCACAAACAAAAGGAATCAGCCATTCATATAGCAAATGACTTTTTCCCCAAAACGCCTGCCTGGATAGTGGGTCTGTGTATATCCCTGATGGTTTCAATTGGTTTGGCTCTCTTTATTCTTTACTTGTTTAAAATAAACAGAAAACTTGCAGCAGAACTTGGGTGG------------------------------------------------------------------------------------------------------------------------------------------------------------------------------------------------------------------------------------------------------------------------------AGAAACATGGTGGTGCCAGTA---GAGAAAGCAAACATAACCCTGGATCCAGACACAGCGAACCATGACCTTATCATCTCTGCGGACCAGAAGAACGTAATCCAA---GGGTTCATGTGGCACAGACTGCCTGACAACCCCAAGAGATTTGACGTGGAACGCTGCGTGTTGGGAAGCGAAGGGTTTTCTTCAGGGAGACATTACTGGGAAGTGGAGGTGGGGGAGGAAGGCTATTGGGCAGTGGGGGTGGCCAGAGACTCTGTGAAAAGGAAAGGGAGGCTC---------ATCCTTGACCCCAAAGAAGGAATCTGGGCTGTAGAAAAGTGT---------CCAGTCCAATACCAAGCACTCACT---GTCCCTGAGACGCCTCTGCCTTTGCAGAAGAAACCCCGGAAGCTTGGGATTTATTTGGACTGTGAGCTGAAGCTGGTGGCATTTCATGATGTCAATCACAAGGCCCCCATCTTCACATTCCCCATAGCTCCTCTCTGTGGGGAGAGGATCTTCCCTTTCCTTCATGTTGGA---GTAGGGTGTTGGCTTACAATT------------------------TGCCCCTGA------------------------------------------

>XM_060239261.1

---------------------------------------------------------------------------------------------------------------------------------------------------------------------------------------------------------------------------------------------------------------------------------------------------------------------------------------------------------------------------------------------------------------------------------------------------------------------------------------------------------------------------------------------------------------------------------------------------------------------------------------------------------------------------------------------------------------------------------------------------------------------------------------------------------------------------------------------------------------------ATGCCTCAATCTACGGACTCACATCTTAGCTTCTGGAACGTC---------------------------------------------------------------------------------------------------------------------------------------------TCATTTACAAACTGTGGCCTTCGGACATTCAAAGCGGTCTTCATCATCCTGTGGCTATCATTATGTCTTCCTGCTGAGTGTGAAGGCGCCAAAAAGGTCTCAGCATCCATG---ATTAAACAGTGGAGGAAAAACTAC------------------------------------------------------------------------------------------------------------------------------------------------------------------------------------------------------------------------------------------------------------------------------------------------------------------------------------------------------------------------------------------------------------------------------------------------------------------------------------------------------------------------------------------------------------------------------------------------------------------------------------------------------------------------------------------------------------------------------------------------------------------------------------------------------------AAAACCAAAGTGACTTTTGATCCCAACACAGCATATCCATGGTTCATCGTATCTGCAAACAAAACTTACGTGGAATCA---GGAGACAGACGCCAAAATGTGCCAGACAATCCTGAACGATTCAATAACACTCCTGCTTTGCTGGGCCACCCTGGATTTACATCCGGGAAGCTTTACTGGGAGGTGGAGTACGGAGACCAGAGGGAGTGGGCAGTGGGGGTGGCCCTCGAGCGTGTGGACAGGAAGGTCTATCTC---------AGGCTTAGTCCAGAGGAGGGCATTTGGCAAAAGGGCCTCTGGTGGCTTCGAGCCCTGGAAAATCGTTCCCATGCA------------------------CTTCCCGTACGCCCTGGGATCATTGGGATATTTTTGAATTACAAAGCAGGTACAGTGGCCTTTTACATACAGGGGAAA------------CTGATTCTGAAAAGAGCCTCTTTCAACAGGGAGAAAGTGTTTCCTTTCTTCTATCTGGGT---GGAGGTGTTCATCTCAAATTA------------------------ATATAA---------------------------------------------

>XM_060239250.1

------------------------------------------------------------------------------------------------------------------------------------------------------------ATGGCGGCGGCAGGGCCCATAGGTGGTGGCCTCCTGAACCCCGTGGCCACCCTCCACGAGGAGGCTGTCTGTGCGATCTGCCTGGACTACTTCACGGACCCGGTGTCCATC---GGATGCGGCCACAACTTCTGCCGCGTGTGCATCACTCAGTTGTGGGGTGCC------------GCTGCCGAGGACAGCCCAGATGAGGAAGATGAAGGCGAAGATGATGGA---------GGAGCGTCTGGCTCCGATGGTGACTACAGGCCTGGTTCT---GAGGAGGCGGTGGATGATGGAGGGATAGACGACTTGGGACCGGAGGAAGATGGT---GATGATGAGGATGATTTAGAG------GAGTACCTGGATGGTGGGGACATGGTTGAGGAAGAAGACGACGACGATGCATCTCGGGATGGGTAT------GATGAAGATGATGAGATCGCGTGGGATGAAGAAGTCAACGGAGACTTTTGGGACCAGGATCCTGGGTCGGATGAGACGTGGGAGGATGCCATGGGGGGTGACCTGTTCTTTGATAATTACAATGAGGAGGAGGAAGTGATGGAGGATGAAGCAGTGGAGGACGAG------------------GCAGACTCCCCTCTTACTCCACCACCGCGCCAGTCTTTCACCTGCCCACAGTGCCGCAAAACCTTTCCTCACCGCAACTTCCGTCCCAACCTTCAGCTGGCGAACATGGTGCAGATCATCCGCCAGATGCATCCTCAGCCCTTCAAGAGCACAACCCCTCCAGCTGAGGTGGTGGAGTTGGCGGCGGCAGGGGATGCGTTGGGAGTGTCATCGGCCTTGGCAGCGGGCAGAGGTGTGCGAGGTGAGCGCGTCTTGTGTGAGAAGCACCAGGAGCCCCTCAAGCTTTTCTGTGAGGAGGATGAGGAGCCGATCTGTGTGGTGTGCATAAAATCGCGCAGCCATAAGCACCATAATTTCGTGCCTCTTGAGGAAATTGTGGAAGAATACAAGGCCAAGCTCCAGGGTCACTTGGATCCACTGAAGAAGAAACTGGAGACAGTTCTGAAG---CAGAAGTTGAGTGAACAGGAGAAAGTATCTGAACTGAGAAACAAGATGAAGCTG---------------------------------GATATGAAAGAATTTGAGTCGGACTTTGAGCGGCTACACCAGTTCCTTGTGGGGGAGCAGGCACAGCTGCTCCACCAGCTAGAGGATCGCTACAACATTCTACTAGCCTGCCATAATAGTAATGTTACCCACCTGGAGGAGCAGAGTGCCACGCTCAGCCACCTCATTGCGGAAGCTGAAGACAAGAGCAAGCAGGATGGTTTGCAGCTGCTCAAGGATTTCAAAGGC---ACATTAGTCAGGTGTGAGAACATAAAATTTCAGGATCCTGAGAGGGTG---CCAGTGGACACAGGAAAGAAATATCAGAACTATTTCCTTCTAGACACCTTGATGAGGAAG---------------------------------------------------------------------------------------------------------------------------------------------------------------------------------------------------------------------------------------------------------------------ATGGAGAAAGTCTTCTGCAAAGTCCCTCAAGTTGACTTGACCCTGGACCCAGAGACAGCACATCCAAGCCTCACCCTTTCCTCAGACTGCCACAGTGTCCAACTC---GGGGAGCATTGGCGAGATGTGCCCAACACCTCCAAACGCTTCAACTCAGACTATTGCGTGCTGGCTTTGAAGGGTTTCACCTGGGGCCGCCACTGTTGGGAAGTGGAGGTGAGTGGTGGGTGGGGTTGGGCCGTGGGGGTGACCCGCAAAATGGCCAGACGCAAAGAGAAGTCC---AGCAGAGGACCCCACCTGAAGCGGGAGATCTGGTGCGTGGGGACTAAT---------GGGAAGAAATACCAGGCCCTGACGGCCACTGAGGAAACGTCCCTCTCTCCATCGGAGAAGCTCCGGCGCTTTTGTGTCTACCTGGACTATGAGCGGGGCCAGCTGAGCTTCTACAATGCCGAGAGCATGACCCACATCCACACTTTCAATGTTTCTTTC------CATGAGCGCATTTTCCCTTTCTTTTGCGTCCTCTCCAAGGGGCCCTGCATCAAACTT------------------------TGCAATTGA------------------------------------------

>XM_060239249.1

------------------------------------------------------------------------------------------------------------------------------------------------------------ATGGCGGCGGCAGGGCCCATAGGTGGTGGCCTCCTGAACCCCGTGGCCACCCTCCACGAGGAGGCTGTCTGTGCGATCTGCCTGGACTACTTCACGGACCCGGTGTCCATC---GGATGCGGCCACAACTTCTGCCGCGTGTGCATCACTCAGTTGTGGGGTGCC------------GCTGCCGAGGACAGCCCAGATGAGGAAGATGAAGGCGAAGATGATGGA---------GGAGCGTCTGGCTCCGATGGTGACTACAGGCCTGGTTCT---GAGGAGGCGGTGGATGATGGAGGGATAGACGACTTGGGACCGGAGGAAGATGGT---GATGATGAGGATGATTTAGAG------GAGTACCTGGATGGTGGGGACATGGTTGAGGAAGAAGACGACGACGATGCATCTCGGGATGGGTAT------GATGAAGATGATGAGATCGCGTGGGATGAAGAAGTCAACGGAGACTTTTGGGACCAGGATCCTGGGTCGGATGAGACGTGGGAGGATGCCATGGGGGGTGACCTGTTCTTTGATAATTACAATGAGGAGGAGGAAGTGATGGAGGATGAAGCAGTGGAGGACGAG------------------GCAGACTCCCCTCTTACTCCACCACCGCGCCAGTCTTTCACCTGCCCACAGTGCCGCAAAACCTTTCCTCACCGCAACTTCCGTCCCAACCTTCAGCTGGCGAACATGGTGCAGATCATCCGCCAGATGCATCCTCAGCCCTTCAAGAGCACAACCCCTCCAGCTGAGGTGGTGGAGTTGGCGGCGGCAGGGGATGCGTTGGGAGTGTCATCGGCCTTGGCAGCGGGCAGAGGTGTGCGAGGTGAGCGCGTCTTGTGTGAGAAGCACCAGGAGCCCCTCAAGCTTTTCTGTGAGGCGGATGAGGAGCCGATCTGTGTGGTGTGCAGGGAATCGCACAGCCATAAGCACCATAATGTCGTGCCTCTTGAGGAAATTGTGGAAGAATACAAGGCTAAGCTCCAGAGTCACTTGGATCCACTGAAGAAGAAACTGGAGACAGTTCTGAAG---CAGAAGTCGAGTGAACAGGAGAAAGTAGCTGAACTCAGAAACAAGATGAAGCTG---------------------------------GATATGAAAGAGTTTGAGTCAGACTTTGAGCGGCTACACCAGTTCCTTGTGGGGGAGCAGGCACAGCTGCTGCATCAGCTAGAGGATCGTTACGACATTCTACTAGCCCGCCAGAATAGTAATGTTACTCACCTGGAGGAGCAGTGTGCTGCGCTCAGCCGCCTCATTGCAGAAGCTGAAGACAAGAGCAAGCAGGATGGTCTGCAGCTGCTCAAGGATTTCAAAGGC---ACCTTAGTCAGGTGTGAGAACATAAAATTTCAGGATCCTGAGATGGTG---CCAGTGGACACAGGAAAGAAATATCGGAACTATTTCCTTGTGGACACCCTGATGAGGAAG---------------------------------------------------------------------------------------------------------------------------------------------------------------------------------------------------------------------------------------------------------------------ATGGAGAAAGTCTTCTGCAAAGTCCCTCAAGTCGATTTGACCCTGGACCCGGAGACAGCCCATCCACGCCTCACCCTTTCCTCAGACTGCCGCAGTGTCCGACTC---GGGGAACATTGGCGAGATGTGCCTGACACTCCCAAACGCTTCAACTCCGACTACTGCGTGCTGGCTTTGAAGGGCTTCACCTGGGGCCGCCACTGTTGGGAAGTGGAGGTGAGTGGTGGGTGGGGTTGGGCCGTGGGGGCGGCCCGTGAAACAGCCAGACGCAAAGAGAAGTCC---AGCAGAGGACCCCACCTGAAGCGGGAGATCTGGTGCGTGGGGACCAAT---------GGGAAGAAATACCAGGCTCTGACAACCACCGAGGAAACGTCCCTCTCTCCATCAGAGAAGCTCCGGCGCTTTTGTGTCTACCTGGACTATGAGCGGGGCCAGCTGAGCTTCTACAATGCCGAGAGCATGACCCACATCCACACTTTCAATGCCTCTTTC------CATGAGCGCATTTTCCCTTTCTTCCGCATCCTTTCCAAGGGCACCCGCATCAAACTC------------------------TGCAATTGA------------------------------------------

>XM_060239255.1

------------------------------------------------------------------------------------------------------------------------------------------------------------------------------------ATGGCTGCTGCAAATCCAGCAAAAACCCTTCTGGAGGAAGCTACTTGTTCCATCTGTCTGGATTTATTTCAGGATCCTGTGATGATAATAGAATGTGGGCACAATTTCTGCCGAAAGTGCATCACCAAATATTGG------------------------------------------------------------------------------------------------------------------------------------------------------------------------------------------------------------------------------------------------------------------------------------------------------------------------------------------------------------------------------------------------------------------------------------------------------------------------------------------AAAGGATCTGATTGCAGTGCCTGCTGCCCGGAATGCAGATGCTTGTTTTCCTGGACAAATCTCAAACCAAACAGACAACTGGGGAATATGGTAGAAACGGCTAAACAACTCGGTTTG---------------------------------------------------------------------------------------CAGCTAGAGAACAGCTCGGAAGGAGAGAGAATGTGCAAGGAACACCAGAAGCCCCTGACTCTCTTCTGCAAAACTGATGAGACTCTCATTTGCCAGGTCTGTGATAGATCCATAGCTCACCGGAGTCACACAGTTGTTCACACAGAAGAAGCTGCTGCAGATTACAAGGAGATGCTGCTAAAGCATCTTAGAAATCTGAAGGAAGGAAGAAACAAGATTCAGTCA---GCTAAATCAAATGGAGAAAAGCCAAGCCAGGAATTGTTGAAAGAGACACAAACC---------------------------------AAGATACAAGAGATTGTATCTGAATTCCAACAACAGAGGCAGTTTTTGGAACAACAAGAGCAACTCCACTTATCCAAGCTGAGAGACCTGGAAAAGAAGATTGAGAAGAAAAGAAATGATTATGCCAGCAAGTGTGTGCATGAAATATCTTGCATCACTGCTCTGATTGGTGATGTGGAAAAGAAAGATAAGCAGCCAGCAAATGAATTCCTGCACGATATTGGAAAC---ATATTGGACAGATGTAAGAATGGGAAATTCTATTATCCAGCTGTACCGGATACTTCCGATATGAAGAGAAGGCTTAATCAGCTCTGCCAAGAGAGCACATCTCTGCAAAGTTCTTTTAAGAATTTCCGA---------------------------------------------------------------------------------------------------------------------------------------------------------------------------------------------------------------------------------------------------GAGAACATGTTGAAATCAAAATGGATCAAAGAAAATGTGTTGCTGGATCCAGAAACATCCCATCCCCGATATATTGTGTCTGCTGATCGGAAGACTGTAAAATGG---GGGAGTGCCCGTCAGCAGTTTCCCTACAACAAAAAGAGGTTTCAGTGTGTTCGCTGTGTTCTTGGCTCTGAGGGGTTCACTTCAGGAAAACATTATTGGACTGTGGATGTAGGAGATGGAGATTATTGGGCTGTGGGAGTGGTCAGAGAATCTGTGGAGCGGGAGGAGGAAATT---------CCTCTTGAACCTGATGAGGGAATCTGGGCAATTGGGCTTTAT---------GACAATCAGTACAAAGCTCTGACC---TCACCTCCTACTGTGCTAGAGGTAGAGGATGAGCCAACAGAGATCCAGGTATCTTTGAATTATGAAGCAGGGACAGTTAATTTTTATGATGCAGAAGATAGGACCCGTCTGTTCACTTTTCAGTCAGTTGATTTTGAAGGGGAGGAAGTTTTCCCTTTTTTCCGGATTGTGAATTCATCAACTGTCCTGACGATG------------------------TCCTATTAA------------------------------------------

>XM_060239253.1

------------------------------------------------------------------------------------------------------------------------------------------------------------------ATGTCCACAGAAAACATCATGGCTGCTGCAAATCCAGCAAAAACCCTTCTGGAGGAAGCTACTTGTTCCATCTGTCTGGATTTCTTTCAGGAACCTGTGATGATCATAGAATGTGGACACAATTTCTGCCGAAACTGCATCATCAGATATTGT------------------------------------------------------------------------------------------------------------------------------------------------------------------------------------------------------------------------------------------------------------------------------------------------------------------------------------------------------------------------------------------------------------------------------------------------------------------------------------------AAAGAATCCGATTGCAGGGCCTGCTGCCCGGAATGCAGATGCTTGTTTTCCTGGACAAATCTAAAATCAAACAGACAACTGGGTAATATTGTAGAAACAGCTAAAGAACTCAGTTTG---------------------------------------------------------------------------------------CAGCTGAAGAACAGCTGGAAAAGCGAGAGAATATGCAAGGAACACCAGACGCCCCTGACTCTCTTCTGCAAAACTGACAGGACCCTCATTTGCGATGTCTGTGATGGATTCGAAGCACATCAGGGACATGCAGTTGTTCACAAAGAAGCTGCTACTGCAGATTACAAGGAGACACTGCTGAAGCATCTTAAAAATCTGAAGGAAGAAAGAAACAAATTTCAGACA---GCTAGATCAAATGGAGAAAAGCCAAGCCAGGAGTTATTGCAAGAGACACAAGCC---------------------------------AAGAGACAAGAGATTGTGTCTGAATTCCAAAAACGGCGGCAGGTTTTGGAACAACAAGAGCAACTCCAGCTGTCCCAGCTGAGAGAGCTTGAAAAGGAGATTGAGAGGAAAAGAGATGACTATGCCGACAAGTGTGTGTACGAAATATCTTGTATCAACGCTCTGATTGGTGACGTGGAAAGGAAGAATAAGCAGCCAGCAAATGAATTCCTGCAGGATATTGGAAGC---ATGTTGGACAGATGTAAGAAGGGGAAATTCTGTTATCCAGTTGTACCGGATACTTCAGCTATGAAGGGAAGGCTTAATCAGCTCCCTCAAGAGAGCACATCTCTGCAAAGTTCTTTGAAGAATTTCCAA---------------------------------------------------------------------------------------------------------------------------------------------------------------------------------------------------------------------------------------------------GCGAACACGTCAAAATCGAAATGGATCAAAGAAAATGTGTTGCTGGATCCAGAAACAGCCCATCCCCGATATGTTGTGTCTGAAGATCAGAAGACTGTAAGATGG---GGGAGTTTCCGACAGCAGTTTCCCTACGACCCAAAGAGATTTGAATGTATTCGCTGTGTTCTTGGCTCTGAGGGGTTCACTTCAGGAAAACATTATTGGACTGTGGATGTTGGAGATGGAGATTATTGGGCTGTTGGAGTGGCCAGAGAAAATGTGGAGCGGGAGGAGGAAATT---------CCTCTTGAACCTGATGAGGGAATCTGGGCAATTGGGCTTTAT---------GACAATCAGTACAAAGCTCTGACC---TCACCTCCTACTGTGCTAGAGGTAGAGGATGAGCCAACAGAGATCCAGGTATCTTTGAATTATGAAGCAGGGACAGTTAATTTTTATGATGCTGAAGATAAGACCCGTCTGTTCACTTTTCAGTCAGTTGATTTTGAAGGGGAGGAAGTTTTCCCTTTTTTCCGGATTGTGAATTCATCAACTGTCCTGACATTG------------------------TCCTATTAA------------------------------------------

>XM_060238765.1

---------------------------------------------------------------------------------------------------------------------------------------------------------------------------------ATGGCCACGGGCCGGGATCCAGGCGCAGCTGTCCGAGATGAAACGACCTGCTCCATCTGCTTGGATCATTTCCAAGATCCACTGATGATCGTAGACTGCGGGCACGATTTCTGCCGCGCCTGCATCACCCGGTACAGCGGCGGA------------------------------------------------------------------------------------------------------------------------------------------------------------------------------------------------------------------------------------------------------------------------------------------------------------------------------------------------------------------------------------------------------------------------------------------------------------------------------------GGATCGGATCTCAGCGTCTTGTCTTGTCCCGAGTGCAGAAAACCCTTCTCTTGGGGAGATCTCCGGGCAAACCGGCGCTTGGGGAATGTGGCAGAGCTGATCCAGCAGTTCCCTCTG------------------------------------------------------------------------------------GAGCGGTTGAGTGAGGCCGAAGGAGCGCAGAAGGTGTGTCGGGAGCACAAAGAAGCCTTCCAGTTCTTCTGCCGAGAGGATCGAGCTTTTATCTGCTCGATCTGCAAAGATTCCGAAGCTCACAAGACACACGTGGCGCTTCCCATAGACCAGGCGGTCCAGAATTACCAGGACCAAATACAAAATTATGTGACATTACTGAAAAAGGAGAAAGATGAAATTCTAAAG---AAGATA---AATAATGAGAAAGCAATTAAGGAAGTGATAAGGAGTATCATATAT---------------------------------ATTAGAGAGCAGAAAGAACATGGATTTCCAAAG------------------GAAGAAGTCCACACTCTGTTGGCGCTGCTGAGAGATGTAGAAGAAAAAACTGTGCAGATGGCAGATGAAAATGCAAGTGATACCCTGGAGCACAGTGCCCATCTCACCGAGCTGATCAGAGAGATGGGGGCCAAGTGCCTGCAGCCAGCCTTTGAATTCTTGCAG---------------------------------------------------------------------------------------------------------------------------------------------------------------------------------------------------------------------------------------------------------------------------------------------------------------------------------------------------------------------------------------------------------------------------------------------------------------------------------------------------------------------------------------------------------------------------------------------------------------------------------------------------------------------------------------------------------ATGCGAGAGAGAAATGGCTAG------------------------------------------------------------------------------------------------------------------------------------------------------------------------------------------------------------------------------------------------------------------------------------------------------------------------------------

>XM_020777528.1

---------------------------------------------------------------------------------------------------------------------------------------------------------------------------------ATGGCTGTCGGCGGGGACCCCTCGCTGACTCTCCACGATGAGGCGACCTGCTCCATCTGCTTGGATTACTTCCAGGAGCCGATGATGATCGTCGAGTGCGGGCACAACTTCTGCCGAGCCTGCCTGGCCGACTACGAGGCGAAC------------------------------------------------------------------------------------------------------------------------------------------------------------------------------------------------------------------------------------------------------------------------------------------------------------------------------------------------------------------------------------------------------------------------------------------------------------------------------------GAAGCGGGTCGCGACGGCCTCCAGTGTCCCCAGTGCCGGGGAGCGTTTGCTCACCACCACCTCCGGCCCAACCGGCACCTGGGGAATATGGTCGACGTGATCCGCCGGCTGAACCGG------CAGCGG---------------------------------------------------------------CTGGGCAAGGCCGGAGAGGAGGCAAGCCGGGCGGCCTCGCAGGTGTGCGAGAAGCACCAGGAGGCCGTCCAGCTCTTTTGCCAAGAGGACCGAGCCTTCCTCTGCCTCGTCTGCAGGGAATCCCGGGCGCACAAAGCCCACGCGGTCCTCCCCGTAGACGAGGCTGCCCAGGAGTACAAGGACCAGCTCCAGGGCTTCTTGACAACATTGACAACAGAGAGAGGGGAAATTCTGAAG---GAAGAATGGTGCAATCAGGATGAAGTTCAACAAATGAAGAGAGATATTCAAATC---------------------------------ATCCAGAATGGCATTCTAACCCTATTTCCGGAG---------------------GCGAAAGACCAGACCCAGCTGGCCCTGCTTGATGTGTGTGTTGAAAATATGAAAGCGATAGACAGCATTGCACATGACTTCTCAGAGCGCTGTTTCCTTCTCACCAAGCTGATCAGCGAGATTGAGGACAGGCTCCAGCAGTCCACCTTTAAATGCTTACAAAACATTGGGGAC---TTACTGAGCAGGTGTAAGAGAGAGGTATCTGAAACAGCC------------AAAGCTGATCTGAAAAGGAGAGCTGAATCGGTCATTGAAAAACTGGCACATGTTTCGACCAAATTGAATGAATTGGCA---------------------------------------------------------------------------------------GGAAGCAGCCGTGGCCAATCACCGCATCCCCGTGGCCGGTCCAGGGAGACAGCTGCCAGTCCCTCTGTCAACACCAGCCAGCATGGTCAGGCTTTTGCAACAGTCCCCAGTGTGTTCTGTCCTTATTCAGTCGTTGTTGCTCATCCTCAGCAACAGCAAAGAGGGAACAGGAAAAAATGGAAAAAAGAAAATGTGCTGCTGGATCCTGAAACAGCCCACCCTCGATACATCATATCTGATGACTGCAAAAGAGTAGATTGG---GGAGATGTCCGTCAGGATTTCTTTTACAACCCAAAGAGGTTTCAGTATGCTCGTTGTGTGCTTGGAACAAGGGGATTCAAGTCAGGAAAACATTATTGGACAATAGATGTACAGGATGGACACCACTGGGCTGTGGGTGTGGCACGAGAGTCTGTGGAGAGGGATGTGGTGCTC---------TCTTTTGAACCCGATGAAGGGATCTGGGCCATGGCACGTTAT---------TGTGATCAGTACAAAGCCATGACT---TCACCTCCTACCATTCTGGAACTAGATGATGAGCCAACAGAGATTCAGGTTTCTCTGAACTGTGACGCTGGCACTGTAACTTTTTATGATGCTGAAGACTTAACCCGCCTCTATAAATTTGAGTGTGTTGATTTTGAAGGAGAGAAGGTTTTTCCTTTTTTTCGGATTTCTGATTCGTCTACTTCCCTCTCGATC------------------------ACTTCCTAA------------------------------------------

>XM_020777535.1

------------------------------------------------------------------------------------------------------------------------------------------------------------------------------------ATGGCTGCTCCCAATCCAGAAAAATCCCTCCAGGACGAAGCCACTTGTTCCATCTGCCTGGATTATTTTCAGAACCCGACGATGATCATAGATTGTGGGCACAACTTCTGCGAAGCCTGCATCGGCCAGTGTTGCGAA---------------------------------------------------------------------------------------------------------------------------------------------------------------------------------------------------------------------------------------------------------------------------------------------------------------------------------------------------------------------------------------------------------------------------------------------------------------------------------------GGGCCCGTTTTGAACCAGTTTTCTTGCCCTCAATGCAGGAAGCCCTTTTTTTGGCAAAACGTCCGGCCCAACAGGCACCTGTGGAATATTGTAGAACTTGCCAAGCAGTTCAGCCTG---------------------------------------------------------------------------------------CGGCGGGCAAATGAAGCGGGAGAGCGGAAGTTGTGCAAGAAGCACCAGGAACCCCTCAAACTCTTCTGCGAGCAGGATCAGACCTCCATCTGCCTGGTTTGTGATCGATCAAGGGTGCATAAAAACCACACTGTTGTTCCTGTCGAAGAGGCTGCGCAGGTTTACCAGGAGAAAATACACAACTGTTTGAAGACTCTGAAAGAAGAAAGAGGCAAGATTTTGTCA---TTCAAATCAGAGTGGTCCAAGCCAAGTGAAGATTTGCTGAAGGAAACAGAAGTT---------------------------------GAAAGGCAGAAGCTGGTGTCAGAATTCCAGCAGCTGCGCCAGGTTCTTGAGAAGGAGGAGCAAACCCTTTTGACCCAACTTGAAAAGCTGCAGGCAGATGCGAAGAAGAAGATGGAGGAAGGCGAAGCTAAATATTCTGCAGAGATTTCCCATCTCACTGCCCTCATTGGTGAGATGGAGGAGAAGTGCCAGGAGCCTCCAGGTGGGCTGCTGCAGGATATTGGATGC---ATGTTCAAAAGGTGCCAGAGAGGCAAGTTTCAGTCTCCTGCTGTACCTGATTCTTCAGAGCTAAAGAAAAGGCTTCAAATATTCTCCAGAGACAGTACATCACTACAGGGGCAACTGAAAAAAATCAGA---------------------------------------------------------------------------------------------------------------------------------------------------------------------------------------------------------------------------------------------------GAGACACTAACTGAACCAAAGTGGATAAAAGAGGATATGACATTGGACGTCGCAACAGCTCATCCACGATTTGTTGTGTTTGAGGATCGGAAAACTGTCGCATGG---GGCCCTGTTCGTGAGGAGCTGCCTTACGATGCCAGAAGATTTGACCCTTCCCGCTGTGTGCTTGGTTCGCGAGGGTTCACCTCAGGGAAGCATCACTGGACAGTGGATGTGGACCAGGGAACTTTCTGGGCCGTGGGTGTGGCCCGAGAGTCTGTGAAGAGGAAAGGACAATTC---------ACTATAATCCCCGCAGAGGGAATCTGGGCTGTTGGACTGTCC---------AGTGGCCACTACAAAGCTCTGACT---TCACCTCCTGTCATTCTGAGCCATTGCAAACCCATGAGAAAGATCCGGGTTTGTTTAGACTGTGAAGGAAAAACGGTGATGTTTTTTGATGCTGAAGAAAAGGAGGCCCCTGTTCTGTTCTCATTCCCAATTTCTGGATCAGGAAAATTTTTTCCTTTTTTCCGAGTCGGAGACATGAACACCGTCCTTCGCCTG------------------------TGTTAA---------------------------------------------

>XM_020777526.1

---------------------------------------------------------------------------------------------------------------------------------------ATGGCCGCCGCCGCCGCCGCCTCCGTGCCTGCAGGGAGTGTCTATGGGGGCCCCCTGAACCCGGTGACGACCCTCCACGAGGAGGCGGTCTGCGCCATCTGCTTAGACTATTTCATCGACCCGGTGTCGATC---GGATGTGGCCACAACTTTTGCCGCGTCTGCATCACGCAGTTGTGGGGCGCAAGCGAAGGTCAAGGAGGGGAGGAGGAGGAGGAGGAGGAAGCCGAAGCCGCGGAGGAAGGAGAGGAGGTGGGGACCTCTGGTTCGGATGGAGGCTATGGGCCTGCTTCTGAGGATGGAGGTGGGGAGGAGGGAGGCATAGACGACCTCCCACGGGAAGAAGACGCCGAAGATGAAGAGGATGATCTCGAGGATGATGACTACTTGGACGATGGGGGTATGGAAGAGGAGGAGGAGGAAGAGGATGTCTCTAGGCACGAGTAT------GATGGTGACGATGACGATGTGTGGGATGAAGGAGATGATAGAGACTACTGGGATGAGGATCCTGGATCGGATGAGATGTGGGATGATGCCATGGGGAGTGACTTGTTCTTTGATAACTATGGAGAGGAGGAGGAGGTAATGGATGAGGAGGAAGAGACGGAGGAAGATGCAGAGTATAATGTCCCAACGCCACCGCCGCCTCCCCCTGCGCGCCAGTCATTTACTTGCCCCCAGTGCCGTAAAAACTTTCCCCATCGAAACTTTCGACCCAACCTTCAGCTGGCCAATATGGTGCAGATCATCCGCCAGATGCATCCACAACCCTTCAAGCCTGCGACTGCATCACCGGAGACAGTGGAATTGGAAGCGGCAGGGGATTCATCTGGGGCACCATCATCTTCAGTAGCAGGCCGAGTTCCAAGCGCTGAACGAACGCTATGTGAGAAACACCAGGAACCACTCAAACTTTTCTGTGAGACAGATGAGGAGCCCATCTGTGTGGTCTGCAGAGAATCTCGAAACCACAAACACCACAGTGTCATGCCACTAGAAGAAGTTGTACAAGAATATAAGGCAAAGCTCCAAGGCCACTTGGATCCACTGAAGAGGAAACTGGAGACTGTTCTGAAA---CAGAAGTCGCGTGAAGAGGAAAAGATAGCAGAACTCAAGAACAAGATGAAACTT---------------------------------GATATGAAAGAGTTTGAATCTGATTTTGAGCGGCTGCACCAGTTTCTTGTCGGGGAGCAGGCATTGTTGTTGCACCAGCTGGAAGACCGCTACGAGGTGCTGCTAGGCACACAGAACAGCAACATCACTCATCTTGAGGAGCAAGGTGCTGCACTCAGCCGCCTCATTGCTGAAGCTGAAGACAAGAGCAAGCAGGATGGTTTGCAGCTGCTCAAGGACTACAAAGGC---ACTTTAGTCAGGTGTGAAAATATTAAGTTCCAAGATCCAGAGATAGCA---CCTGTGGACACAGGAAAGAAATACCGGAACTACTTCCTCGTGGATGTCTTAATGAGAAAG---------------------------------------------------------------------------------------------------------------------------------------------------------------------------------------------------------------------------------------------------------------------ATGGAGAAGGTCTTCTGCAAAGCCCCTCAAGCTGACTTGACCCTGGACCCAGAAACAGCCCATCCACGACTCATCCTTTCCTCAGACTATCGGAGTGCCCGACTC---GGAGATCGCTGGCGGGAGCTTCCTGATAACCCAAAACGCTTCGACTCCGACTATTGTGTGTTGGCTCTTCAGGGTTTCAACCAAGGCCGCCATTACTGGGAAGTGGAGGTTGGAGGCCGACGGGGGTGGGCAGTGGGGGCTGCCCGCGAATCAGCTCGACGTAAAGAAAAGGCTAACGCTGGAGGTCCCCACCAGAAACGGGAGATCTGGTGCGTGGGGACCAAT---------GGCAAGAAGTACCAGGCACTGACCACCACCGAGCAGACATCTCTGCCTCCGTGTGAGAAACTCCGGCGTTTTTGTGTCTACCTGGACTATGAACGGGGCCAGCTGGCCTTCTATAATGCTGAGAATATGGCCCACATCCACACTTTCAATGCCTCATTC------CGTGAGCGCATCTTTCCCTTCTTCCGTATTCTTTCTAAGGGGACCCGCATCAAAATC------------------------TGCAATTGA------------------------------------------

>XM_020777533.1

------------------------------------------------------------------------------------------------------------------------------------------------------------------------------------------------------------------------------------------------------------------------------------------------------------------------------------------------------------------------------------------------------------------------------------------------------------------------------------------------------------------------------------------------------------------------------------------------------------------------------------------------------------------------------------------------------------------------------------------------------------------------------------------------------------------------------------------------------------------------------------------------------------------------------------------------------------------------------------------------------------------------------------------------------------ATGCAGACAGGCAGTGTTCTACAGAAGTTCAGAGGACTCTGTTGTATTACGTGGCTGCTGCTGTGCCTTCTTGCTCAGTACGAAAATGGTGGAAAG---------GTCCTGGCTTTATCTTCATCAGCCAAACCCTAC------------------------------------------------------------------------------------------------------------------------------------------------------------------------------------------------------------------------------------------------------------------------------------------------------------------------------------------------------------------------------------------------------------------------------------------------------------------------------------------------------------------------------------------------------------------------------------------------------------------------------------------------------------------------------------------------------------------------------------------------------------------------------------------------------------AAAACCAGTGTGACTTTTGATCCAAAGACGGCGCACCCAAATCTTGTGGTGTCTCAAGACAAGAAGACGGTCACATGG---GTCCAGGAAGCCCAAAGTGTGCCTGACAACCCAGAGAGATTCAACAGCACCCCTTGCTTGCTGGGTTCCCCAGGGTTCACATCTGGGAAACATTACTGGGAAGTGGAGTATGGGAACCAGAGAGAGCTCGCAGCGGGGGTGGCCCGGAAGTCTGTGAAGAGGAAGGACCACCTC---------CGACTTACTCCAGAGGAAGGGATTTGGCAAGTGGGTCGCTGGTGGCTTCGGCGAGGGGGGCCTGAAAACCAAAGT---------------------------------GACTCAGGAAAGATTGGCGTCCTTCTGGATTATGGAAGGAATCAGGTGACTTTTTATATGGATAATAAAGTCACTGTAGTC------------CGGGCCTCATTCAATGGGGAGGAGGTTTTCCCCTTCTGCTACGTGGGG---TCCACTGTTTCCCTCCGTTTA------------------------AATCCCTGA------------------------------------------

>XM_053295997.1

------------------------------------------------------------------------------------------------------------------------------------------------------------------------------------ATGGCGACTGCAAGCCCAGCAAAATGCCTTCAAGATGAAGCTACTTGTTCAATCTGTCTAGACTACTTCCAGGATCCTGTGATGATCAGCAATTGTGGGCATAATTTCTGCCGAACCTGCATCATCGAGTAT---------------------------------------------------------------------------------------------------------------------------------------------------------------------------------------------------------------------------------------------------------------------------------------------------------------------------------------------------------------------------------------------------------------------------------------------------------------------------------------------AAGAAGTCTGATATTAATATCTGCTGCCCAGAGTGCAGACGATCTTTTTCCCGGGAACATCTTAAACCCAATAGACCCCTGGGGAGAATGGCAGAGGCAGCTAAACAACTCAGTTTG------------------------------------------------------------------------------------------CAGATCAACAGCCCAGCAAAGAAGAGAATGTGTCTGAAACATAAGCAGCCTCTGATCCGCTTCTGCAAAGAAGATGAGACCCTCCTTTGCTGCCTCTGTGAGGAATCCAAGGCTCACCGAAATCACACGGTTGTTTACGCAGAAGCTGCTGCTGCGGATTACAAGATGAAACTTCAGAAACGTCTAGAGAACCTGAAGAAAGAAAAAAACAAAATTCTGTCA---GTCAAATCAAATGGAGAAAAGCCAAGCCAGGAACTACTGAAGGAAATGGATGAC---------------------------------AAAAAGTCGTCGATGGTATCTGAATTCCAACAGATAGTGCAGTTTTTGAAGGAAAAAGAGCAATTCCAGTTGAGCAGGTTGGGAGAACTGGAAGCGGAGACTGAGAGGACAAGGAAAGATTATGGGAATAAATTTGATGATGAAATATCCTACATCACCAATCTGATTGGTGAGATAGAAAAGAAGAATGGCCAACCACCCAGTGAATTTCTGGAGGATATTGGAAGC---ATATTGGATAGATGTGAAAAGGAGAAATTCAAGTTTCCACCTCCACCTGATACTTCAGATATAAAGAAAAGGCTTGAGCAGTTCTCTCAACAAAGTACAGGCGAGCAAAGTACTCTGAGAAAATTCAGAGCCCTCCAACAGCAGCTGCAGGAGGAAGAGAGGAGACAGCAAATGGGGAGGAACGTTGCTGCCTCTGCAGGCGCCAGGCGGGGGAGAAACCCAGCGCAGGAAATGCAGGAAGAACTCCAGCAAGTTGTCATCAGTCGCTTGAGAGCCCTCCAACAGCAGCTGCAGGAGGAAGAGAGGAGACAGCAAATGGGGAGGAACGTTGCTGCCTCTGCAGGCGCCAGGCGGGGGAGAAACCCAGTGCAGGATAACATGTTGAGACCAAGATGGATTAAAGAAAATGTGTATCTGGATCCAGAAACAGCTCATCCTCGGTACATTGTGTCTCAAGACGGGAAAAGTGTCGAATGG---GGAGAAATCCGTCAGGATTTTCCCTACAACCCAAAGCGATTTCAGTATGCTCGTTGTGTTCTTGGTTGTGAGGGATTCACCTCAGGGAAACATTATTGGACAGTGAGCGTAGGGGATGGAGACTATTGGGCTGTGGGAGTGGCCAGAGAGTCTGTGGAGAGGGGGGAGGAAATT---------GATTTTGATCCTGATGAGGGGATCTGGGCTTTGGGACGTTAC---------GATGATCAATATAAGGCTCTGACT---TCCCCTCCTACTCTTTTGGAGCTAGACTATGAGCCAACAGAGATCCAGGTTTCTCTGAATTATGAAGCAGGAAAAGTGCATTTTTATGATGGAGAAGATAAGACCCGCATCTATACTTTC---GATGCTGATTTTGAAGGGGAGGCAATTTTCCCTTTTTTCCGGATTGTTGATTCATCAACTTGCCTCCAGCTC------------------------TGTTCTTAA------------------------------------------

>XM_053287828.1

------------------------------------------------------------------------------------------------------------------------------------------------------------------------------------ATGGCTGCTCTGAATCCTGAAAAATCTCTCCAAGATGAAGCCACTTGTTCTATCTGTTTGGATTATTTTCAGAGTCCAGTGATGATCATAGATTGTGGGCACAATTTCTGCCGAGACTGCATTGGCCAGTGCTGTGAA---------------------------------------------------------------------------------------------------------------------------------------------------------------------------------------------------------------------------------------------------------------------------------------------------------------------------------------------------------------------------------------------------------------------------------------------------------------------------------------GGGTCTGTTACCAAAGTGCTTTCTTGCCCTCAGTGCAGAAAACCTTTTCTCTGGAAAAACATCCGGCCCAACAGGCATCTGTGGAATATTGTAGAACTGGCCAAACAGTTCAGCGCA---------------------------------------------------------------------------------------CGAAGGGCTAATGAAGCAGGAGATCAGCGGTTGTGCGAAAAACACCAAGAGCCCCTGAAACTCTTCTGCGAACAGGATCGAACTCCTATCTGTGTGGTGTGTGATCGATCTAAGGTGCATAAAAACCACAATATTATTCCCATAGAGGAGGCTGCTCAGGCGTACCAGGATAAAATACATAATTGTCTGCAGACTCTGAAAGAAGAGAGAGAAAAGATTTTATCG---CTTAAATCGGAGGTAGAGAAACCAAGCCAGGATCTACTGAAAGAAGCCAAAGCA---------------------------------GAGAGGCAGAAACTTGTGGCAGCATTTCAGGAGCTGCGTGAGCTTCTGGAGAAAGAAGAACACCTCCTGTTGGCCCAGCTGAAAAACCTAGAGACGGAAATTGAGAAGAAGAAGGAAGAAAATGCATCTAGATATTCTGAGGAGATTTCCCTTCTCGGTTCCTTAATTGGCGAGCTTGAGCAGAAGCGCCAGGAGCCGCCAAGTGGACTCCTTCAGGATATTGGAGGT---ATTTTGAAAAGGTGCAAGAAGGACAAATTTCAGCCTCCA---GTTCTGGATTCTTCAGAATTCAAGAGAAAGCTCAAAGAATTTTCCCAAAAAAGTGCATCGCTACAGGGTGACCTGAAGAAATTCAGA---------------------------------------------------------------------------------------------------------------------------------------------------------------------------------------------------------------------------------------------------GAGACTCTGATTGAACCAAAGTGGATAAAAGAGAACGTTCTGTTGGATCCATTGACGGCGCACCCTCGATTTGTGGTGTCCGATAATCAGAAAAGTGTTGCATGG---GGATCTTTCCGCATGGACTTGCCCTACAACGCCAAGAGGTTTGACCCTGCCCGCTGTGTGCTTGGCTCTCAAGGATTCTCCTCGGGAAAACATCATTGGACGGTAGATGTGGCCAATGGAACTTTCTGGGCTGTGGGAGTGGCCAGAGAGTCTGTAACGAGAAAAGGACAACTC---------AGTTTTATTCCAGAGGAAGGAATCTGGGCTGTGGGTTTTTAC---------AATGGTCAGTACAAAGCTCTAACT---TCACCACCTACCTTTCTGAATCCCAGGAAACAACTTAAAAAGATCCAGATCTGTTTGGACTATGAAGAAGAGACTGTGACTTTTTGTGATGCTGAAGGAGGAGGGCGCCTTGTTAAATTTAAACCAGCTCATTTTGCAGGAAAGAAAATCTTTCCTTTCTTCCGTGTTGGGGATATAAACACGTGCATCGGGCTG------------------------TGTTAA---------------------------------------------

>XM_053302967.1

---------------------------------------------------------------------------------------------------------------------------------------------------------------------------------------------------------------ATGAGTGCCTTGTTCTCATCCATTCTCACCATTTTCTTAGTGGTTTCGCCTATTAATAATGTGGACCCAGTCCAGTTGGCAGTGAATGGCCCACTCCACCCAGTCACCGCT---------------------------------------------------------------------------------------------------------------------------------------------------------------------------------------------------------------------------------------------------------------------------------------------------------------------------------------------------------------------------------------------------------------------------------------------------------------------------------------GCGCTGAATGAGGACATTGTGCTGCATTGTCATCTTTACCCCAGGACAAGTGCTCAGAATATGGAGATAAAATGGTTCCGCTCCCAGAACTCTTCATATGTATACTTTTATCACAGT------------------------------------------------------------------------------------------GGCAAAGATCATTTGGAAAGGCAGCAGCCTGAATACCAAGGAAGGACAGAATTTTTGAAAGATGGCCTTGGTGATGGAAAGATAGCTTTGAAAATTCTAAATGTCAGCCTTTCTGATGAGGGACAATACCACTGTTCTGTTAAAAATGGTAGTTTCCATCAGGAAGCCATGGCCACGTTGAACATGAAAGTGACTGCTTCAGGCACTGGTCCTCATATATAC---ATTGAAGGATATCAAGATAGAAGCATCCGAATTATTTGTCAATCCAATGGTTGG---------------------------------TACCCAAAGCCTGAGATTCTGTGGAGACATCCCAATGAGAAACATCTACTTTCACGGGCTACAAAAAATTCCCAGAAGCACAATGGGCTGTTTGAAGTACAAAATGATATAATCTTAACAGGAAGTTCAAATCAGAACTTGACCTGTGTGATCAGGAATATCCTCCTCAACCAGGAAAAGGAAGCAACCATTCATTTAGCAAATCACTTATTTCCAAAAACCTCTCACTGGATAGTGGTTCTGTGTGTCTCCCTGATGGCTGTGCTTGGCCTTATTGTCGTCATCCTTTACCTGTTTAACCAAGAGAAAAAGCTTGCTGCGGAACTTAAGTGG------------------------------------------------------------------------------------------------------------------------------------------------------------------------------------------------------------------------------------------------------------------------------AGGCAGTTGGTGATGCCGGTAGAAAAAGCAACTACTATAACTCTGGATCCAGACACAGCGAACCATGAGCTTATCCTGTCTGCAGACCGGAAGAACGTTATTCAA---GGGTTCATGTGGAACAGACTGCCTGACAATCCTAAAAGATTCAGTGCTGAACGTTATGTGTTGGGAAGTGACGGATTTTCCTCAGGGCGGCACTACTGGGAAGTGGAGGTGGGGGATCATGGGTATTGGGCTGTGGGGGTGGCTCGAGAGTCTGTGAGGAGGAAGGGGAAGCTC---------TGCCTTGACCCCAAAGAAGGAATCTGGGCTGTGGAGAAGTAT---------CATTTCCAATACCAAGCGCTGACC---ATCCCCAAGACTCCTCTCTCTCTGCAGAAGAGACCCAGGAAACTTGGGATTTATTTGGACTATGAGATGGAACAGGTGGCGTTCTATGATGTCAATCACAAGACACTCATATTTAGCTTCTTCTCAGCTGCTTTCAATGAGGAGAAGATCTTCCCTTTCCTGCATGTGGGG---GTAGGGTGCTGGCTTACCCTG------------------------TGCCCCTGA------------------------------------------

>XM_053302966.1

------------------------------------------------------------------------------------------------------------------------------------------------------------------------------ATGACCTCGGCTGCTCTGAATCTAGAAGAAGCTCTCCAGGATGAAGTCAGCTGTCCTGTTTGTCTAGATTATTTTCAGACTCCAGTGATGATCATAGATTGTGGGCACAACTTCTGCCGTGACTGCATCACCCGGTGCAGG---AAA------------------------------------------------------------------------------------------------------------------------------------------------------------------------------------------------------------------------------------------------------------------------------------------------------------------------------------------------------------------------------------------------------------------------------------------------------------------------------------GGACAAACTTCCTCCAAATCTTCCTGTCCACACTGCAGGAAGCCCTTTTCCTGGCAAAACCTCCGGCCAAACAGGCACCTGGCAAATATTGTGGAGCTGGCCAAACAGTTCAACAAG---------------------------------------------------------------------------------------TGGAAGGCAAATGGGACAGGGGAGCAGAAGCTGTGCAAGAAGCACCGGCAGCCCCTGGAACTCTTCTGTGAATTGGATCAAGCCCCCCTCTGCAGGCGGTGTGATATATTCAATGCGCATAGGGACCACCCTGTACTTTCCATAGAAAAAGCTGCTCTGGTTTACCAGGCTAAAAGTCATCGCTATTTGCAAATGTTGAAAGAAGAGAGAGAGGCAGTTATGTTA---CTAAAATCAGATAGGGAGAAGTCAACTCGGGAATTGCTGAGTGAAACAGAAGCT---------------------------------GAGAGACAGAAGCTTGTGTCAAAATTTCAGCAGCTGCACCATCTTCTTGAACAACAAGAACAAATCCTTTTGGACCAATTGAAAGATCTGACGATGGAGGTTGAAAAAAAGAAGGAAGAACACGCAACTAAATTTTCTGAGGAGATCTCTCATCTCGGTGCCCTTATCAATGAAACTGATCAGATGAGCCAGAAGTCAGCAAATGAACTCCTGTGGAACCCTGGAAAC---TTCTTGAACAGACCCAAGAAGAGGAAGTTTCAGCCTCCTGCTGTGTCTGATTCATCAGATCTCAGGAAAAAGCTCAAGAAGTTCTCCAGACAAAGCACATCATTGCAAAGTGTTCTGAAGAAAATCAAA---------------------------------------------------------------------------------------------------------------------------------------------------------------------------------------------------------------------------------------------------GATAACCTGTCTGAACTAAAGTGGGTAGAAGAAAATGTATTGTTGGCTCCAGAAACAGCCCATCCCCAATATATTGTGTCTGCAGATCGGAAATGTGTAAGATGC---AGAGATATCCTTCAGGAATTTCCCTCCAGCTCAAAGAGAATTGGAAATTTTCACTGTGTACTTGGAATGAAGGGATTCATCTCCGGAAAACATTATTGGAAGGTCAATGTAGGCCATGGGAAAAGTTGGGCTTTAGGAGTA---------TCTGTGGAGAGAAAAGGAGATGTC---------TGTATTATTCCCAAGGAGGGGATCTTTGCTCTGGGGTGTCAG---------AATGGTTCCTATGTAGCATTCAGT---TCCCAAGTCACTCCACTATACCTGAACAACAACCCAGAAGAGATCAAGGTTTCTTTGGATTATGAAGAAGCAACAGTGTCATTTTGTGATACATATGACTACCGTGTTCTTTATATGTTCCAATCTGTCAATTTTAAAGGTGAGAAAATCTTCCCTTTTTTCCAAGTTGCAGATATGGGCACAATTCTCAGTTTA------------------------CATTAA---------------------------------------------

>XM_053293405.1

---------------------------------------------------------------------------------------------------------------------------------------------------------------------------------------------------------------------------------------------------------------------------------------------------------------------------------------------------------------------------------------------------------------------------------------------------------------------------------------------------------------------------------------------------------------------------------------------------------------------------------------------------------------------------------------------------------------------------------------------------------------------------------------------------------------------------------------------------------------------------------------------------------------------------------------------------------------------------------------------------------------------------------------------------------------------------------------------ATGTTTATGCAGTTCCACAAGAGCATTCTTGCGCAACTGCCGGAAATTGTGAAAATGCAGTTGTGCCTGAACATCACCCCC------------------------------------------------------------------------------------------------------------------------------------------------------------------------------------------------------------------------------------------------------------------------------------------------------------------------------------------------------------------------------AAAATTGCATTTGTGCAATTTTTGGAAGTTGGACCA---------------------------------------------------------------------------------------------------------------------------------------------------------------------------------------------------------------------------------------------------------------------------------------------------------------------------------------------------------------------------------------AGAGTCCAAGCAGACTTAATTAATTGTGCAACCAGCATAACATTTGATCCAAACACGGCACATCCAAGTCTTGCTGTTTCTGAAGACAGGATGTATGTGGAATCA---GGTGGTGTC---CAAGATGTGCCTGACAATCCAGAAAGATTCGATAGCACGGCCTGTCTGCTGGGTTCTCCAGGATTTACATCTGGGAAACATTACTGGGAAGTGAAGTATGAAAACCAGAGGGAATGGGCAGTAGGGGTGGCCAAGGGATCTGTGGAAAGGAAAGGCTATATC---------ACACTTACTCCCGAGGAAGGAATTTGGCAAGAGGGTCTCTGGTGGCTTCGGCGAATGGACACAACTTCCCAAACA------------------------CTTCCCAACCAACCTGAAAAGATTGGGGTGTTTCTGGACTATGAACATGGTACTGTGTCTTTTTATATGGGTAGCAAAGTCATTCGA------------AAGAAAGCCTCCTTTGATGGGGAGAAAGTTTTCCCTTTCTTCTATGTGGGT---AGAGGTGTATGGCTCAAATTG------------------------ATCTCCTGA------------------------------------------

>XM_054978680.1

------------------------------------------------------------------------------------------------------------------------------------------------------------------------------------ATGGCCACTGCAAATCCAGCAAAGACCCTTCAGGATGACGCTACTTGTTCCATCTGTCTGGACTACTTCCAGGACCCTGTGATGATCATAGAATGTGGGCACAATTTCTGTCGAGCCTGCATCACCAAGTACTGG------------------------------------------------------------------------------------------------------------------------------------------------------------------------------------------------------------------------------------------------------------------------------------------------------------------------------------------------------------------------------------------------------------------------------------------------------------------------------------------AAAGCATCGGATTGCAGTGCCTGCTGCCCTGAATGCAGGCAACTGTTTTCCTGGAAGAATCTCAAACCCAACCGACAGCTGGGGAATATGGTAGAAACAGCTAAACAGCTCAATTTG---------------------------------------------------------------------------------------CAGCTGAAGAACAGCTCAGAAAAGGAGAGAATGTGTGAGGAACACCAAAAACCTCTGACTGTCTTCTGCAAAACAGATCAGACCCTCATTTGCACAGTCTGTGACCGATCTAAGGCTCACCGAAATCACGAAGTTGTTCATACACCAGAAGCTTCTGCGCATTACAAGGAGATGCTGCTGAGGCATCTAGAAAATTTGAAGAAAGAAAGAAACAAGATTCAGTCA---GCTAAATCAAATGGAGAAAAGCCATGCCAGGAGTTATTGAAAGAGACACAAGTC---------------------------------AAGAGGCAAGAGATTGTGTCTGAATTCCAACGACAGCATCAGTTTTTGGAAGAACAAGAACAGCTTCAGCTGTCCAAGCTGAGAGACCTGGAAAAGGAGATTGAGAGGAAAAGAGATAATTATGCAAGCAAGTTTGTGGATGAAATATCTTACATCACTGATCTGATTGGTGACCTGGAAAAAAAAGACAAGCAACCAACAAATGAATTTCTGCAGAATATTGGAAGC---GTATTAGACAGATGTAAGAAGGGAACATTTCAATATCCAGCTATACCTGATGCTTCAGATATGAAGAAAAGATTCTATCAATTCTGTGAAGGGAGCACATTTCTGCAAAGTTCTTTGAGGAAATTCAGA---------------------------------------------------------------------------------------------------------------------------------------------------------------------------------------------------------------------------------------------------GATAACCTGTTAAAACCAAAATGGATCAAAGAAAATGTGTTGTTTGATCCAGAAACAGCCCATCCCCGATATGTTGTGTCTGCAGATCGTAAAACTGTAAGATGG---GGGAGTATCCGTCAGGAATTCCCCTACAACCCAAAGAGATTTCACTATGTTCGCTGTGTTCTGGGCTGCAAGGGGTTCACTTCAGGGAAACATTACTGGACCGTGGATGTCGGAGATGGAGATTACTGGGCTGTGGGAGTGGCCAGAGAATCTGTGGAGCGGGAGGAGGAAATT---------GAGTTTGAACCTGATGAAGGAATCTGGGCTCTTGGGCTTTAT---------AATGATCAGTACAAAGCTCTGACT---TCACCTCCTACCCTACTGGATGTAGAGGATGAACCAACACAGATCCAGATTTCTTTGAACTATGAAGCGGGAACAGTTGCTTTTTATGATACTGAAGATAATACCCGTCTCTTCACTTTCCAGTCAGTTGATTTTGAAGGGGAGGAAATTTTCCCTTTTTTCCGGATTGTTGATTCATCAACTGTCCTGAAGTTG------------------------TGTTCTTAA------------------------------------------

>XM_054976459.1

---------------------------------------------------------------------------------------------------------------------------------------------------------------------------------ATGGCCACGTCCGGGGATCCTGCCGCGGCTCTCCAGGAGGAAGCGACCTGCTCCATCTGCCTTGATTATTTTCAAGACCCGCTGATGATCACCGATTGCGGGCACAATTTCTGCCGCACCTGCATCACCCAGTACAGG---GAA------------------------------------------------------------------------------------------------------------------------------------------------------------------------------------------------------------------------------------------------------------------------------------------------------------------------------------------------------------------------------------------------------------------------------------------------------------------------------------AAACTGGATCTCGGCGCCTCGTGTTGTCCCGAGTGCAGGAAACCCTTTTCCTGGGCCAATCTCCAGACAAACAGGCGCTTGGGGAATGTGGCGGAGCTGATCCAGCAGTTGCGTTTG------------------------------------------------------------------------------------CACCGGCTGAGCGAGCCCGATGGAGAGCAGGGGGTGTGCGGCAAGCACAAGGAGGTCCTCAAGCTCTTCTGCCCGAAGGACGGAGCTTTCCTCTGCTTGATCTGCAAAGAATCCCGAGCTCACAAAACGCACGCAGCGCTTCCCATAGACGAGGCGGTCCAGGATTACCAGGATCAAATCCGAAGTCACTTGACATTATTGAAGAAGGAGAGAGATGAAATTCTAAAG---AAAAAGGAAAGTAACGAGCAAGTAATTAAGGATACAATAAGGAATATCATACAT---------------------------------ATTAGGGAGCGGAAAGAGCGCACATTTCCAAAG------------------CAAGAAGCCCAGCCTCTGTTAGCCCTGCTAAAAAATGCCGAGGAAGAAGCTGTGCAGATGCTAGACGAAAATGCAAGCGATGCCTTGGAACACAGTTCCCATCTCACAGAGTTGATTGGAGAGATGGAGGGAAAGTGCCTGCAGCCAGCCCTTGAATTCTTGCAGGACATTGGGGCC---TTCCTGAGCAAATGTGAGAAGGAAATGGCTAGGGAACCC------------AAAGCTGAAATGAGTGTGACAGCTGCTTCCGTCCTTCAGAAAATTCTGTTGGTTCAGAGCAAACTGAATGTCTTTTCAGGAGACAACTCGCCACATGATCTCAGCAAGCCCAAGTCTGCAGGTACCACCTTGCTACCCCTCCAACAGCCAGCTCGCTCGCCCACTCACCTGCCCGCCAAT------------------------------------------CATCGGGGTGGATCCCTGGCTCGCCTGCCCATCCATCCTTGGCCACAGCAAGGAGGAACAACTACAGGAGAGAGAGGATTTCCATATTCAGATTTGATTTTACAACCTCTGATTCCTACACCATTCATTGAATCATTGACATTTGATGTGGAGACAGCTCATCCTCGTCTGGTGGTCTCCAGAAGTGGCAAAAGCGTGAGATGGGGAGGAGATATCCAACATGCCCTGCTTCCTGGGCCCTGGAGATTTGACCATAGTCGCTGCTTGCTGAGTAGCCAAGGATTCACCTCTGGGTATCACTGCTGGATAGTAGAGGTGGTCAAGGAGGGGCCTTGGGCTATTGGGGTTGCTCTAGAGTCTGTGCAGAGGAAGGGGTCAGTC---------AACTTGATCCCTAGGGAAGGAATTTGGGCCATGGCATTTAAC---------AACAGCAAATACTTGGTTCATACA---TCTCCCCCTACCCCTCTGATTCTGAGCTCTTCACCCAGAATAATACAAGTGTATCTGCATTATGAAAATGGGAAGGTTGTATTTATGGACTTCCATAGTAAGACTGTACTGTATGAATTCCTGTCTGCCTCTTTCTTGGGGCAGAAAGTCTATGCTTTCTTCCGTGTTGGCGGACCATCAGCTCATCTCAGATTG------------------------TGCTGTACGGGCTATCTTGCTGGACTTCGTTTTCTGTAG------------

>XM_054976458.1_cds1

------------------------------------------------------------------------------------------------------------------------------------------------------------------------------------ATGGCTACTCCAAATCCAGAAAAATTTCTCCAGGATGAAGCCACCTGTTCCATCTGTCTGGATTATTTTCAGGATCCAGTGATGGTCATAAACTGTGGGCACAATTTTTGCCGAAACTGCATCACACAATGCTGCGAA---------------------------------------------------------------------------------------------------------------------------------------------------------------------------------------------------------------------------------------------------------------------------------------------------------------------------------------------------------------------------------------------------------------------------------------------------------------------------------------GGGTCACCTTTCAAAGCCATTCCCTGCCCACAGTGCAGAAGACCCTTTCCCTGGAAAAACCTTCATCCCAACAGACATCTGTGGAACATTGTGGACCTCGCCAAACAGTTCAGCAAC---------------------------------------------------------------------------------------------AAGCGGGCAAATGAAACTGGAGACTTGTGTCAGAAACACCGGGAGCCCCTGAAACTCTTCTGTGAACATGACCAGACTCCCATTTGTGTGGTGTGTGACAGGTCCAAGGCGCATAAAAATCACAGTGTGGTTCCCATAGAGGAGGCTGCTCAGGTTTACCGGGGTAAACTGCAGCATTATCTTCAAGTGCTGAAAGAAGACAGAGAGGAGATTTTGTCA---CATAAATCAGAGTTAGAGAAGCCAACCCAGGATCTACTGAAAGACACAGCTGCA---------------------------------GAGAGGAAGAAACTTTCATTGGAATTACAACAGCTGCGTAAGCTTTTTGAGAAAGAAGAACAGCGTCTGTTGTCCCAACTGGATGAAGTGGAAAAAGAAGTTGAGAAGAGAGAGGCTGAAAATGTGACTGCGTTTTCTGAGGAGATTTCCCGTCTTGATGCCTTGATCCGTGAGCTTGAGCAGAAGTGTCAGGAGCCACCATGTGGGCTTCTCCAGAATATTGGAAGT---ACCATGAACAGATGTAAG---GACAAGTTCCAGCTTCCTGTTACACGTGATTCTTCAGAGCTGAAGAGAAAATTGCAAATCCTCTCTAAGGAAAGTGCGTTACTACAGGCAGATTTGAAGAAATTCAAA---------------------------------------------------------------------------------------------------------------------------------------------------------------------------------------------------------------------------------------------------GAGACACTGTCTGAACCAAAGTGGATAGAAGAGAATGTGATGCTGGACCAGGAGACGGCTCATCCTAGATTTCTTGTGTCTGATGATCGGAGAAGCGTGTCATGG---GGATCGTTCCGGCAGGAGGTGCCCTACAGCCCCAAGAGGTTTGAGCCGGCCCGCTGTGTGCTTGGCTCTCATGGATTCATGTCAGGGAAACATCACTGGACGGTGGATGTGGAGGATGGAACTTTCTGGGCTGTGGGTGTGGCCCAGGAGTCTGTGCAAAGAAAAGGAAAGTTT---------AATTTTGTACCTGAGGAGGGGGTTTGGGCTGTGGGCTTTTCA---------AACGGTCAGTACAAAGCTCTAACT---TCACCTCCTGACTTTCTAATGTTACTTACCCACCCTAATAAGATCCAGGTTTATTTAGACTATGAAAACGAGGCTGTCACTTTCTGGGATCCAGAAGAGCATGCTGAGTTGTATCGCTTCGAATCAGCAAACTTTAAGGGGAAAACACTTTTTCCTTTCTTCCGGATTGGGGACATGAAGACTAGCCTTTTGCTA------------------------TGTTAA---------------------------------------------

>XM_054976458.1_cds2

------------------------------------------------------------------------------------------------------------------------------------------------------------------------------------------------------------------ATGACTTCCTTCCTGTTCATTCTCATACTTTCCTTCATTGCCCACCCTGTGAACAACATGGGTCCAGTCCAATTCACAGTGACTGGACCCCTTTCCCCAGTTACTGCC---------------------------------------------------------------------------------------------------------------------------------------------------------------------------------------------------------------------------------------------------------------------------------------------------------------------------------------------------------------------------------------------------------------------------------------------------------------------------------------ACTTTGGGTGAGGATGTTGTACTACGCTATCATCTGTTCCCCAGGACAAGTGTTCAGAACATGGAGATAAAATGGTTCCGTTCGCAGAACTCTTCATATGTACATTATTATCAGAGT------------------------------------------------------------------------------------------GGCAGAGATTATTCTGAAAGACAGCAAGTAAAATATCAAGGAAGGACACAGTTTCTGAGAGATGGCATTGATGAGGGAAAGGTTGACTTGAGAATTTTTAATGTCAGCCTTTTTGATGAAGGAGAATACTATTGTTCTGTGAACAATGGGAGT------TTCTATCAGGAAACCTCCTGGGATGTGAAAGTAACCGCATTGGGTCCTGCTCCAGTCATTTCC---ATTGAAGGTTATAAAAAAGCAGGGATCCGAGCTGTATGCCAGTCCAGCGGTTGG---------------------------------TACCCCAGACCCAAGATGCTCTGGAGAGACCCCAGTGGGAAACATCTTTCTTCGTTT---CAGAGAAATTTCCAGATGGATAATGGGCTCTTTGATGTACACAATGAGATCATCGTCACACAAAGTTCAAATCACAATTTAACCTGTGTGGTCAGGAACATTCTTCTTAACAAAGAAAAGGAATCAACCATTCATATAGCAATTGACGTTTTCCCCCATACATCACCCTGGATAGTGGGTCTGTGTATCTCCTTGGTGGTTTCGCTTGGTTTGGCTCTCTCTATTCCTTACCTGTTTAAAATAAACAGAACACTTGCTGCAGAACTTGGGTGG------------------------------------------------------------------------------------------------------------------------------------------------------------------------------------------------------------------------------------------------------------------------------AGGAATATGGTGATGCCAATA---GAGAAAGCAAACATAACCTTGGATCCAGAAACCGCGAACCATGACCTCATCCTGTCTGCAGACCAGAAGAATGTAATCCAA---GGCTTCATGTGGCACAGACTGCCTGACAACCCTAAAAGATTCGACATGGAACGTTGTGTGTTGGGGAGTGAGGGGTTTTCCTCAGGGAGACATTACTGGGAAGTGGAGGTGGGGGAGGAGGGGTACTGGGCAGTGGGGGTGGCCAGAGACTCTGTGAGAAGAAAAGGGCGGCTT---------ACCCTTGACCCCAAAGAAGGGATCTGGGCTGTAGAAAAGTGC---------CGGGTCCAATACCAAGCTCTCACT---GTTCCAGAGACGCCTCTATCTCTGAGGAAGAGCCCCCGGAAGCTTGGGATTTATTTGGACTATGAGATGGAGCAGGTGGCATTTCATGATGTCAGTCACAAGACCCCCATCTTCACCTTCCGTCTAGCTCCTCTCAATGGGGAGAGAGTCTTCCCTTTCCTTCATGTCGGG---GTAGGATGCTGGCTTACACTTTTGTTGGAGGCTTATGCACTGCCCTGTGAATGCATGAAGCTGCCTTCTACCGAATCAGGCACTTGGTCCATCAAG

>XM_054976457.1

---------------------------------------------------------------------------------------------------------------------------------------------------------------------------------------------------------------------------------ATGGCTCGTTCTATGGACTTCCCACAAACA---------------AATTGTGGC------------------------------------------------------------------------------------------------------------------------------------------------------------------------------------------------------------------------------------------------------------------------------------------------------------------------------------------------------------------------------------------------------------------------------------------------------------------------------------------------------------------------------------------------------------------------------------------------------------CTGTGGAAC---------------------------------------------------------------------------------------------------------------------------------------------------------------TCCAAAGGACCCTTCTTCCTCCTGTGT------------CTATTGCTATGTCTTCTTGCTGAGTGTGAAGGCAGCGAAAAA------------GTCCCAGTGTCCACAGTTACATTGCCAAAGGAAAAC---------------------------------------------------------------------------------------------------------------------------------------------------------------------------------------------------------------------------------------------------------------------------------------------------------------------------------------------------------------------------------------------------------------------------------------------------------------------------------------------------------------------------------------------------------------------------------------------------------------------------------------------------------------------------------------------------------------------------------------------------------------------------------------------------TACAAAACCAAAGTGACTTTTGATCCTGCCACAGCATATCCATGGCTCGTTGTGTCTGCAGACAGGACCTATGTGGAATCA---GGGAATAATTCCCAAAATGTGCCGGACACTCCTGAACGATTCAATAGCACACCGTGTCTGCTGGGCCTCCCAGGCTTTACATCTGGGAAGCATTACTGGGAGGTGGAGTATGGAAACCAGAGGGAATGGGCAGTGGGGGTGGCCCTTGAGACTGTGGATAGGAAGAAACATCTC---------TTACTTCGTCCCGAAGACGGGGTTTGGCAAGAGGGTCTCTGGTGGCTTCGGGTGCTGGAAAACGATTCTCATACA------------------------CTTCCCAGTCGCCCTGGGATCATTGGGATCTTTCTGGATTATGACGCAGATTCAGTGGCATTTTACATAGACAGGAAAGTGATTAAG------------AAAAAAGCCTTTTTCAAAGGGAAGAAAGTGTTTCCTTTCTTTTATCTGGGA---GGAGGTGTTCACCTCAAGATA------------------------ACAAAATAATTTTTTAAAAAGTCAGAGTAA---------------------

>XM_053379005.1

---------------------------------------------------------------------------------------------------------------------------------------------------------------------------------ATGGCCGCCGCCGGGGACCCCTCCAAAACTCTCCAGGACGAAGCGACCTGCTCCATCTGCTTGGATTACTTCAGCGACCCTGTGATGATCATCGCTTGCGGGCACAGCTTTTGCCGAGCCTGCATCGCCCAGTGCCCGGAG---------------------------------------------------------------------------------------------------------------------------------------------------------------------------------------------------------------------------------------------------------------------------------------------------------------------------------------------------------------------------------------------------------------------------------------------------------------------------------------GGGCCGAGCCGCGACCTCTCCTCTTGCCCCCAGTGCAGGATCGCTTTCCCCCGGGGCCACCTGCGGCGCAACAGGCACTTGGCGAATATGGCCGAAGCGATCCAGCGCCTCCGCCTG---------------------------------------------------------------------CAGAGCCTCCGCCTGGCCGAAGGGGAGCCCAAGGAGAAAGTGCCGAAGTTGTGCGGGAAGCACTACGAAGCCCTCCGGCTCTTCTGCCTAGAGGACCGCGCCTTCATCTGCTTGGTCTGCAGGGAATCGCGGGCGCACAAAGCCCACGTCGCCCTCCCCATCGACGAGGCTGCCCAGGATGCTAAGGATCACCTCCAAAGCTGCCTAACAACCTTAAAGAAGGAGAGAGATGAAATTTTGAAG---GGGAAATGGTGCAGTGAGAGAGAAATTCAGGAAATGAAGAAACGCGTGGGAAGC---------------------------------CTCAGGAATCAAGCTGCGGGAAAGTGGGAA------------------------------------------------CTGCTAGATGTGTGTGAAGAGATCATGAAGTCAATCGACAAAATAGCCAGTGAATTCTCAGAGCGCAGTTCTCATCTTGCCAAGCTGATTAAAGAGGCAGAGGAGATGTGTCTGAAGTCTGCCTTTGAACTCCTGCAGGCTATGGATCCC---TTGCTGAGCAGGTGCAACACAGAAATATATGGGAAACCC------------AAAGCTGATCTGAATAAGAAAGTTGAGTGGCTTAATGAAAAACTACAGCTGGTGCAGACCAAGTTGTGTGAGCTTTCA---------------------------------------------------------------------------------------------------------------------------------------------------------------------------------------------------------------------------------------GACAGAATGCATATCAATACAGGGAAATCACAATGGATAAGAGAAAACGTGGTGTTTGATCCGGAAACAGCCCATCCCCGATATATTGTGTCTCCAGATGGGAAAGCGTTATGGTGG---GGAAATGTCCGTCAGGACTACCCCTACAGTTCAATTCGGTTTGAATACGCCCGCTGTGTACTTGGAATGTGGGGATTCAGCTCAGGGAAACATTATTGGATGGTGGATGTCCGAAATGTCAACCATTGGGCTGTGGGCGTAGCCAGAGAGTCTGTGGAGAGGGATAGGGAAATT---------AATTTTGAACCCTATGAAGGGATCTGGGCAGTGGGCTTTAGC------CGCCATGATCAGTTCAAAGCTCTGACT---TCACCTCCTATCTATTTGGACCCAGAAGAGGGCCCAGCACAAATCCAAGTTTCTTTGAACTATGACGCAGGGAAAGTGTCTTTTTATGATGCTGAAGATAACACCTGCCTCTTTATTTTCGACTCAATTGATTTTGATGGGGAGGAAGTTTTTCCCTTTTTCCGGATTGTCGATTCATCAGCTTGCCTCCAGCTG------------------------TGCTATTAA------------------------------------------

>XM_053378956.1

------------------------------------------------------------------------------------------------------------------------------------------------------------------------------------ATGGCTACCGCAGATCCAGCAAAACACCTTCAGGACGAAGTCACTTGTTCCATTTGTCTGGATTACTTCAAGGATCCAGTTATGATCACGGAGTGCCAGCACGACTTTTGCCGAGCCTGCATCACAAAGTATTGG------------------------------------------------------------------------------------------------------------------------------------------------------------------------------------------------------------------------------------------------------------------------------------------------------------------------------------------------------------------------------------------------------------------------------------------------------------------------------------------AAAAGATCAGGAGCTTCCATTTGCTGCCCGGATTGCAGAAGAGCAGCTTCCTGGCAAAGCCTGAAACCCAACCGGCGCCTGGCGAACATGGTGGAAGCGGCTAAGCAGCTCCGTCTG---------------------------------------------------------------------------------------CAGCTGGAGCAAAGTCCAGGAAGGGAGAAAATGTGCAGGGAGCACAAGAAGCCTCTGAGTCTGTTCTGCACAACAGAAAATACCCTTATTTGCATGTTCTGCGAGAGATCCAAGGCTCACAGAAACCACCGTGTTATTTCCCCAGAAAGAGCTGCTGCAGATTACAAGGGTATACTACTTGGACATCTAAAGGAACTGAAAAAAGAAAGAAACAAGATTCTGTCG---GTTAAATCAAATGGTGAGAAGCCGTGCCAGGACCTATTGAAAGAGGCAGAAGCC---------------------------------AAGAGGGTGGAGATTGTGTCTGAATTCCAACAACAGCGGCAGTTTTTGGAGGAACAAGAGCAACGCCAGCTGGCCAGGTTGGAAGACATAAAAAGGGAGATTGAGAAGAGAAGAAATGATTATGCAAACACATTTGAGGATGAAATATCCTACATCAACGATCTGATTGGTGATATAGAAAAGAAGGGCACTCAATCACCAGATGGATTCTTGGAGGGTATTGGAAGC---ATACTGGACAGATGCGAGACAGAGAAATTTCAGTTTCCAGCTACTCCTGCTTCTGCAGACATGAAGGAAAGGCTGAGGAAATTCTCTCAAGAAAGTGCATCTCTGATGAGTTCTCTGAGGAGAGTCAGA------------------------------------------------------------------------------------------------------------------------------------------------------------------------------------------------------------------------GACGGTTCTGAGTCCAACTGCTCTCAGTCCAACACAGTCAAATCAAAATGGATAAGAGAAAATGTGGTGTTGGATCCGGAAACAGCCCATCCCCGGTATATTGTGTCTTCAGATGGAAAATCAGTACGGTGG---GGAGATGTCCGCCAGGACTACCCCTACAACCCAAAGAGGTTTGAATACGCCCGCTGTGTGCTTGGAACGCAGGGATTCAGCTCTGGGAAACATTATTGGACAGTGGATGTTGAGGATATTGACCACTGGGCTGTGGGCGTAGCCAGTGAGTCTGTGGAAAGGGATAGAGAAATT---------GATTTTGAACCTGATGAAGGGATCTGGGCGATGGGCTTTAGT---------TATGATCAGTACAAAGCTCTGACT---TCACCTCCTACCGTTTTGGAGCCAGAAGAGGATCCAGTAGAGATCCAAGTTTCTTTGAACTATGAAGCAGGAACTGTGGCTTTTTATGATGCTGAAGATAAGACCCGCCTCTTTATTTTCCAGTCAATTGATTTTGAAGGGGAGGAAGTTTTCCCCTTTTTCCGGATTGGAAATTCATCAGCTCCCCTCCAGCTG------------------------TGCTCTTAA------------------------------------------

>XM_053378765.1

------------------------------------------------------------------------------------------------------------------------------------------------------------------------------------ATGGCTGCGCCGAACCCAGAGAAAAGCCTCCAGGATGAAGCCACCTGTTCCATCTGCCTGGATTATTACCAGAACCCGATGATGGTCATCGACTGTGGCCACAACTTCTGCCGGGATTGCATTGCCCAGTGCTGTGAA---------------------------------------------------------------------------------------------------------------------------------------------------------------------------------------------------------------------------------------------------------------------------------------------------------------------------------------------------------------------------------------------------------------------------------------------------------------------------------------GGCTCCATTGCCAATGTGTTCTCCTGTCCGCAATGCAGAAAACCCTTCCCTTGGCAGAACCTGCGGCCAAACAGGCACCTGTGGAATATCGTTGAGCTGGCCATGCAGTTCAGCAAG---------------------------------------------------------------------------------------CGGCGAGCCAAGGAGTTGGGAGAACAGACACTCTGCAAGAAACACCAGGAGCCGCTGAAGCTCTTCTGCGAATACGATCAAACGTCCATCTGCGTGGTTTGTGACAGGTCCAAGCTGCATAAATATCACAGCGTGATTCCTATAGAGGAGGCCGCTCAGGTTTACCAGGTAACGTGA------------------------------------------------------------------------------------------------------------------------------------------------------------------------------------------------------------------------------------------------------------------------------------------------------------------------------------------------------------------------------------------------------------------------------------------------------------------------------------------------------------------------------------------------------------------------------------------------------------------------------------------------------------------------------------------------------------------------------------------------------------------------------------------------------------------------------------------------------------------------------------------------------------------------------------------------------------------------------------------------------------------------------------------------------------------------------------------------------------------------------------------------------------------------------------------------------------------------------------------------------------------------------------------------------------------------------------------------------------------------------------------------------------------------------------

>XM_053378764.1

------------------------------------------------------------------------------------------------------------------------------------------------------------------------------------ATGGCTACCGCAGATCCAGCAAAACACCTTCAGGACGAAGTCACTTGTTCCATTTGTCTGGATTACTTCAAGGATCCAGTTATGATCACGGAGTGCCAGCACGACTTTTGCCGAGCCTGCATCACAAAGTATTGG------------------------------------------------------------------------------------------------------------------------------------------------------------------------------------------------------------------------------------------------------------------------------------------------------------------------------------------------------------------------------------------------------------------------------------------------------------------------------------------AAAAGATCAGGAGCTTCCATTTGCTGCCCGGATTGCAGAAGAGCAGCTTCCTGGCAAAGCCTGAAACCCAACCGGCGCCTGGCGAACATGGTGGAAGCGGCTAAGCAGCTCCGTCTG---------------------------------------------------------------------------------------CAGCTGGAGCAAAGTCCAGGAAGGGAGAAAATGTGCAGGGAGCACAAGAAGCCTCTGAGTCTGTTCTGCACAACAGAAAATACCCTTATTTGCATGTTCTGCGAGAGATCCAAGGCTCACAGAAACCACCGTGTTATTTCCCCAGAAAGAGCTGCTGCAGATTACAAGGGTATACTACTTGGACATCTAAAGGAACTGAAAAAAGAAAGAAACAAGATTCTGTCG---GTTAAATCAAATGGTGAGAAGCCGTGCCAGGACCTATTG---------------------------------------------------------------------------------------------------------------------------------------------------------------------------------------------------------------------------------------------------------------------------------------------------------------------------------------------------------------------------------------------------------------------------------------------------------------------------------------------------------------------------------------------------------------------------------------------------------------------------------------------------------------------------------------------------------------------------------------------------------------------------------------------GTAAGAACTGGCATAACTCAGGGGTCAGCAAACTTT---------------------------------TTCAGCAGGGGGCCGGTTCACTGTCCCTCAGACCTTGTGGGGGGCTCGGCT------------------------------------------------------------------------------------------------------------------------------------------------------------------------------------------------------------------------------------------------------------------------------------------------------------------------------------------------------------------GTTAACTGA------------------------------------------

>XM_053378966.1

------------------------------------------------------------------------------------------------------------------------------------------------------------------------------------ATGGCTGCGCCGAACCCAGAGAAAAGCCTCCAGGATGAAGCCACCTGTTCCATCTGCCTGGATTATTACCAGAACCCGATGATGGTCATCGACTGTGGCCACAACTTCTGCCGGGATTGCATTGCCCAGTGCTGTGAA---------------------------------------------------------------------------------------------------------------------------------------------------------------------------------------------------------------------------------------------------------------------------------------------------------------------------------------------------------------------------------------------------------------------------------------------------------------------------------------GGCTCCATTGCCAATGTGTTCTCCTGTCCGCAATGCAGAAAACCCTTCCCTTGGCAGAACCTGCGGCCAAACAGGCACCTGTGGAATATCGTTGAGCTGGCCATGCAGTTCAGCAAG---------------------------------------------------------------------------------------CGGCGAGCCAAGGAGTTGGGAGAACAGACACTCTGCAAGAAACACCAGGAGCCGCTGAAGCTCTTCTGCGAATACGATCAAACGTCCATCTGCGTGGTTTGTGACAGGTCCAAGCTGCATAAATATCACAGCGTGATTCCTATAGAGGAGGCCGCTCAGGTTTACCAGGAGAAAATACATCACTGCTTGCAAACTCTGAAAGAAGAGAGAGACAAGATTTTATCG---TTTAAATTAGATGCAGAGAAGCCAAGCCAGGATCTGCTGGCAGAAACTGAAGCC---------------------------------GAGAGGCAGAAGCTTGTGTTGGAATGTCAGCAGCTCCGCCAGTTGCTAGAGAAAGAAGAGCAGCTCCTGTTAACCCGGCTGCAGAAGCTGGAGATAGAGATTGAGAAGAAGAAGGAAGAGCATGCAGCTAAATTTTCCGAGGAGATTTCCCATATCAGCACCCTAATTGGCGAGCTGGAGCAGAAGGGTCAGGAGCCGCCAGGTGGGCTGATGCAGGATATTGGAAGT---ATCTTCAACAGGTGCAAGAGGGGCAAATTTCAGTTTCCTACTCCACCTGATTCTTCGGAGCTCAGTACAATGCTTCAGGAATTCTCCAAAGAGAGTGCTTCACTGCAGGGCAAACTCAGGAAATTCAGA---------------------------------------------------------------------------------------------------------------------------------------------------------------------------------------------------------------------------------------------------GAGACTTTGGATGAACAAAAGTGGATAAATGAGAACGTGACACTGGACCCCATGACGGCTCACCCTCGATTTGTTGTGTTTGATGGTCAGAAAGCTGTCGCATGG---GGATCTGTCCGCCAGGAGCTTCCCTACAGCACCGGACGATTTGACCCCACCCGCTGCGTGCTTGGCTGTCAAGGATTCACCTCAGGGAAGCATCATTGGACAGTGGATGTGAGCAAAGGAACTTTCTGGGCTGTGGGTGTAGCCAAAGAATCTGTGAAGAGGAAAGGACATCTG---------AATATTGTCCCTGAGGAGGGAATCATGGCTGTGGGTTTTAAC---------AACGGTCAGTACAAAACCTTAACG---TCGCCCCCGACCATTCTGAACCCAAGTAAACACCCCCAAAAGATTCAAATCCGTTTGGACTATGAAGGAGGAACTGTGGGGTTTTTTGATGCGGAAGAAAACAAGCGCCTTGTTCTCTTTAGTGTCAAATTT------GTGTCAAAAATATTTCCTTTCTTCCGAGTTGGGGACATGAACACTGTGCTCCGCCTG------------------------TGTTAA---------------------------------------------

>XM_053378997.1

------------------------------------------------------------------------------------------------------------------------------------------------------------------------------------------------------------------------------------------------------------------------------------------------------------------------------------------------------------------------------------------------------------------------------------------------------------------------------------------------------------------------------------------------------------------------------------------------------------------------------------------------------------------------------------------------------------------------------------------------------------------------------------------------------------------------------------------------------------------------------------------------------------------------------------------------------------------------------------------------------------------------------------------------------------------------------ATGGAGGCGCCCATCTTCTCTTGTGACCCAGAAATGCGTTCCTTTCTGTTCATTCTC------------------ACCATTTTCTTTGTTATTCCACCTATAAATAATGTGGACCCT------------------------------------GCAACAGGCCTTGCTCCCCTTATTTCC---ATCGAAGGCTATCAGGACAAGGGGATCCGAATAGTTTGCCGATCCAGTGGTTGG---------------------------------TACCCAAAGCCTGAGATCCTGTGGAGAGGTCCCAATGGGAGGCACTTCCCTTCACTGGGACAAAAAATTTCCATGGAGGGCAATGGGTTGTTTGAGGTGCAGAACGACATCATCCTAACAGGAAGTTCAAATCATAGCCTAACCTGTGTGGTCCGGAACAGCTTCCTCAACCAGGAAAAGGAATCAACTATTCATATAGCAGATCACTTCTTCCCCAAGATCTCTCAGTGGATAGCTGGTTTGTGTGTCTCCCTCTTGGCTCTTCTCGGTTTCATTCTCCTCCTCCTTTACCTGTTTAATATAAACAGAAAGCTTTCTGCAGAACTGAGTTGG------------------------------------------------------------------------------------------------------------------------------------------------------------------------------------------------------------------------------------------------------------------------------AGGAACTTGGTGATGCCAATC---AAAAAAGCAAATGTAACCCTGGATCCGGACACAGCGAACAACGAGCTCATCCTGTCTGCGGACCGGAAGACCGTCATACAA---GGCTTCACGTGGCACAGCCTGCCTGACAGCCCCCTGAGATTCAGCCTCGAGCGCTGCGTTTTGGGAAGCGAGGGCTTTTCCTCAGGAAGGCACTACTGGGAAGTCGAGGTGGGCGAGGAGGGGTACTGGGCCGTGGGGCTGGCAAGGGGTTCCGTGAAGAGGAAGGAGTGCCTC---------AGCCTTGACCCCAGCGAAGGGATCTGGGCCGTGGAGAAGTGC---------CGGGTCCAGTACCAAGCTCTCACC---GTCCCTGAGACGCCCCTGTCTTTGCGGAAGAGACCCAGGAAACTTGGGATTTACTTGGACTACGAGACGGGGCAAGTGGCGTTTCATGACGTCAACCACAAGACACCCATCTTCACTTTCCCCTCCGCTGCTTTCGACGGGGAGGTGGTCTTCCCTTTCCTGCACGTCGGG---GTGGGATGCTGGCTTCGACTG------------------------TGCCCGAGAATTTCCTTGTGTGAGGCTTAA---------------------

>XM_053379042.1

---------------------------------------------------------------------------------------------------------------------------------------------------------------------------------------------------------------------------------------------------------------------------------------------------------------------------------------------------------------------------------------------------------------------------------------------------------------------------------------------------------------------------------------------------------------------------------------------------------------------------------------------------------------------------------------------------------------------------------------------------------------------------------------------------------------------------------------------------ATGTCTTTTTCTCTGGTCTCTTGGCTGAGCTTCCGG---------------------------------------------------------------------------------------------------------------------------------------------------------------AACTTTCCAATGAGAAGTGGTGTTCAGCCGACGCTGAAAGGATTCTATTGTATCCTGTGGCTAATGCTGTGCTTTCTGGCTGAGCAGAAAGGTGGTGGAAAAGTCCTGGCTTCACTACCAAGGGGA------------------------------------------------------------------------------------------------------------------------------------------------------------------------------------------------------------------------------------------------------------------------------------------------------------------------------------------------------------------------------------------------------------------------------------------------------------------------------------------------------------------------------------------------------------------------------------------------------------------------------------------------------------------------------------------------------------------------------------------------------------------------------------------------------------------TGCTACCGAGTTGCAGCTAAAGTGACCTTTAACCCGGACACTGCCCACCCAGCACTTGTTGTGTCTACAGACCGGAAGACTGTGACATCG---GAAGGTGTGGAGCACCCCGTGCCTCCCAACCCCAAGAGGTTCACTAAAAGCCCTGCCGTGTTGGGCTCTCCAGGGTTTAAATCAGGGAAACATTGCTGGGAAGTGGTGTACGGAAACCAGAGGGAATGGGCGGTCGGGGTAGCGCGGGAGTCTGTGAAAAGGGATGTCTACCTC---------TCCCTTACCCCAGAGGAGGGGATTGTGCAGGAAGGTCTCTGGTGGCTTCGGCGTCGGCAAAGCGACCCCCAACCA------------------------CCTCCCCAAGGCTCTGGAACCATTGGGGTTCTTCTGGATTGCGATCAAGACACCGTGACTTTTTACATGGCTGGTAAAGTCATTAAG------------AAAGATGTTCCCCGCAACGGAGAGGTAGTTTATCCTTTCTTCTATGTGGGT---GGAGATGTTTCACTCCGCCTG------------------------AACGACTTAAAAGAGTGA---------------------------------

>XM_033141496.1

---------------------------------------------------------------------------------------------------------------------------------------------------------------------------------ATGGCCGCCGCCGGGGACCCCTCCATAACTCTCCAGGACGAAGCGACTTGCTCCATCTGCTTGGATTACTTCCGCGACCCGGTGATGATCATCGACTGCGGGCACAGCTTTTGCCGAGCCTGCATCACCCAGTGCTCGGAG---------------------------------------------------------------------------------------------------------------------------------------------------------------------------------------------------------------------------------------------------------------------------------------------------------------------------------------------------------------------------------------------------------------------------------------------------------------------------------------GGGCCGGGCCGCGACCTCTCCTCTTGCCCCCAGTGCAGGATAGCTTTCCCCCGGGGCGACCTCCGGCCCAACAGGCACTTGGCGAATATGGCCGAAGCGATCAAGCGCCTCCGCCTG---------------------------------------------------------------------CAGCGCCTGAGCCTGGCCGAAGGGGAGCCCAAGAAGGAAGCGCCAAAGTTGTGCGGGAAGCACTACGAAGCCCTCCGGCTCTTCTGCCAAGAGGACCGCGCCTTCATCTGCCTGGTCTGCAGGGAATCCCGGGCGCACAAAGCGCACGTCGCCCTCCCCATCGACGAGGCTGCCCAGGATGCTAAGGATCAGCTCCAAACTTGCCTAAGAACCTTGAAGAAGGAGAGAGATGAAATTTTGAAG---GAGGACTGGTGCAGTGAGAGAACAATTCAGAAGATGAAGGATGAGCTCCAAAGTTGCCTAACAATCTTGAAGAAGCATAGAGATGGAATTATGAAGCAGAAAACAGTTCAGAAATCGAAGAAAGGTGCGGAAAACACCAGGAATCAAATTGCGAGAAACCAGGAACTTCTAGATGGAGTTGAAGAGATTAAGAAGTCAATTGACAAAACAGCCAGGGAATTCTCAAAGCGCAGTTCTCATCTCGCCAAGCTGATTAAAGAGATAGAGGAGAAGTGTCTGCTGTCTGCCTTTGAACTCCTGCAGGATATGGATCCC---TTGCTGAGCCGGTGCAACATAGAAATATCTGGGAAACCC------------AAAGCTGATCTGAATGAGAAAGTTGAGTTGTTTAATGAAAAACTACAGCTGGTGAAGACCAAGTTGCGTGAGGTTTCA---------------------------------------------------------------------------------------------------------------------------------------------------------------------------------------------------------------------------------------GACCGTTATCATTCCAACACAGTGAAATCAAAATGGATAAAAGAAAATGTGGTGTTGGATCCGGATACAGCCCATCCCAGATGTATTGTGTCTTCTGATGGAAAATCAGTACGGTGG---GGAAAGGTCCGTCAGGACTACCCCTACAGTCCAAATAGGTTTGAATATGCTCGCTGTGTACTAGGAATGCAAGGATTCAGCTCCGGGAAGCATTATTGGACGGTGGATGTTGAGGATGTTGACCATTGGGCTGTGGGTGTAGCCAAAGAGTCTGTGGAGAGGGATAGAGAAATT---------GATTTTGAACCTGAAGAAGGGATCTGGGCAGTGGGGTTTAGC---------AATGGTCAGTTGAAAGCTCTGACT---TCACCTCCTACCCCTTTGGAACCAGAAGAGGATCCAAGAGAGATCCAAGTTTCTTTGAACTATGAAGCAGGGACTGTGGCTTTTTATGATGCTGAAGATAAGACCCGCCTCTTTATTTTCCGCTCAATTGATTTTGAAGGGGAGGAAGTTTTTCCCTTTTTCCGGATCGTCAATTCGTCAACTTGCCTCCAGCTG------------------------GGCTCTTAA------------------------------------------

>XM_033141517.1

---------------------------------------------------------------------------------------------------------------------------------------------------------------------------------ATGGCCGCCGCCGGGGACCCCTCCATAGCTCTCCAGGACGAAGCGACTTGCTCCATCTGCTTGGATTACTTCCGCGACCCGGTGATGATCATCGACTGCGGGCACAGCTTTTGCCGAGCCTGCATCACCCAGTGCTCGGAG---------------------------------------------------------------------------------------------------------------------------------------------------------------------------------------------------------------------------------------------------------------------------------------------------------------------------------------------------------------------------------------------------------------------------------------------------------------------------------------GGGCCTGGCCGCGACCTCTTCTCTTGCCCCCAGTGCAGGATAGCTTTCCCCCGGGGCAACCTCCGGCCCAACAGGCACTTGGCGAATATGGCCGAAGCGATCAAGCGCCTCCGCTTG---------------------------------------------------------------------CAGCGCCTGAGCCTGGCCGAAGGGGAGCCCAAGAAGGAAGCGCCAAAGTTGTGCGGGAAGCACTACGAAGCCCTCCGGCTCTTCTGCCAAGAGGACCGCGCCTTCATCTGCCTGGTCTGCAGGGAATCCCGGGCGCACAAAGCGCACGTCGCCCTCCCCATCGACGAGGCAGCCCAGGATGCTAAGGATCAGCTCCAAAGTTGCCTAATAACCTTGAAGAAGGAGAGAGATGAAATTATGAAG---CGGAAATGGTGCAGTGAGGGAGGAATTCAGGAAATTAAGAAAGATGTGGAAAAC---------------------------------ATCAGGAATCAAGTTGCGGGAAGGTGGGAA------------------------------------------------CTGCTAGATGCGTGTGAAGAGATCATGAAGTCAATCGACGAAATAGCCAGTGAATTCTCAAAGCGCAGTTCTCATCTCGCCAATCTGATTAAAGAGGTAGAGGAGAAGTGTCTCAAGTCTGCCTTTGAACTCCTGCAGGCTATGGATCCC---TTGCTGAGCAGGTGCAACATAGAAATATCT------------------------------------------------------------------------------------------------------------------------------------------------------------------------------------------------------------------------------------------------------------------------------------------------------------------------------------------------------------------------------------------------------------------------------------------------------------------GGGAAACCCAAAGCTGATCTGAATGAGAAAGTTGAGTTGTTTAAT------------------------------------------------------------------------------------------------------------------------------------------------------------------------------------------------------------------------------------------------------------------------------------------------------------------------GAAAAACTACAGCTGGTGACGACCAAGTTGCGTGAGGTTTCAGGTAACAGATTA---------------------------------------------------------------------TGTTTTAACTAG---------------------------------------

>XM_033141497.1

------------------------------------------------------------------------------------------------------------------------------------------------------------------------------------ATGGCTACAGCGGATCCAGCAAAACACCTTCAGGACGAAGTCACTTGTTCCATTTGCCTGGATTACTTCAAGGATCCAGTAATGATCACGGAGTGCCAGCACGACTTTTGCCGAGCCTGCGTCACTATGTTTTGG------------------------------------------------------------------------------------------------------------------------------------------------------------------------------------------------------------------------------------------------------------------------------------------------------------------------------------------------------------------------------------------------------------------------------------------------------------------------------------------AAAAGACCGGGAGCTTCCATTTGCTGCCCGGATTGCAGAAGAAAAGCTTCCTGGCAAAGCCTGAAACCCAACCGGCGCCTGGCGAACATGGTGGAAGCCGCTAAGCAGCTCAGTCTG---------------------------------------------------------------------------------------CAGCTGGAGCAAAGTCCAGGAAGGGAGAAAATGTGCAAGGAGCACAAGAAGCCTCTGAGTCTCTTCTGCAAAACCGAAAATACCCTTATTTGCATGTTCTGCGAGAGATCCAAGGCTCACAGAAACCACTCTGTTATTTCCGCAGAAACAGCTGCTGCGGATTACAAGGGCATACTACTTAGACATCTAAAGGAACTGAAAAAAGAAAGAAGCAAGATTCTGTCG---GTTAAATCAAATGGTGAGAAGCCATGCCAGGAGCTATTGAAAGAGGCAGAAGCC---------------------------------AAGAGGGAGGAGATTGTGTCTGAATTACAAAAACAGCGGCAGTTTTTGGAGGAACAAGAGCAAAGCCAGCTGGCCAGGTTGGAAGACATAAAAAGAGAGATTGAGAAGAGAAGAAATGATTATGCAAACAAATTTGAGGATGAAATATCCTACATCAGCGATCTGATTGGTGAGATAGAAAAGAAGGACAATCAATCAGCAGATGGATTCTTGGAGGGTATTGGAAGC---ATACTGGACAGATGCGAGACAGAGAAATTTCAGTTTCCAGCTACTCCTGCTTCTACAGACATGAAGGAAAGGCTCAGGAAATTCTCTCAAGAAAGTGCATCTTTGACGAGTTCTCTGAGGAGAGTCAGA------------------------------------------------------------------------------------------------------------------------------------------------------------------------------------------------------------------------GACTGTTCTGAGCCCAACTGTTCTCAGTCCAACACGGACAAATCAAAATGGATAAGAGAAAATGTGGTGTTGGATCCGGATACAGCCCATCCCAGATGTATTGTGTCTTCTGATGGAAAATCAGTACGGTGG---GGAAAGGTCCGTCAGGACTACCCCTACAGTCCAAATAGGTTTGAATATGCTCGCTGTGTGCTTGGAATGCAAGGATTCAGCTCCGGGAAACATTATTGGACGGTGGATGTTGAGGATGTTGACCATTGGGCTGTGGGCGTAGCCAGAGCATCTGTGGAGAGGGATAGAGAAATT---------GACTTTGAACCTGATGAAGGGATCTGGGCGATGGGCTTTAGT---------TATGATCAGTACAAAGCTCTGACT---TCACCTCCTACCATTTTGGAACCAGAAGAGGATCCAGTAGAGATCCAAATTTCTTTGAACTATGAAGCAGGGACTGTGGCCTTTTATGATGCTGAAGATAAGACCCGTCTCTTTATTTTCCGGTCAATTGATTTTGAAGAGGAGGAAGTTTTTCCTTTTTTCCGGATCGTCGATTCGTCAACTTGCCTCCAGTTG------------------------GGCTCTTAA------------------------------------------

>XM_033141533.1

------------------------------------------------------------------------------------------------------------------------------------------------------------------------------------ATGGCTGCGCCGAACCCAGAGAAAAGCCTCCAGGATGAAGCCACCTGTTCCATCTGCCTCGATTATTTCCAGAACCCGATGATGGTCATCGACTGCGGGCACAACTTCTGCCGGGACTGCATTGCCCAGTGCTGCGAA---------------------------------------------------------------------------------------------------------------------------------------------------------------------------------------------------------------------------------------------------------------------------------------------------------------------------------------------------------------------------------------------------------------------------------------------------------------------------------------GGGTCCGTTGCCAATGTGTTCTCCTGTCCGCAGTGCAGAAAACCCTTCCCTTGGCAGAACCTGCGGCCAAACAGGCACCTGTGGAATATCGTTGAGCTGGCCATGCAGTTCAGCATG---------------------------------------------------------------------------------------CGGCGAGCCAAGGAGTCGGGTGAACAGCAGCTCTGCAAGAAACACCAGGAGCCCCTGAAGCTCTTCTGCGAATACGATCAAACGTCCATCTGCGTGGTTTGTGACCGGTCCAAGCTGCATAAATATCACAGCATCATTCCTATAGAGGAGGCTGCTCAGGTTTACCAGGAGAAAATACATCACTGCTTGCAAACTCTGAAAGAAGAGAAAGACAAGATTTTATCA---TTTAAATTAGAAGCAGAGAAGCCAAGCCAGGATCTGCTGGCAGAAACTGATTCT---------------------------------GAGAGGCAGAAGCTTATGTTGGAATTTCAGCAGCTCCGCCAGTTTCTAGAGAAAGAAGAGCAAATCCTGTTAACCCGTCTGCAGAAGCTGGAGATAGAGATTGAGAAGAAGAAGGAAGAGCATGCAGCTAAATTTTCCGAGGAGATTTCCCGTATCAGCACCCTAATTGGCGAGCTGGAGCAGAAGGGTCAGGAGCCGCCAGGTGGGCTGATGCAGGATATTGGAAGT---GTCTTCAACAGGTGCAAGACGGGCAAATTTCAGTTTCCTACTCCACCTGATTCTTTGGAGCTCAGTACAATGCTTCAGGAATTCTCCAAAGAGAGTGCTTCACTGCAGGGCGAACTGAGGAAATTCAGA---------------------------------------------------------------------------------------------------------------------------------------------------------------------------------------------------------------------------------------------------GAGACTTTGGATGAACAAAAGTGGATAAATGAGAACGTGACACTGGACCCCATGACGGCTCACCCTCGATTTGTTGTGTTTGATGGTCAGAAAGCTGTCGCATGG---GGATCTGTCCGCCAGGAGCTTCCCTACAGCACCAGACGATTTGACCCCTCCCGCTGCGTGCTTGGCTCTCAAGGATTCACCTCGGGGAAGCATCATTGGACAGTGGATGTGAGCAAGGGAACTTTCTGGGCTTTGGGTGTAGCCAAAGAATCCGTGAAGAGGAAAGGGCAGCTG---------AATATCGTCCCTGAGGAGGGAATCATGGCTGTGGGTTTTAGC---------AACGGTCAGTACAAAACCTTAACG---TCGCCCCCGACTTTTCTGAACCCAAGCAAAGACCCCCGAAAGATTCAGATCCGTTTGGACTATGAAGGAGGAACATTGGGGTTTTTCGATGCAGAAGAAAATAAGCGCCTCGTTCTCTTTACTGTCAAATTT------GTGTCAAAATTATTTCCTTTCTTCCGAGTTGGGGACATGAACACTGTGCTTCGACTG------------------------TGTTAA---------------------------------------------

>XM_033141505.1

------------------------------------------------------------------------------------------------------------------------------------------------------------------------------------------------------------------------------------------------------------------------------------------------------------------------------------------------------------------------------------------------------------------------------------------------------------------------------------------------------------------------------------------------------------------------------------------------------------------------------------------------------------------------------------------------------------------------------------------------------------------------------------------------------------------------------------------------------------------------------------------------------------------------------------------------------------------------------------------------------------------------------------------------------------------------------ATGGAGGCACCCATCTTCTCTTGTGACTCAGAA------------------ATGTGTTCCTTTCTGTTCATTCTCACCATTTTCTTTGTTATTCCAACTGTAAATAATGTGGACCCT------------------------------------GCAACAGGCCCTGCTCCCCTTATTTCC---ATCCAAGGCTATCAGGACAGGGGGATCCGAATAGTTTGCCGATCCAGTGGCTGG---------------------------------TACCCAAAGCCTGAGATCCTATGGAGAGATCCCAATGGGAGGCATTTGCCTTCACTGGGACAAAAAATTTCCATGGAGGGCAATGGGTTGTTTGAGGTGCAGAACGACATCATCCTAACAGGAAGCTCAAATCATAGCCTAACCTGTGTGGTCCGGAACAGCTTCCTCAACCAGGAAAAGGAATCAACTATTCATATAGCAGATCACTTCTTCCCAAAGATCTCTCAGTGGATAGCTGGTTTGTGTGTCTCCCTCCTGGCTTTTCTCGGCATCCTTCTCCTCCTCCTTTACCTGTTTAATATAAACAGAAAGCTTTCTGCAGAACTGAGTTGG------------------------------------------------------------------------------------------------------------------------------------------------------------------------------------------------------------------------------------------------------------------------------AGGAACTTGGTGATGCCAATC---AAAAAAGCAAATGTAACCCTGGATCCGGACACAGCGAACTACGAGATCATCCTGTCTGCAGACCGGAAGAACGTCATACAA---GGCTTCACGTGGCACAGCCTGCCTGACAGCCCCCTGAGATTCAGCGTCGAGCGCTGCATTTTGGGGAGCGAGGGCTTTTCCTCAGGAAGGCACTACTGGGAAGTCGAGGTGGGCGAGGAGGGGTACTGGGCCGTGGGGATGGCAAGGGGTTCCGTGAGGAGGAAGGAGTGCCTC---------AGCCTTGACCCCAGCGAAGGGATCTGGGCCGTGGAGAAGTGC---------CGGGTCCAGTACCAAGCTCTCACC---GTCCCCGAGACCCCCCTGTCTTTGCGGAAGAGACCCAGGAAACTTGGGATATACTTGGACTACGAGACGGGGATAGTGGCGTTTCATGACGTCAACCACAAGACGCACATCTTCACTTTCTCTTTCGCTGCTTTCAACGGGGAGGTGGTCTTCCCTTTCCTGCAGGTCGGG---GTGGGATGCTGGCTTCGAATG------------------------TGCCCCTGA------------------------------------------

>XM_033141520.1

------------------------------------------------------------------------------------------------------------------------------------------------------------------------------------------------------------------------------------------------------------------------------------------------------------------------------------------------------------------------------------------------------------------------------------------------------------------------------------------------------------------------------------------------------------------------------------------------------------------------------------------------------------------------------------------------------------------------------------------------------------------------------------------------------------------------------------------------------------ATGTCTTTTTCTCTGGTCTCAATGAGAAGCAAT------------------------------------------------------------------------------------------------------------------------------------------------------------------------------GTTCAGCCGATGCTCAAAGGATTCTTTTGTATCCTGTGGCTAATGCTGTGCTTTCTGGCTGAGCAGAAAGGTGGTGGGAAAGTCCTGGCTTCCCTACCAAAGGAA------------------------------------------------------------------------------------------------------------------------------------------------------------------------------------------------------------------------------------------------------------------------------------------------------------------------------------------------------------------------------------------------------------------------------------------------------------------------------------------------------------------------------------------------------------------------------------------------------------------------------------------------------------------------------------------------------------------------------------------------------------------------------------------------------------------TGCTACCGAGTTGCAGCTAAAGTGACCTTTGACCCGGCCACTGCCCACCCAGCACTTGTTGTGTCTAAAGACCAGAAGACTGTGACATCG---GAAGGTGTGGACCACGTCGTACCTCCCAACCCCAAGAGGTTCACTAAAAGCCCTGCCGTGTTGGGCTCTCCAGGGTTTAAATCAGGGAAACATTGCTGGGAAGTGGTGTACAGAAACCAGAGGGAATGGGCGGTCGGGGTAGCGCTGGAGTCTGTGAAAAGGAATGTCTACCTC---------TCCCTTACCCCAGAGGAGGGGATTGTGCAGTATGGTGTCTGGTGGCTTCGGCGTCGGGAAAGCGACCCCCAACCA------------------------CCTCCCCAAGGCTCTGGAACCATTGGGGTTCTTCTGGACTGCGATCAAGACACCGTGACTTTTTACATTGATGGTAAAGTCATTAAG------------AAAAATGTTCCCCGCAACGGAGAGGTAGTATATCCTTTCTTCTATGTGGGT---GCAGAGGTTTCACTCCGCCTG------------------------AACGACTTGAGAGAGTGA---------------------------------

>XM_033141495.1

---------------------------------------------------------------------------------------------------------------------------------------ATGGCGGCGGCGGCGGCGGCGGCAGCGCCCGCAGGTAGTGGCTCCCGGGCCGCCCCCAACCCGGTGGCGACCCTCCACGAGGAGGCCGTGTGCGCCATCTGCCTAGACTACTTCATCGACCCGGTGTCCATC---GGCTGCGGCCACAACTTCTGCCGTGTGTGCATCACTCAGCTGTGGGGCTCG---------------------------------------------GGCTCGGAGGAAGGGGACGACGCGGGCGCCTCCGGCTCGGACGGAGGGTACGGGCCCGGCTCC---GACGCGGGCGCGGATGACGGAGGGGTCGACGACCTGGGGCCGGAGGAGGACGCCGACGACGACGAAGACGACCTCGAGGACAACGAGTACTTGGATGATGGCGGGGACATGGAAGATGAGGAGGAGGAGGTTGTGTCCAGGGAGGAGGAGTTCGACGACGAGGAGGACGACGACGGGTGGGATGATGAAGATGAGAGAGCGTACTGGGCCCAGGGCCCCGGGTCGGATGACATGTGGGACGATGACATGGGGAACGATTTGCTGTTCGACAATTAC---GATGACGAGGAGGTTATGGACGAGGAAGAGGAGCCGGAGGAT------GAGTATACCGTGCCAGCGACCCCGCCGACCCCCCAGATGCGCCCAGCTTTCAGCTGCCCCCAGTGCCGCAAGGTTTTCCCTCGCCGCAGCTTCCGCCCCAACCTTCAGCTGGCCAATATGGTGCAGATCATCCGCCAGATGCACCCGCAGCCGTTCAACTCCGTGGTGGCGTCCACGGAGGCCGTGGAACTGGCGGCAGCAGGGGATTCATCCGGAGCGCCATCTGTGTTAGCAGCGGGGCGAGCTCCAAGTGCTGAGCGAACCCTGTGCGAGAAGCACCAGGAGCCCCTCAAGCTTTTCTGTGAGGCAGATGAGGAGCCCATCTGTGTTGTGTGCAGAGAATCACGGGCCCATAAGCACCACAGCGTCATGCCGCTGGAAGAGGTTGTCCAAGAATACAAGGCCAAACTTCAAGGTCACTTGGACCCACTGAAGAAGAAACTGGAGACTGTTATGAAG---CAGAAGTTGGTCGAACAGGAGAAAATAGCAGACCTCAGGAACAAGATGAAGCTC---------------------------------GATATGAAAGAGTTTGAGGCAGACTTTGAGCGGCTGCACCAGTTCCTCATTGGGGAGCAGGCGCTGATGCTGCACCAGCTGGAGGACCGCTATGAGGTGCTGCTTGCCAGCCAGACTAACAATGTCTCCCACCTTGAGGAGCAAGGTGCCGCCCTCAGCCGCCTCATTGCAGAAGCGGAGGACAAGAGCAAGCAGGATGGCTTGCAGATGATCAAGGACTTCAAAGGC---ACTTTAGTCAGGTGTGAGAACATTAAGTTCCAGGATCCAGAGATGGTG---CCAGTGGATGCAGGAAAGAAATACCGAAACTACTTCCTTATAGACGTCTTAATGAAGAAG---------------------------------------------------------------------------------------------------------------------------------------------------------------------------------------------------------------------------------------------------------------------ATGGAGAAGATCTTCTGCCAAGTCCCTCAAGCTGATTTGATTCTGGACCCTGAGACGGCCCACCCGCGCCTCACCCTTTCCTCGGACTGCCTGGGCGTCCGGCTC---GGAGAGCGCTGGCGGGACCTGCCTGACAGCCCCAAGCGCTTCGACTCCGACTACTGCGTTCTGGCCTTGCAGGGTTTCACCTACGGCCGCCACTACTGGGAAGTGGAGGTGGGGGGCCGGCGGGGATGGGCTGTGGGGGTGGCCCGCGAATCGGCCCGGCGCAAAGAGAAGTCC---GGTGGCGGCTCCCACCAGAAGCGCGAGATCTGGTGCGTGGGGACCAAC---------GGCAAGAAGTACCAGGCTCTGACAACCACTGAGCAGACCTCCCTCTCCCCATCCGAGAAGCTCCGCCGCTTCTGCGTCTACCTGGACTATGAACGGGGCCAGCTGAGCTTCTACAATGCCGAGAGCATGGGCCATATCCATACCTTCAACGCCTCGTTC------CGCGAGCGCATCTTCCCCTTCTTCCGCATTCTTTCTAAGGGGACCCGCATCAAAATC------------------------TGCAATTGA------------------------------------------

>XM_042455062.1

------------------------------------------------------------------------------------------------------------------------------------------------------------------------------------ATGGCTGCTCTGAATCCAGAAAAATCCCTCCAGGATGAAGCCACTTGTCCCATCTGCCTGGATTATTTTCAGAGTCCGATGATGATCATCGACTGTGGGCACAACTTCTGCCAGGGTTGTATAACCCAGTACTGCGAA---------------------------------------------------------------------------------------------------------------------------------------------------------------------------------------------------------------------------------------------------------------------------------------------------------------------------------------------------------------------------------------------------------------------------------------------------------------------------------------GGGTCTCCTGGTGATGCATTATCCTGCCCACAATGCAGAAATCCCTTTCTCAGGAAAAACCTCCAACCCAACAGGCACTTGTTAAATATTGTGGCATTGGCCAAGGGTTTTGGTATG------------------------------------------------------------------------------------------CAGCGAGCCAAAGAAGGAGAGCAGAAGCTGTGTGAGAAACACCAGGAGCCTCTGAAACTCTTCTGTGAAGAGGATCAGGTTTCCATCTGTGTGGTTTGTGATAGGTCCAGGGTGCACAAAAACCACAGTGTTATTCCAATGGAGGAGGCTGCTCAGCTTTACAAGGAGAAAATAGAATGCTGTTTGCAGATTCTGAAAAAAGAGAGAGGCAAGATCTTATCA---TATAAATCAGATTCAGCCAAGTCAAATGAGGATCTTTTGAGCAAGATAAAAAAC---------------------------------GAAAGGAAGGATCTAGTTTCGGAATTTCAGCAACTGCGCCAGGTTCTCGAGAAGGAAGAGCAACTCCTGTTAGTCCAGCTGGAAGAACTGGAAGCAGAGGCTGAGAAAAAGAAGGCAGAG---GCAAATGAATTGTCTGCAGAGATTTCCCATCTTGACATCCTAATTGGTGAGCTTGAAAAGAAGTACCAGGAGTCAACAGTTGAAATGTTACAG---------------------------------------------------------------------------------------------------------------------------------------------------------------------------------------------------------------------------------------------------------------------------------------------------------------------------------------------------------------------------------------------------------------------------------------------------------------------------------------------------------------------------------------------------------TCTCCTATGATCAGTTTGACATCCTGCAATGGTGTGTTGTCA---AATTTTTCCTGGACT------------------------------------------------------------------------------------------------------------------------------------------------------------------------------------------------------------------------------------------------------------------------------------------------------------------------------------------------------------------------------------------TCTTCATAG------------------------------------------

>XM_042455066.1

------------------------------------------------------------------------------------------------------------------------------------------------------------------------------------------------------------------------------------------------------------------------------------------------------------------------------------------------------------------------------------------------------------------------------------------------------------------------------------------------------------------------------------------------------------------------------------------------------------------------------------------------------------------------------------------------------------------------------------------------------------------------------------------------------------------------------------------------------------------------------------------------------------------------------------------------------------------------------------------------------------------------------------------------------------------------------------------------------------------------------------------------------------------------------------------------------------------------------------------------------------------------------------------------------------------------------------------------------------------------------------------------------------------------------------------------------------------------------------------------------------------------------------------------------------------------------------------------------------------------------------------------------------------ATGAAAAAGCTCCAAAAATTAACCAAGGAGAGTGAGTCACTGCAGAAGGAACTGAAGAAATTCAGA---------------------------------------------------------------------------------------------------------------------------------------------------------------------------------------------------------------------------------------------------GAGACTCTGAAGGAACCAAAGTGGATAAAAGAGGATGTGACACTGGACCCAGTGACAGCTCACCCTCGATTTATCGTGTCTGAAGATTTGAAAAGTGTCAGGTGG---CGCTCTGTCCGTGAGGAATTGCCCTATGATGCCAGGAGATTTGACACTGTCCGCTGTGTCCTTGGCTCCCAAGGGTTCTCTTCAGGGAAGCATCACTGGATAGTGGACGTGGATCAAGGAAATTTTTGGGCTGTGGGGGCGGCCAGAGAATCTGTGAAGAGGAAAGGCCAGTTC---------AATATAGTTCCTGAGGAGGGAATCTGGGCTGTGGGTCTTCTC---------AGTGGTCAGTACAAAGTTCTAACT---TCACCTCCTACTATTATAAATCAGAGTAAGCATTTGAAGAAAATCCAGATTTGTTTGGACTGGGATGGAAAAACTCTGGCATTTTTTGATTTTGAAGAAAAGAAGCGCCTTGTTCTCTTTTCATCACTCACTTTTAAGTCAGAGAAAATATATCCTTTTTTCCGAGTTGGTGATATTAATACTGTCCTTTGTCTA------------------------TGTTAA---------------------------------------------

>XM_042455007.1

------------------------------------------------------------------------------------------------------------------------------------------------------------------------------------ATGGCTGCTGCTGGTGGACGCATGGATCCCCTGGAGGAAGCCATTTGCTCCATCTGCCTGGATCCCTTTCAGGACCCAGTGATGATCACCAGCTGTGGGCACAACTTCTGCAGAGCCTGCATCACTCAGTGCCAG------------------------------------------------------------------------------------------------------------------------------------------------------------------------------------------------------------------------------------------------------------------------------------------------------------------------------------------------------------------------------------------------------------------------------------------------------------------------------------------------------------TGCCTTTGCCCCCAGTGCAGGACCAGGTTCCCCCCCAGCAGCCTCAGGCCCAACAGGGAGCTGGGGAATGTGGTGCAGGCACTCCTCCAGCTCAAGCTG---------------------------------------------------------------------------------AGGAGCCTCCAGGATTCGAACCCAGACCCCAGGAGGATGTGCAAGAAGCACCAGGAAGTGGCCAAGCTCTTCTGCCTGGAGGACAAAGCCCTCCTCTGCCTGGTCTGCAGGGAGTCCAGGGCCCACAAGACACACTCTGCTCTCCCCATAGAGGAGGCAGCCCAGGAGTACCAATATGAAATTGAAAATTTCCTGAAAATACTGACAGATATAAAAGAGAAAACCCTGAAG---GCAAAATCATGTCATGAGAAAAAAATTCAGCAACTGAAGAGAGATGTGGGATAT------------------------------------------------------GCCTGGCAGCGCATTGCATTCAGATTTCCAAAGCAAGAAGCCAAGACTTTGGTGCCCCTGCTAGACACATGCGAAACCAAAATGGATCAGATTGTTGAC------------TTTGCGGAGCGTGGTCCCTGCCTGACCAAGTTGATTGGTGAGATAGAAGACAAGTGTGGGCAGACATCCTTCGCATTCTTGCAGGACATTGGTGAT---TTACTGAGCAGATGTGAGAAAGAAATAGAAAAACCC---------------AGACCCAATCTGAATGAGATAATTAGATTCGTCACCGAACAAATGAAGCTGGTTCACACCAAGTTGGCTCAATTTCCA---------------------------------------------------------------------------------------------------------------------------------------------------------------------------------------------------------------------------------------------------GAGAAACCATCAAAGCCAACATGGATAAAAGAAAACGTGATGCTGGATCCAGAAACAGCCCATCATCGGTACATCGTGTCTCAAGATCTGAAAAGTGTACATTGG---GCAAAAAGCCGTCAGGATCTGCCCTACAGTTCCAAAAGATTTGAGGGTACTCGCTGTGTGCTTGGAAGGAAGGGCTTCACCTCAGGAAAACATTATTGGATAGTGGATGTAGAGAGTGAAGGATACTGGGCAGTAGGTGTAGCCCAAGGATCTGTGGACAGGGAGGAGAAACTT---------GATATTGAACCAGATTCGGGAATCTGGGCTCTGGCACGTTAT---------AATGATCAGTACAAAGGTCTTACA---TCACCACCTACCCTTCTGGATCTAAACTATTATCCAGTAGAGATCCGGGTGTATCTGAACTATGATGATGGAACAGTGGATTTTTATGAT---GAAGATATGACCCGTCTCTTTATTTTCCAGTCAATTGATTTTCAAGGGGACGAAGTCTTCCCTTTTTTCCGGATTTTTAAGCCATCGTCTTCTCTACGCCTA------------------------TGTTAA---------------------------------------------

>XM_042455000.1

---------------------------------------------------------------------------------------------------------------------------------------------------------------------------------ATGAAACCTTCTGCTTTCCCTCAGGACCCAGGAATGAATTCCTACCTGTTCCTTTTCATCCTTTTCTTTGTTATTTCACCTGTTAATAATGCGGGCCCTGTCCAGTTGACAGTGAATGGACCACTCCACCCAGTCACTGCC---------------------------------------------------------------------------------------------------------------------------------------------------------------------------------------------------------------------------------------------------------------------------------------------------------------------------------------------------------------------------------------------------------------------------------------------------------------------------------------TCTTTGAGTGAGGATATTGTGCTACGCTGTCATCTGTCCCCCAGAACAAGTGCTGAAAACATGGAGATAACATGGTTTCGCTCCCAGAACTCTTCCTATGTGCATCTTTATCACAGT------------------------------------------------------------------------------------------GGCAAAGATCATTTGGAAAAGCAGCAGCCAGAATACCAAGGAAGAACAGAAATTTTGACAGATGGCATTGGCGACGGAAAAATTGGCCTGAGAATTCTTAATGTCAGCTTTTTTGACGAAGGACAGTACCACTGCTCTGTTAAGAAC------AGGAGCTTCCACCAGGAAGCCATACTGGATATAAAAGTGGCTGCATCAGGTTCCAGTCCTCTGATTTCA---ATTGAAAGCTATCAGGAGAAGGGGATATACTTGGTTTGTCGATCCTGGGGTTGG---------------------------------TACCCAAAGCCTGAGGTCTGGTGGAGAGATCCCAGTGGGAGGCATCTACCATCACTGTCTAAGGAAATATCCCAGAAGAACAATGGGCTGTTTGAAGTACAGAAGGGTATTCTCTTAACAGGGAGTTCAAATCAGCCCTTGACCTGTGTGGTCAGGAACATCCTCCTCAACCAGGAAAAGGGATCAACTATTCACATGGCAGATAACCTCTTCACAATGATTTCCTCTTGGACAGCTGGTCTGTGTGTATCGCTCCTAATATCTTTTGGCCTCGTTATTGCTATTCTTTATTTGATTAACAGAAACAGAAAGCTCATCACAGAACTGAGTTGG------------------------------------------------------------------------------------------------------------------------------------------------------------------------------------------------------------------------------------------------------------------------------AGGAAAATGGTGGTGCCAATA---GAGAAAGCCAACGTAACACTGGATCCAGACACAGCAAACCATGACTTGATTCTGTCTGCAGACCAGAAGAACGTCATACAA---GGCTTCATGTGGAACAGTCTCCCTGACAATCCTCAAAGATTTGACACGGAGCGCTGCGTGTTGGGAAAGGAGGGGTTTTGCTCGGGGAGGCATTACTGGGAAGTGGAAGTGGGCGAAGATGGGTACTGGGCCGTGGGTGTGGCTCGGTACTCTGTAAAGAGGAAAGGGAGGCTC---------AGCCTTGATCCTGATGAAGGAATCTGGGCTGTGGAAAAGTACTGGGTCCAAACTGTCCAGTACCAAGCTCTTACC---ATCCCACAGACTCCTCTGCCTCTACGAAAGAGACCCCGGAAACTTGGGATTTATTTGGACTATGAGTTGGAACAGGTGGCTTTTCATGATGTCAACCACAAGATGCACATCTTCACATTCTCCTCAGCAGATTTCAATGAGGAGAAGATTTTCCCTTTCCTGCATGTGGGG---GTAGGATGCTGGCTTAGACTG------------------------CGCCCTTGA------------------------------------------

>XM_042455067.1

---------------------------------------------------------------------------------------------------------------------------------------------------------------------------------------------------------------------------------------------------------------------------------------------------------------------------------------------------------------------------------------------------------------------------------------------------------------------------------------------------------------------------------------------------------------------------------------------------------------------------------------------------------------------------------------------------------------------------------------------------------------------------------------------------------------------------------------------------------------------------------------------------------------------------------------------------------------------------------------------------------------------------------------------------------------ATGAAAACAGGTGTCCTCCAGAGGTTCAAAGGACTGTGTTGCATTCTGTGGCTATCACTGTGCTTTCTTGGTGAATTGGAAGGTGGCAGAAAGGCTTCA------------------------------------------------------------------------------------------------------------------------------------------------------------------------------------------------------------------------------------------------------------------------------------------------------------------------------------------------------------------------------------------------------------------------------------------------------------------------------------------------------------------------------------------------------------------------------------------------------------------------------------------------------------------------------------------------------------------------------------------------------------------------------------------------------------------------------ATACGAAAAAAGAAATACAAAATCAATGTGAGCTTTGATCCTCAAACTGCACATCCAAATCTCCTTGTATCTCCAGACAGGAAGACTGTGACATGG---GTACCTAAACCCCAAGATGTGCCTGACAATCCAGAGAGATTCAACAGTACTCGGTGTTTGCTGGGTTTTCCAAGATTTAGTTCAGGGAAATATTACTGGGAAGTGGAATATGCAAACCAGAAGGAATGGGCTGTCGGGGTGGCCAGGGAGTCTGTGGAGAGGAAGAGGTATCTC---------AGGCTGACTGCTGAGGAAGGGATTTGGCAGAAGGGTCTCTGGTGGCTTTGGCGAAGGGGAACTCATTCCCGTCGA------------------------CGTCCCAACCACTCTAGAAAGATTGGGGTATTTCTGGATTATGAAAGGGGGAAGGTAACTTTTTACATGAAGAATAAGGCCATGGTAATC------------AATACCTCTTTTAATAGGGAAATAGTTCGGCCTTTCTTCTATGTGGGG---GCAACTGTTTCACTCACCCTG------------------------------TGA------------------------------------------

>XM_042447427.1

------------------------------------------------------------------------------------------------------------------------------------------------------------------------------------------------------------------------------------------------------------ATGCACCTGGCCATTAAGCAATGTTTCGGACTCTTTTGCTTTGTGTGCCTC------------------------------------------------------------------------------------------------------------------------------------------------------------------------------------------------------------------------------------------------------------------------------------------------------------------------------------------------------------------------------------------------------------------------------------------------------------------------------------------------------------------------------------------------------------------------------------------------------------------------------------------------------------------------------------------------------------------------------------------------------------------------------------------------------TTGCTGTGCATTCTGGCTGACCAGGAAGGAGGTGGAAAGGTCCTGGCTGTG---------------------------------------------------------------------------------------------------------------------------------------------------------------------------------------------------------------------------------------------------------------------------------------------------------------------------------------------------------------------------------------------------------------------------------------------------------------------------------------------------------------------------------------------------------------------------------------------------------------------------------------------------------------------------------------------------------------------------------------------------------------------------------------------------------------------------AAAAAGGAAAAATACAAAAGTGATGTGTCTTTTGATCCTGACACCGCCTATCCAAGGCTCATAGTGACTCCAGACAAGAAGACTGTGAAATCC---TCATCTACAGTCCAAGATGTGGCAGATAATCCAGGGAGATTCAGTAAAAATCCCTGTGTGCTCGGTAGTCCAGGATTTACCTCTGGGAAACATTACTGGGAGGTGGAGTACGGAAACTCGAGGGGGTGGGCTGTGGGGGTGGCCAGGAATTCTGTGAAAAGAAAAAATGGTCTT---------TCCCTTACCCAAGAAGAAGGGATTTGGCAACAGGGTCTCTGGTGGGCTCAGCAATTGGACCTTGGCTCACATGGA------------------------CTTCCCACCGGATGCGGAAAGATTGGCGTACTTCTGGATTATGAAGGGGACAACATGACCTTTTACAATGGCAACCAAGTTACTGAAATC------------AAGGCCTCTTTCAATGGTGAAAAAGTGTTCCCTTTTTTCTATGTGGGG---GCAGATGTTTCACTTCATTTA------------------------GTTCCTTGA------------------------------------------

>XM_008105630.3

ATGGGGAAACCGCAGGGAAAGAGGACAGAGTTCCCTCCCTCCCTGCCTGCCTGCTTCTCTGCTCTGGCGCCTCCCTTTCGCCTTGTCTTCTTCGGTTTCGCTTTCGTTTCTGAGCAGCCTCTCTTTCTGCATTCCCCTCCCACAGCTCTTCCCTCTCGTCTCCAGCAAAGAGGAGCCATGGCTGTTGGTGGGCACCCCTCGGCCAGGCTCCAGGACGATGCCACTTGCTCCATCTGCCTGGACTATTTCCAGGACCCGGTGATGATCATCGACTGCGGGCACAACTATTGCCGGGCGTGCATCTCGCAGTGCCAGGGGGAG------------------------------------------------------------------------------------------------------------------------------------------------------------------------------------------------------------------------------------------------------------------------------------------------------------------------------------------------------------------------------------------------------------------------------------------------------------------------------------AGG---------------TCCCTTTGCCCCCGCTGCAGGATCCCTTTCCCTTCGGACAATCTCCTGCCCAACAGGGATCTCCGGAATCTGGTGGAGGCTATCCGGCAGCTCAGCCTG---CAGCCGCTT---------------------------------------------------------------GAGAAGGCGGCCCGGCCAAAGCCGGCGGGGGGCGGCCCGAAGATGTGCGAGAAGCACCAGGAAGCCGCCAAGCTCTTCTGCCAGGAAGACAAAGCCTTCCTCTGCCTGATCTGCAGGGAGTCCCGAGCCCACAAAGCCCACACCGCGCTCCCCATCGAAGAGGCTGCCCAAGAGTACCGGGATCAAATCCAGACTTTTTTGCAAAAACTGGCAAAAGAAAGAGATGAACTACAGAGA---GCAGTTGTGTACAATAGGCACCATTATCAACAACTGAAGAGTATTGTGGAATCT---------------------------------GCCAAGCAGCGCATTGGACCCAATTTTCCAAAA---------------------AAAGAATCCCAGACTTTGATGCCCCTACTAAACACATGTGAAGCAAATCTGCAGATAGTTACTGAAGCATCCAATGATTTA------------TCCTGTCTTAATACGCTGATTACCGACATAGAAGAGAAGTCTAAGCAGTCATCCTTTGGACTCTTGCAGGGCATTGGTGAT---TTATTGAACAGATGTGAGCAGGAAACA---AGAAGACCC------------AGACGTGAACTGAAAACAATCCTTGAATCCATCGCTGAAAACATGGATCTGGTTAACACCAGGTTGGATCAGTATGCA---------------------------------------------------------------------------------------------------------------------------------------------------------------------------------------------------------------------------------------------------GAAAACACGCCAAAGAAAAAATGGATAAAAGAAAATGTGACGCTGGATGCAAAAACAGCCCATCCTCGGTACTTTGTGTCTGACGATCGGAAAAGCGTACACTGG---GCGAAGAGCCGCCAGGATCTGCCATACAGCTCCAAAAGATTTGAGTTTGCCCGCTGTGTGCTTGGGAAGAAGGGATTCACCTCTGGAAAACATTATTGGATCGTGGATGTAGAAGATGGTGACTATTGGGCGGTGGGCGTAGCCCAGGAATCTGTGGACAGGGATGGGGAGCTT---------GATTTTGAGCCTGATGAAGGAATCTGGGCTATGGCACGTTAC---------GACGATCAGTACAAAGCTCTTACT---TCACCCCCTACTCTTCTAGATCTAGACTATGATCCAAAAGAGATCCAGGTCTCTTTGAACTATGATGCTGGAATAGTGCATTTCTATGAT---GAAGATAAGAAACCCCTCTATTCCTTCCGATCAATCGATTTTGAAGGGGAGAAAGTCTTCCCTTTTTTCCGCATTGGTGATTCATCCACATCTTTGTACCTG------------------------CTCTAA---------------------------------------------

>XM_008105632.3

---------------------------------------------------------------------------------------------------------------------------------------------------------------------------------ATGAAATCTCCTGTCTTCCCTTATGCCTCGGAAATGAGCTGCTTTTGGTTCATTTTCATGCTTTTCTTTGTTATTTCACCTATTAATGATGTGAGCCCTGCCCAGTTGACTGTGAATGGGCCACTCCACCCAGTCACTGCC---------------------------------------------------------------------------------------------------------------------------------------------------------------------------------------------------------------------------------------------------------------------------------------------------------------------------------------------------------------------------------------------------------------------------------------------------------------------------------------TCTTTGAGTGAGGATATTGTGCTGCACTGTCATCTGTCCCCCAGAACGAGTGCAGAAAACATGGAAATAAAATGGTTTCGCTCCCAGAATTCTTCGTATGTGCACCTTTATCACAAT------------------------------------------------------------------------------------------GGCAAAGAGCATTTGGAAAAGCAGCAGCCAGAATACCAAGGAAGAACAGAATTGTTGACAGATGGCATTGAGGATGGAAAGATTGGCCTGAGGATTTTAAATGTCAGCTTCTATGATGCAGGACAGTATCACTGCTCTGTGAAGAAC------AAGAGCTTCCATCAAGAAGCCACACTGGATATAAAAGTGGCTGCATTAGGCTCTACTCCTCTCATTTCC---ATTGAACGTTATCAGGAGAGAGGGATCTACTTGGTTTGCCGATCCAGGGGTTGG---------------------------------TACCCACAGCCAGAGGTCTTGTGGAGAAATCCCAGTGGGAAGCATCTGTCCTTTTTGGCTAAGGAAACATCCCAGGAGAACAATGGGCTGTTTGAAGTACAGAAGGACATTCTCCTAACAGAAGGTTCAAACCAGCCCTTAACCTGTGTGGTCCGGAACAGGCTCCTCAACCAGGAGAAAGGATCAACCATTCACGTTGCAGATAACTTCTTCCCAAAGATTGATCCCTGGGCAGTTGGTCTATGTGTATCCATCCTAGTTCCTGTTGGCTTACTTATTGCTATTCTGTACCTGATTAACAGAAACAGAAAGCTCATCACAGAACTGAGTTGG------------------------------------------------------------------------------------------------------------------------------------------------------------------------------------------------------------------------------------------------------------------------------AGGAATATGGTGGTGCCCATA---GAGAAAGCCAATGTAACACTGGATCCAAGCACAGCAAACAAAGAATTGATCCTGTCTGCAGACCAGAAGAACGTCATACAA---GGCTTCATGTGGCACAACCTCCCTGACCATCCTTGGAGATTTGATGTGGAGCGCTGCATTTTGGGAAAAGAGGGCTTTTCCTCAGGGAGGCATTACTGGGAAGTGGAAGTCGGGGAAGAAGGGTACTGGGCTGTGGGGGTGGCTAGGGAATCTGTCAGGAGGAAAGGAAGGCTC---------AGCCTTGATCCAGATGAAGGCATCTGGGCTGTGGAGAAGGCTCGAGTCCAAACTGTCCAGTACCAGGCTCTTACC---ATCCCACAGACTCCTCTAACCCTGCGCAAGAGACCCAGGAAAGTTGGGATTTATTTGGACTATGAGCTGGGAAAGGTGGCCTTTCATGATGTCAATCACAAGATGCACATCTTCACATTCTTCGCTGCAGATTTCAATGGGGAAACAATTTTCCCTTTCCTACAGGTGGGA---ATAGGATGCTGGCTTACAGTC------------------------TGCCCCTGA------------------------------------------

>XM_062969547.1

------------------------------------------------------------------------------------------------------------------------------------------------------------------------------------------------------------------------------------------------------------------------------------------------------------------------------------------------------------------------------------------------------------------------------------------------------------------------------------------------------------------------------------------------------------------------------------------------------------------------------------------------------------------------------------------------------------------------------------------------------------------------------------------------------------------------------------------------------------------------------------------------------------------------------------------------------------------------------------------------------------------------------------------------------------ATGAAGGTGAGAGGTGTCTTCCAGACATTTGGAGGACTCTGCTGCATTCTATGGCTATCGTTGTGTTTTCTTGCCGAACTGGAAGGTGGTGGAAAGGTCCTGGCTTCAACACCA---------------------------------------------------------------------------------------------------------------------------------------------------------------------------------------------------------------------------------------------------------------------------------------------------------------------------------------------------------------------------------------------------------------------------------------------------------------------------------------------------------------------------------------------------------------------------------------------------------------------------------------------------------------------------------------------------------------------------------------------------------------------------------------------------------------------------------------GCCGATGTGCTTTTTGATCCAAACACTGCACACCCAAATCTCCATGTGTCTCCAAACAAGAAGAAGGTGACGTGG---CTACCTGAACCCCAAAATGTGCCTGACAACCCAGAGAGATTCAATGCCACCTATTGCTTGCTGGGTACTCCAAGATTCACCAAGGGGAAACATTACTGGGAAGTGGAGTATGGAGGCCAGAGGGAGTGGGTCGCAGGGGTGGCCCGGGAGTCCGTGCAAAGGAAAGGGTTTGTC---------AGAATGACGCCCAAGGAAGGGATTTGGCAGATGGGTCTTTGGTGGCTTCGTCGTGGGGAGATGGATTCCCACCCA------------------------CCTCCCAAGCACCCTGCAAAGATCCGGATATCTCTGGATTATGAGCAGGGGACAGTGACTTTTTACCTGGAGAATAAGGTCAATGTA------------CAAAAAGTCTCTTTTAAAGGGGAGAGTGTTCGCCCCTTCTTCTATGTAGGG---CCAACTGTTTCACTTGGACTC------------------------------TGA------------------------------------------

**All pry-spry domain containing proteins (including non-TRIM proteins)**

>anole_thaicobrin_XP_060615369.2

MKVRGVPQTFGGLCCVLWLSLCFLTEFEGGGKVLASTPANVLFDPITAHPNLRVSPNKKTVTWLPEPQNV

PDNPERFNASYCLLGTPRFTKGKHYWEVEYGGQREWAAGVARASVQRKGYVSMTPEEGIWQVGLWWLRRG

ETDSHQPPKHPAKIKISLDYEQGTVTFNLENKVTVQKASFKGEMIRPFFYVGATVSLKV

>anole_hTRIM_XP_067319465.1

MAVGGHPSARLQEDATCSICLDFFQDPVMIIDCGHNYCRACISQCQGQGSRCPRCRIPFPSENLRPNRDL

RNLVEAIQQLSLQPLEKTTRPKPAGGGPKVCEKHQEAAKLFCREDQAFLCLVCRESRAHRAHTALPIEEA

AQEYRDQIQTFLKKLAKERDEMQRVNLSNKHNYQQMKSNLESAKQRIGSNYPKKEAQTLMPLLNTCEANL

KILSEAPNDLSCLNKLISEIEKKSQQSSFGFLQGIGDLLNRCERETRRPRHDLRAIIGSLHEKLCVVNTK

LNQCAENTPKKPWIKENVTLDAKTAHPRFFVSDDRKSVHWAKTRQDLPYSSKRFEFARCVLGKKGFTSGK

HYWIVDVEDGDNWAVGVAQESVDRGGKLDFEPDEGIWAMARYDDQYKALTSPPTLLDLDYDPVEIQVSLN

YDAGIVHFYDEDKKPLFTFRSFDFEDEKVFPFFRIGNSSTSLYLL

>anole_BTN_XP_067319452.1

MKFPVFPYASEMSCFLFLFFVISPINNVSPVPFTVTGPLHPVIASLSEDIVLRCHLSPRMSAENMEIKWF

RSQNSSYVHLYHSGKEHLEKQQPEYQGRTELLTEGIGDGKISLRISNVSLFDEGQYHCSVKNRSFHQEVT

LDIKVAASGSTPLISIERYQERGIYLVCRSRGWYPEPDIFWRDPSGRHLSFLAEETSQKNDGLFEVQKDI

LLTEGSNQSLTCVVRNRLLNQEKESTIHFADNFFPKIDPWIVGLCVSILVPVGLLVAILYLINRNRKLIT

ELSWRNMVVPIEKANVTLDPDTANKELILSADQKNVIQGFMWHNLPDHPWRFNKERCILGKEGFSSGRHY

WEVEVGEEGYWAVGVARQSVRRRGKLSLDPNEGIWAVEKSRVQTVQYQALTVPLTPLLLRKRPRKVGIYL

DYEMGKVAFHDVNHKMHIFTFFEADFNRETVFPFLHVGIGCWLTVCP

>common_lizard_hTRIM_XP_060126836.1

MAAAGDPSITLQDEVTCSICLDYFRDPVMIIDCGHSFCRACISQCPEGPGRDLSSCPQCRIAFPRGNLRP

NRHLANMAEAIQRLRLQCLSLAEGEPKKEAPKLCGKHYEALRLFCQEDRAFICLVCRESRAHKAHATLPI

DEAAQDAKDQLQSCLTTLKKERDEILKRIEGCSEREMKNRVGSIRDQVAGNWELLLACAEIMESIDKIAS

EFSERSSHLAKLIKEVEEKCLKSAFELLQDMDPLLSRCNIEISGKPKADLNKKVELLNEKLQLVQTKLRQ

LSAGVPQQLNQHRYTSDFHYPVCDADYVDANTVKSKWKRENVVFDPETAHPRYIVSSDGKSVWWGKVRQD

YPYSSIRFEYARCLLGMWGFSSGKHYWTVDVEGVNHWAVGVARESVERDREINFEPDEGIWAVGFSRNDQ

FKALTSPPTYLDPEEGPTQIQVSLNYEAGTVAFYDAEDKTRLFIFESIDFEGEEVFPFFRIVDPSVCLQL

CY

>common_lizard_ligase_left1_XP_034960918.2

MATADPAKHLQDEVTCSICLDYFKDPVMITECQHDFCRSCITKYWKRSGASICCPDCRRKASWQSLKPNR

RLANMVEAAKQLRLQLEQSPGREKMCKEHRKPLSLFCKTENTLICMFCERSKAHRNHRVISAETAAADYK

GILLGHLKELKKERNKILSVKSNGEKPCQELLKEAEAKREEIVSEFQHQRQFLEEQEQRQLARLEDIKRE

VEKRRKDYANKFEDEISYISDLIGEIEKKGNQSADEFLEGIGSILDRCETEKFQFPATPASTDMKERLRQ

FSQESASLTSSLRRVRDCSEPNCSQSNKVNSKWKRENVVLDLETAHPRYVVSSDGKSVWWGDVRQDYPYN

PKRFQYARCVLGTQGFSSGKHYWTVDVEDVDNWAVGVARESVERDREIDFEPDEGIWALGFSYDQYKALT

SPPTILEPEEDPIEIQVSLNYEAGTVAFYDAEDKTRLFIFKSIDFEGEEVFPFFRIVDSSTPLQLCS

>common_lizard_ligase_left2_XP_034962875.1

MAAPNPEKSLQDEATCSICLDYFQNPMMVIDCGHNFCRDCIAQCCEGSIANVFSCPQCRKPFPWQNLRPN

RHLWNIVELAMQFSMRRAKESGEQKLCKKHQEPLKLFCEYDQTSICVVCDRSKLHKYHNVIPIEEAAQVY

QEKIHHCLQTLKEERDKILSFKLDAEKPSQDLLAETEAERQKLMLEFQQLHQFLEKEEQLLLTQLQKLEI

EIEKKKEEHAAKFSEEISHISTLIGELEQKGQEPPGGLMQDIGSIFNRCKRGKFQFPTPPDSSELSTMVQ

EFSKESASLQGKLRKFRETLDEQKWINENVKLDPMTAHPRFIVFDGQKAVAWGSVRQELPYSTGRFDPTR

CVLGCQGFTSGKHHWTVDVSKGTFWAVGVAKESVKRKGQLNMVPEEGIMAVGFNNGQYKTLTSPPTVLNP

SKHPQKIQIRLDYEGGTVGFFDAEEKKRLVLFSVKFVSKIFPFFRVGDMNTVLRLC

>common_lizard_BTN_XP_060126844.1

MSFSLVSRMSFRNFPTRSSVQPTLKGFYCILWLMLCFLAEQRGGGKVLASLPKGCYRATGPAPLISIKGY

QDQGIRIVCRSSGWYPKPEILWRDPNGRHLPSLGQKISMEGNGLFEAQNDIILTGSSNHSLTCVVRNSFL

NQEKESTIHIADDFFPKISQWIAGLCVSLMALLGFILLLLYLFNINRKLYAELSWRNLVMPIKKANVTLD

PDTAHDEIILSADQKNVIQGFTWHTLPDSPLRFSVERCILGSEGFSSGRHYWEVEVGEEGYWAVGLARGS

VRRKECLSLDPSEGIWAMEKCRIQYQALTVPETPLSLRKRPRKLGVYLDYEMGQVAFHDVNHKTPIFTFP

SAAFNGEEVFPFLQVGVGCWLRVCP

>common_lizard_tdhTRIMdup_XP_060126838.1

MAAAGDPSITLQDEATCSICLDYFRDPVMIIDCGHSFCRPCIAQCPEGPGRDLSSCPQCRIAFPRGNLQP

NRHLANMAEAIQRLRLQRLSLAEGEPKKEAPKLCGKHYEALRLFCQEDRAFICLVCRESRAHKAHAVLPI

DEAAQDAKDQLQSWLTTLKKERDDIMKQKSYTEKKNQESKKGVENIRNQIVGKRELLEALEEIDKIAREF

SNRSSHLAELIKEVEEKCLQSAFELLQDMDPLLSRCDIEISGEPKADLNKKVKLRDEKLQLVQTKLRQLS

DRSHSNTAKWIKENVVFDPETAHPRYIVSSDRKSVWWGKVRQDYPYSSNRFEYTRCVLGMRGFYSGKHYW

TVDVEDVDHWAVGVAKESVERDRKIDLEPDEGIWAVGFSNDQLKALTSPPTPLETEDVPTQIQVKLNYEA

GSVAFYDAEDKTRLFIFRSIDFEGEVVFPFFRIVNSSTCLQLYS

>common_lizard_ligase_right1_XP_034962878.2

MAAAGDPSITLQDEATCSICLDYFRDPVMIIDCGHSFCRPCIAQCPEGPGRDLSSCPQCRIAFPRGNLRP

NRHLANMAEAIQRLRQQRLSLAQGESKKKAPKLCGKHYEALRLFCQEDRAFICLVCRESRAHKAHATLPI

NEASQDAKDQLQSCLTTLKEERDEILKGIEVCSQREIQEMKNRLGSIRDQVAGNWELQVVGNCAELMQSI

DKIASEFSERSSHLAKLIKEVEEKCLKSAFELLQDMDPLLSRCNIEISGKPKADLNKKVELLNEKLQLVQ

RQLSEHPKPASLRADYVHANTVKSKWKRKNVVFDPETAHPRYIVSSDGKSVWWGNVRQDYPYSSIRFEYA

RCLLGMRGFSSGKHYWTVDVEGVNHWAVGVARESVERDREINFEPDEGIWAVGFRRNDQFKALTSPPTYL

DPEEGPTQIQVSLNYEAGTVAFYDAEDKTRLFIFESIDFEEEEVFPFFRIVDPSACLQLCY

>gecko_hTRIM_XP_060095237.1

MASANPVKDLQDEVTCSICLDFFQDPVMIVECGHNFCGACITKHWQGSDRSDLCPECRRLFSYQSLKPNR

QLGKTAEKVKQVSAQLKSNKADTLERMCKQHQKALTLFCKTEERLICQVCEKTKAHRSHAVVYAGEAAAD

YKEVLLRHLADLKQERNRILSIQSNGEKPSQNLLKEVQAMKLEIVSEFHEQQQFLEQQKQLQLSKLRDLE

KEIEKRRADYASKFVDEISYITDLIGDVEKKAKQPANELLQDIRYELDRYEKEEFKWPVVPHSSDVQKKL

NTFSQDCMYLQRILRKFRETPLKPTWIKENVLLDRETAHPRYVVSEDWKSVRWGDIRQEFPYNPKRFQYI

RGVLGCEGFTSGKHYWTVDVGNGGCWAVGVARESVEREVQIDFQPDEGIWAIAHSSDQYKALTSPPTLLD

VEDEPTQIKVCLNYEKGTVAFYDVYDDTRLFKFFSVNFEEEEVFPFFRIVDSSTVLELCS

>gecko_ligase_left1_XP_060094749.1

MAAPDPEKNLQNEATCSICLDYFQDPVMVIDCGHNFCRNCIAQCHNGSIFRLLICPQCRKPFPLKNLHPN

RHLWNIVELAKQFSKRRANETGDQKLCEKHLEPLKLFCEHDQTSICVVCDRSVAHKKHNVVPIEEAAQVY

KDKLHHYLQMLKEEKGKILSLKLDVEKPSQDLLKDTAAERQKLLSEFQQLHEFLEKHKQHLLSQVDEVEK

EVERKKMGDTAKFSDEISRLGALIQELEEKCQEHPGGFLQDIGSTLNRCKVNKFQPPAVLDCSDLKRRLH

VLSKESESLQEELNKFKAILSEPIWIEEYVVLDQGTAHPRFVVSDNRRMVTWGPFRQEVPYSPERFDPAR

CMLGSHGFSSGKHHWTVNVEDGTFWAVGVAQESVKRKGQFHFVPQEGIWAVGLHNGQYRALTSPPDLLLL

NSPLNKIQVCLDYEGGTVAFCDAEQQRQFYCFPANFEGEKLFPFFRVGDMKTTLFLC

>gecko_ligase_left2_XP_060094750.1

MATKSPTKSFQDEATCSICLEYFREPVTLNCGHNFCQACIAQYWETLEASTTCPQCRETTQQRSFRPNKQ

MANLVKIVKQMKAEQVMPAGEERACERHQEGLKLFCEEDQTPICLICHLSREHRDHTVFPIEEAAQDYKE

KIQIHLSILKEEKREQEKFHTAGKRKCQKNIKKIEAEKQKAVSVFQEMRCFLEEQEQLILAPLEQLKEKF

LKMRDELLTKLSTEVSCLATLITEMEKKCQQPASEFLEDIRSTLNKCQQKEVQKPADIPPELEKSLVNLS

EQNAVLTTAFKKCKDLLSSELKKYDPDVYEMGKIKHKVYGYLS

>gecko_btn_XP_060095246.1

MSSFVFILIISFIVFPVNNMGPVQLTAIEPLVSVTAILGEDVVLHYHLSPRTSAQDMEIKWFRSQNSSYV

HHYHNGKDYSERQQLKYQGRTEFLRDGIDAGKIDLRIFNVSLFDEGQYHCSLKNGTFYQEASWDVKVTAS

GSTPFISIKGYKKFGIRAVCQSSGWYPKPEMLWKDASGKRLSSFLERNVQKDNGLFEVQNEIIVTQSSDQ

NLTCVVRNILLHKQKESAIHIANDFFPKTPAWIVGLCISLMVSIGLALFILYLFKINRKLAAELGWRNMV

VPVEKANITLDPDTANHDLIISADQKNVIQGFMWHRLPDNPKRFDVERCVLGSEGFSSGRHYWEVEVGEE

GYWAVGVARDSVKRKGRLILDPKEGIWAVEKCPVQYQALTVPETPLPLQKKPRKLGIYLDCELKLVAFHD

VNHKAPIFTFPIAPLCGERIFPFLHVGVGCWLTICP

>gecko_thaicobrin_left_of_BTN_XP_060095244.1

MPQSTDSHLSFWNVSFTNCGLRTFKAVFIILWLSLCLPAECEGAKKVSASMIKQWRKNYKTKVTFDPNTA

YPWFIVSANKTYVESGDRRQNVPDNPERFNNTPALLGHPGFTSGKLYWEVEYGDQREWAVGVALERVDRK

VYLRLSPEEGIWQKGLWWLRALENRSHALPVRPGIIGIFLNYKAGTVAFYIQGKLILKRASFNREKVFPF

FYLGGGVHLKLI

>gecko_ligase_T41_left_of_BTN_1_XP_060095233.1

MAAAGPIGGGLLNPVATLHEEAVCAICLDYFTDPVSIGCGHNFCRVCITQLWGAAAEDSPDEEDEGEDDG

GASGSDGDYRPGSEEAVDDGGIDDLGPEEDGDDEDDLEEYLDGGDMVEEEDDDDASRDGYDEDDEIAWDE

EVNGDFWDQDPGSDETWEDAMGGDLFFDNYNEEEEVMEDEAVEDEADSPLTPPPRQSFTCPQCRKTFPHR

NFRPNLQLANMVQIIRQMHPQPFKSTTPPAEVVELAAAGDALGVSSALAAGRGVRGERVLCEKHQEPLKL

FCEEDEEPICVVCIKSRSHKHHNFVPLEEIVEEYKAKLQGHLDPLKKKLETVLKQKLSEQEKVSELRNKM

KLDMKEFESDFERLHQFLVGEQAQLLHQLEDRYNILLACHNSNVTHLEEQSATLSHLIAEAEDKSKQDGL

QLLKDFKGTLVRCENIKFQDPERVPVDTGKKYQNYFLLDTLMRKMEKVFCKVPQVDLTLDPETAHPSLTL

SSDCHSVQLGEHWRDVPNTSKRFNSDYCVLALKGFTWGRHCWEVEVSGGWGWAVGVTRKMARRKEKSSRG

PHLKREIWCVGTNGKKYQALTATEETSLSPSEKLRRFCVYLDYERGQLSFYNAESMTHIHTFNVSFHERI

FPFFCVLSKGPCIKLCN

>gecko_ligase_T41_left_of_BTN_2_XP_060095232.1

MAAAGPIGGGLLNPVATLHEEAVCAICLDYFTDPVSIGCGHNFCRVCITQLWGAAAEDSPDEEDEGEDDG

GASGSDGDYRPGSEEAVDDGGIDDLGPEEDGDDEDDLEEYLDGGDMVEEEDDDDASRDGYDEDDEIAWDE

EVNGDFWDQDPGSDETWEDAMGGDLFFDNYNEEEEVMEDEAVEDEADSPLTPPPRQSFTCPQCRKTFPHR

NFRPNLQLANMVQIIRQMHPQPFKSTTPPAEVVELAAAGDALGVSSALAAGRGVRGERVLCEKHQEPLKL

FCEADEEPICVVCRESHSHKHHNVVPLEEIVEEYKAKLQSHLDPLKKKLETVLKQKSSEQEKVAELRNKM

KLDMKEFESDFERLHQFLVGEQAQLLHQLEDRYDILLARQNSNVTHLEEQCAALSRLIAEAEDKSKQDGL

QLLKDFKGTLVRCENIKFQDPEMVPVDTGKKYRNYFLVDTLMRKMEKVFCKVPQVDLTLDPETAHPRLTL

SSDCRSVRLGEHWRDVPDTPKRFNSDYCVLALKGFTWGRHCWEVEVSGGWGWAVGAARETARRKEKSSRG

PHLKREIWCVGTNGKKYQALTTTEETSLSPSEKLRRFCVYLDYERGQLSFYNAESMTHIHTFNASFHERI

FPFFRILSKGTRIKLCN

>gecko_ligase_right_1_XP_060095238.1

MAAANPAKTLLEEATCSICLDLFQDPVMIIECGHNFCRKCITKYWKGSDCSACCPECRCLFSWTNLKPNR

QLGNMVETAKQLGLQLENSSEGERMCKEHQKPLTLFCKTDETLICQVCDRSIAHRSHTVVHTEEAAADYK

EMLLKHLRNLKEGRNKIQSAKSNGEKPSQELLKETQTKIQEIVSEFQQQRQFLEQQEQLHLSKLRDLEKK

IEKKRNDYASKCVHEISCITALIGDVEKKDKQPANEFLHDIGNILDRCKNGKFYYPAVPDTSDMKRRLNQ

LCQESTSLQSSFKNFRENMLKSKWIKENVLLDPETSHPRYIVSADRKTVKWGSARQQFPYNKKRFQCVRC

VLGSEGFTSGKHYWTVDVGDGDYWAVGVVRESVEREEEIPLEPDEGIWAIGLYDNQYKALTSPPTVLEVE

DEPTEIQVSLNYEAGTVNFYDAEDRTRLFTFQSVDFEGEEVFPFFRIVNSSTVLTMSY

>gecko_ligase_right_2_XP_060095236.1

MSTENIMAAANPAKTLLEEATCSICLDFFQEPVMIIECGHNFCRNCIIRYCKESDCRACCPECRCLFSWT

NLKSNRQLGNIVETAKELSLQLKNSWKSERICKEHQTPLTLFCKTDRTLICDVCDGFEAHQGHAVVHKEA

ATADYKETLLKHLKNLKEERNKFQTARSNGEKPSQELLQETQAKRQEIVSEFQKRRQVLEQQEQLQLSQL

RELEKEIERKRDDYADKCVYEISCINALIGDVERKNKQPANEFLQDIGSMLDRCKKGKFCYPVVPDTSAM

KGRLNQLPQESTSLQSSLKNFQANTSKSKWIKENVLLDPETAHPRYVVSEDQKTVRWGSFRQQFPYDPKR

FECIRCVLGSEGFTSGKHYWTVDVGDGDYWAVGVARENVEREEEIPLEPDEGIWAIGLYDNQYKALTSPP

TVLEVEDEPTEIQVSLNYEAGTVNFYDAEDKTRLFTFQSVDFEGEEVFPFFRIVNSSTVLTLSY

>gecko_ligase_right_3_ps_only_XP_060094748.1

MATGRDPGAAVRDETTCSICLDHFQDPLMIVDCGHDFCRACITRYSGGGSDLSVLSCPECRKPFSWGDLR

ANRRLGNVAELIQQFPLERLSEAEGAQKVCREHKEAFQFFCREDRAFICSICKDSEAHKTHVALPIDQAV

QNYQDQIQNYVTLLKKEKDEILKKINNEKAIKEVIRSIIYIREQKEHGFPKEEVHTLLALLRDVEEKTVQ

MADENASDTLEHSAHLTELIREMGAKCLQPAFEFLQMRERNG

>iguana_ligase_right1_pryspry_only

QANVTLDPVTAYPQLVLSEDGKKVKHGSSVFNPVTRSQGFDCVPCVLGCEGITSGRHFWDVEITAEGGDWAIGVVKGSIERKGMIAVGLAEGIWALQPETSHLLWRRGTTVRIRVYVDYECGRVAFFDVSSGDFIATLVPADFGGESIFPFFMLSKMGSQAKICP

>iguana_hTRIM

MAAGGNPSVKLQDEATCAICLDYFKDPVMIVVCGHNFCRACIGQYQEAGSGHDQSRCPQCRIHFLSGSLRPNRDLGNLVDAIRQLRLSDPVSGSKRSNPSWKVCEKHAEAAKLFCQEDKAFLCLVCRESRAHKAHSVFPVEEAVQDYQDQMQSFLTRLTKEREEILKANLYNQKEVPQLKVDLGSARQRIASRFPKEEARALLPLLDGCEAKMKTVIDMANDFSVRDSCLTKLISEIEEKRLQPSFEFLQDIGDLLSRCERELGKPRPDLKGIVGYITENMKWVHTKLDQYPEKTSNPKWIKENVMLDPETAHPRYIVSKDRKSVHWGDSRRDLPYNPKRFEYARCVLGKKGFTSGKHYWTVDVEDGEYWAVGVALESVDRDGELDFEPDEGIWALARYDDQYKALTSPPTLLDPDNDPVEIQISLNYDAGTVAFYDEDKTRLFIFRSIDFEGEKVFPFFRISDSSTSLRLC

>iguana_BTN_pryspry_only

LDPDTAHHDLILSADQKNVIQGFLWHRLPDHPHRFDTERCVLGKEGFSSGRHYWEVEVGEEGYWAVGVARDSVRRKGRFSLDPDEGIWAVEKYWVQTVQYQALTIPQTPLPLRKRPRRLGIYLDYEMEEVAFHDVIHKMHIFTFTSADFSGETIYPFLHVGVGCWLTLCP

>beardie_hTRIM_XP_020633187.1

MAVGGDPSLTLHDEATCSICLDYFQEPMMIVECGHNFCRACLADYEANEAGRDGLQCPQCRGAFAHHHLR

PNRHLGNMVDVIRRLNRQRLGKAGEEASRAASQVCEKHQEAVQLFCQEDRAFLCLVCRESRAHKAHAVLP

VDEAAQEYKDQLQGFLTTLTTERGEILKEEWCNQDEVQQMKRDIQIIQNGILTLFPEAKDQTQLALLDVC

VENMKAIDSIAHDFSERCFLLTKLISEIEDRLQQSTFKCLQNIGDLLSRCKREVSETAKADLKRRAESVI

EKLAHVSTKLNELAGSSRGQSPHPRGRSRETAASPSVNTSQHGQAFATVPSVFCPYSVVVAHPQQQQRGN

RKKWKKENVLLDPETAHPRYIISDDCKRVDWGDVRQDFFYNPKRFQYARCVLGTRGFKSGKHYWTIDVQD

GHHWAVGVARESVERDVVLSFEPDEGIWAMARYCDQYKAMTSPPTILELDDEPTEIQVSLNCDAGTVTFY

DAEDLTRLYKFECVDFEGEKVFPFFRISDSSTSLSITS

>beardie_ligase_left_1_XP_020633194.1

MAAPNPEKSLQDEATCSICLDYFQNPTMIIDCGHNFCEACIGQCCEGPVLNQFSCPQCRKPFFWQNVRPN

RHLWNIVELAKQFSLRRANEAGERKLCKKHQEPLKLFCEQDQTSICLVCDRSRVHKNHTVVPVEEAAQVY

QEKIHNCLKTLKEERGKILSFKSEWSKPSEDLLKETEVERQKLVSEFQQLRQVLEKEEQTLLTQLEKLQA

DAKKKMEEGEAKYSAEISHLTALIGEMEEKCQEPPGGLLQDIGCMFKRCQRGKFQSPAVPDSSELKKRLQ

IFSRDSTSLQGQLKKIRETLTEPKWIKEDMTLDVATAHPRFVVFEDRKTVAWGPVREELPYDARRFDPSR

CVLGSRGFTSGKHHWTVDVDQGTFWAVGVARESVKRKGQFTIIPAEGIWAVGLSSGHYKALTSPPVILSH

CKPMRKIRVCLDCEGKTVMFFDAEEKEAPVLFSFPISGSGKFFPFFRVGDMNTVLRLC

>beardie_trim41_left_of_RACK1_XP_020633185.1

MAAAAAASVPAGSVYGGPLNPVTTLHEEAVCAICLDYFIDPVSIGCGHNFCRVCITQLWGASEGQGGEEE

EEEEAEAAEEGEEVGTSGSDGGYGPASEDGGGEEGGIDDLPREEDAEDEEDDLEDDDYLDDGGMEEEEEE

EDVSRHEYDGDDDDVWDEGDDRDYWDEDPGSDEMWDDAMGSDLFFDNYGEEEEVMDEEEETEEDAEYNVP

TPPPPPPARQSFTCPQCRKNFPHRNFRPNLQLANMVQIIRQMHPQPFKPATASPETVELEAAGDSSGAPS

SSVAGRVPSAERTLCEKHQEPLKLFCETDEEPICVVCRESRNHKHHSVMPLEEVVQEYKAKLQGHLDPLK

RKLETVLKQKSREEEKIAELKNKMKLDMKEFESDFERLHQFLVGEQALLLHQLEDRYEVLLGTQNSNITH

LEEQGAALSRLIAEAEDKSKQDGLQLLKDYKGTLVRCENIKFQDPEIAPVDTGKKYRNYFLVDVLMRKME

KVFCKAPQADLTLDPETAHPRLILSSDYRSARLGDRWRELPDNPKRFDSDYCVLALQGFNQGRHYWEVEV

GGRRGWAVGAARESARRKEKANAGGPHQKREIWCVGTNGKKYQALTTTEQTSLPPCEKLRRFCVYLDYER

GQLAFYNAENMAHIHTFNASFRERIFPFFRILSKGTRIKICN

>beardie_thaicobrin_right_of_RACK1_XP_020633192.1

MQTGSVLQKFRGLCCITWLLLCLLAQYENGGKVLALSSSAKPYKTSVTFDPKTAHPNLVVSQDKKTVTWV

QEAQSVPDNPERFNSTPCLLGSPGFTSGKHYWEVEYGNQRELAAGVARKSVKRKDHLRLTPEEGIWQVGR

WWLRRGGPENQSDSGKIGVLLDYGRNQVTFYMDNKVTVVRASFNGEEVFPFCYVGSTVSLRLNP

>cliff_hTRIM_XP_053151972.1

MATASPAKCLQDEATCSICLDYFQDPVMISNCGHNFCRTCIIEYKKSDINICCPECRRSFSREHLKPNRP

LGRMAEAAKQLSLQINSPAKKRMCLKHKQPLIRFCKEDETLLCCLCEESKAHRNHTVVYAEAAAADYKMK

LQKRLENLKKEKNKILSVKSNGEKPSQELLKEMDDKKSSMVSEFQQIVQFLKEKEQFQLSRLGELEAETE

RTRKDYGNKFDDEISYITNLIGEIEKKNGQPPSEFLEDIGSILDRCEKEKFKFPPPPDTSDIKKRLEQFS

QQSTGEQSTLRKFRALQQQLQEEERRQQMGRNVAASAGARRGRNPAQEMQEELQQVVISRLRALQQQLQE

EERRQQMGRNVAASAGARRGRNPVQDNMLRPRWIKENVYLDPETAHPRYIVSQDGKSVEWGEIRQDFPYN

PKRFQYARCVLGCEGFTSGKHYWTVSVGDGDYWAVGVARESVERGEEIDFDPDEGIWALGRYDDQYKALT

SPPTLLELDYEPTEIQVSLNYEAGKVHFYDGEDKTRIYTFDADFEGEAIFPFFRIVDSSTCLQLCS

>cliff_left1_XP_053143803.1

MAALNPEKSLQDEATCSICLDYFQSPVMIIDCGHNFCRDCIGQCCEGSVTKVLSCPQCRKPFLWKNIRPN

RHLWNIVELAKQFSARRANEAGDQRLCEKHQEPLKLFCEQDRTPICVVCDRSKVHKNHNIIPIEEAAQAY

QDKIHNCLQTLKEEREKILSLKSEVEKPSQDLLKEAKAERQKLVAAFQELRELLEKEEHLLLAQLKNLET

EIEKKKEENASRYSEEISLLGSLIGELEQKRQEPPSGLLQDIGGILKRCKKDKFQPPVLDSSEFKRKLKE

FSQKSASLQGDLKKFRETLIEPKWIKENVLLDPLTAHPRFVVSDNQKSVAWGSFRMDLPYNAKRFDPARC

VLGSQGFSSGKHHWTVDVANGTFWAVGVARESVTRKGQLSFIPEEGIWAVGFYNGQYKALTSPPTFLNPR

KQLKKIQICLDYEEETVTFCDAEGGGRLVKFKPAHFAGKKIFPFFRVGDINTCIGLC

>cliff_BTN_XP_053158942.1

MSALFSSILTIFLVVSPINNVDPVQLAVNGPLHPVTAALNEDIVLHCHLYPRTSAQNMEIKWFRSQNSSY

VYFYHSGKDHLERQQPEYQGRTEFLKDGLGDGKIALKILNVSLSDEGQYHCSVKNGSFHQEAMATLNMKV

TASGTGPHIYIEGYQDRSIRIICQSNGWYPKPEILWRHPNEKHLLSRATKNSQKHNGLFEVQNDIILTGS

SNQNLTCVIRNILLNQEKEATIHLANHLFPKTSHWIVVLCVSLMAVLGLIVVILYLFNQEKKLAAELKWR

QLVMPVEKATTITLDPDTANHELILSADRKNVIQGFMWNRLPDNPKRFSAERYVLGSDGFSSGRHYWEVE

VGDHGYWAVGVARESVRRKGKLCLDPKEGIWAVEKYHFQYQALTIPKTPLSLQKRPRKLGIYLDYEMEQV

AFYDVNHKTLIFSFFSAAFNEEKIFPFLHVGVGCWLTLCP

>cliff_leftOfBTN_1_XP_053158941.1

MTSAALNLEEALQDEVSCPVCLDYFQTPVMIIDCGHNFCRDCITRCRKGQTSSKSSCPHCRKPFSWQNLR

PNRHLANIVELAKQFNKWKANGTGEQKLCKKHRQPLELFCELDQAPLCRRCDIFNAHRDHPVLSIEKAAL

VYQAKSHRYLQMLKEEREAVMLLKSDREKSTRELLSETEAERQKLVSKFQQLHHLLEQQEQILLDQLKDL

TMEVEKKKEEHATKFSEEISHLGALINETDQMSQKSANELLWNPGNFLNRPKKRKFQPPAVSDSSDLRKK

LKKFSRQSTSLQSVLKKIKDNLSELKWVEENVLLAPETAHPQYIVSADRKCVRCRDILQEFPSSSKRIGN

FHCVLGMKGFISGKHYWKVNVGHGKSWALGVSVERKGDVCIIPKEGIFALGCQNGSYVAFSSQVTPLYLN

NNPEEIKVSLDYEEATVSFCDTYDYRVLYMFQSVNFKGEKIFPFFQVADMGTILSLH

>cliff_left_of_BTN_2_psonly_XP_053149380.1

MFMQFHKSILAQLPEIVKMQLCLNITPKIAFVQFLEVGPRVQADLINCATSITFDPNTAHPSLAVSEDRM

YVESGGVQDVPDNPERFDSTACLLGSPGFTSGKHYWEVKYENQREWAVGVAKGSVERKGYITLTPEEGIW

QEGLWWLRRMDTTSQTLPNQPEKIGVFLDYEHGTVSFYMGSKVIRKKASFDGEKVFPFFYVGRGVWLKLI

S

>lepgecko_hTRIM_XP_054834655.1

MATANPAKTLQDDATCSICLDYFQDPVMIIECGHNFCRACITKYWKASDCSACCPECRQLFSWKNLKPNR

QLGNMVETAKQLNLQLKNSSEKERMCEEHQKPLTVFCKTDQTLICTVCDRSKAHRNHEVVHTPEASAHYK

EMLLRHLENLKKERNKIQSAKSNGEKPCQELLKETQVKRQEIVSEFQRQHQFLEEQEQLQLSKLRDLEKE

IERKRDNYASKFVDEISYITDLIGDLEKKDKQPTNEFLQNIGSVLDRCKKGTFQYPAIPDASDMKKRFYQ

FCEGSTFLQSSLRKFRDNLLKPKWIKENVLFDPETAHPRYVVSADRKTVRWGSIRQEFPYNPKRFHYVRC

VLGCKGFTSGKHYWTVDVGDGDYWAVGVARESVEREEEIEFEPDEGIWALGLYNDQYKALTSPPTLLDVE

DEPTQIQISLNYEAGTVAFYDTEDNTRLFTFQSVDFEGEEIFPFFRIVDSSTVLKLCS

>lepgecko_right1_XP_054832434.1

MATSGDPAAALQEEATCSICLDYFQDPLMITDCGHNFCRTCITQYREKLDLGASCCPECRKPFSWANLQT

NRRLGNVAELIQQLRLHRLSEPDGEQGVCGKHKEVLKLFCPKDGAFLCLICKESRAHKTHAALPIDEAVQ

DYQDQIRSHLTLLKKERDEILKKKESNEQVIKDTIRNIIHIRERKERTFPKQEAQPLLALLKNAEEEAVQ

MLDENASDALEHSSHLTELIGEMEGKCLQPALEFLQDIGAFLSKCEKEMAREPKAEMSVTAASVLQKILL

VQSKLNVFSGDNSPHDLSKPKSAGTTLLPLQQPARSPTHLPANHRGGSLARLPIHPWPQQGGTTTGERGF

PYSDLILQPLIPTPFIESLTFDVETAHPRLVVSRSGKSVRWGGDIQHALLPGPWRFDHSRCLLSSQGFTS

GYHCWIVEVVKEGPWAIGVALESVQRKGSVNLIPREGIWAMAFNNSKYLVHTSPPTPLILSSSPRIIQVY

LHYENGKVVFMDFHSKTVLYEFLSASFLGQKVYAFFRVGGPSAHLRLCCTGYLAGLRFL

>lepgecko_BTN_XP_054832433.1

MTSFLFILILSFIAHPVNNMGPVQFTVTGPLSPVTATLGEDVVLRYHLFPRTSVQNMEIKWFRSQNSSYV

HYYQSGRDYSERQQVKYQGRTQFLRDGIDEGKVDLRIFNVSLFDEGEYYCSVNNGSFYQETSWDVKVTAL

GPAPVISIEGYKKAGIRAVCQSSGWYPRPKMLWRDPSGKHLSSFQRNFQMDNGLFDVHNEIIVTQSSNHN

LTCVVRNILLNKEKESTIHIAIDVFPHTSPWIVGLCISLVVSLGLALSIPYLFKINRTLAAELGWRNMVM

PIEKANITLDPETANHDLILSADQKNVIQGFMWHRLPDNPKRFDMERCVLGSEGFSSGRHYWEVEVGEEG

YWAVGVARDSVRRKGRLTLDPKEGIWAVEKCRVQYQALTVPETPLSLRKSPRKLGIYLDYEMEQVAFHDV

SHKTPIFTFRLAPLNGERVFPFLHVGVGCWLTLLLEAYALPCECMKLPSTESGTWSIK

>lepgecko_FusedRightOfBTN_XP_054832433.1

MATPNPEKFLQD

EATCSICLDYFQDPVMVINCGHNFCRNCITQCCEGSPFKAIPCPQCRRPFPWKNLHPNRHLWNIVDLAKQ

FSNKRANETGDLCQKHREPLKLFCEHDQTPICVVCDRSKAHKNHSVVPIEEAAQVYRGKLQHYLQVLKED

REEILSHKSELEKPTQDLLKDTAAERKKLSLELQQLRKLFEKEEQRLLSQLDEVEKEVEKREAENVTAFS

EEISRLDALIRELEQKCQEPPCGLLQNIGSTMNRCKDKFQLPVTRDSSELKRKLQILSKESALLQADLKK

FKETLSEPKWIEENVMLDQETAHPRFLVSDDRRSVSWGSFRQEVPYSPKRFEPARCVLGSHGFMSGKHHW

TVDVEDGTFWAVGVAQESVQRKGKFNFVPEEGVWAVGFSNGQYKALTSPPDFLMLLTHPNKIQVYLDYEN

EAVTFWDPEEHAELYRFESANFKGKTLFPFFRIGDMKTSLLLC

>lepgecko_psOnly_leftofBTN_XP_054832432.1

MARSMDFPQTNCGLWNSKGPFFLLCLLLCLLAECEGSEKVPVSTVTLPKENYKTKVTFDPATAYPWLVVS

ADRTYVESGNNSQNVPDTPERFNSTPCLLGLPGFTSGKHYWEVEYGNQREWAVGVALETVDRKKHLLLRP

EDGVWQEGLWWLRVLENDSHTLPSRPGIIGIFLDYDADSVAFYIDRKVIKKKAFFKGKKVFPFFYLGGGV

HLKITKXFFKKSE

>aeolian_hTRIM_XP_053234980.1

MAAAGDPSKTLQDEATCSICLDYFSDPVMIIACGHSFCRACIAQCPEGPSRDLSSCPQCRIAFPRGHLRR

NRHLANMAEAIQRLRLQSLRLAEGEPKEKVPKLCGKHYEALRLFCLEDRAFICLVCRESRAHKAHVALPI

DEAAQDAKDHLQSCLTTLKKERDEILKGKWCSEREIQEMKKRVGSLRNQAAGKWELLDVCEEIMKSIDKI

ASEFSERSSHLAKLIKEAEEMCLKSAFELLQAMDPLLSRCNTEIYGKPKADLNKKVEWLNEKLQLVQTKL

CELSDRMHINTGKSQWIRENVVFDPETAHPRYIVSPDGKALWWGNVRQDYPYSSIRFEYARCVLGMWGFS

SGKHYWMVDVRNVNHWAVGVARESVERDREINFEPYEGIWAVGFSRHDQFKALTSPPIYLDPEEGPAQIQ

VSLNYDAGKVSFYDAEDNTCLFIFDSIDFDGEEVFPFFRIVDSSACLQLCY

>aeolian_left1_XP_053234931.1

MATADPAKHLQDEVTCSICLDYFKDPVMITECQHDFCRACITKYWKRSGASICCPDCRRAASWQSLKPNR

RLANMVEAAKQLRLQLEQSPGREKMCREHKKPLSLFCTTENTLICMFCERSKAHRNHRVISPERAAADYK

GILLGHLKELKKERNKILSVKSNGEKPCQDLLKEAEAKRVEIVSEFQQQRQFLEEQEQRQLARLEDIKRE

IEKRRNDYANTFEDEISYINDLIGDIEKKGTQSPDGFLEGIGSILDRCETEKFQFPATPASADMKERLRK

FSQESASLMSSLRRVRDGSESNCSQSNTVKSKWIRENVVLDPETAHPRYIVSSDGKSVRWGDVRQDYPYN

PKRFEYARCVLGTQGFSSGKHYWTVDVEDIDHWAVGVASESVERDREIDFEPDEGIWAMGFSYDQYKALT

SPPTVLEPEEDPVEIQVSLNYEAGTVAFYDAEDKTRLFIFQSIDFEGEEVFPFFRIGNSSAPLQLCS

>aeolian_left2_ringOnly_XP_053234740.1

MAAPNPEKSLQDEATCSICLDYYQNPMMVIDCGHNFCRDCIAQCCEGSIANVFSCPQCRKPFPWQNLRPN

RHLWNIVELAMQFSKRRAKELGEQTLCKKHQEPLKLFCEYDQTSICVVCDRSKLHKYHSVIPIEEAAQVY

QVT

>aeolian_left3_ringOnly_XP_053234739.1

MATADPAKHLQDEVTCSICLDYFKDPVMITECQHDFCRACITKYWKRSGASICCPDCRRAASWQSLKPNR

RLANMVEAAKQLRLQLEQSPGREKMCREHKKPLSLFCTTENTLICMFCERSKAHRNHRVISPERAAADYK

GILLGHLKELKKERNKILSVKSNGEKPCQDLLVRTGITQGSANFFSRGPVHCPSDLVGGSAVN

>aeolian_left4_XP_053234941.1

MAAPNPEKSLQDEATCSICLDYYQNPMMVIDCGHNFCRDCIAQCCEGSIANVFSCPQCRKPFPWQNLRPN

RHLWNIVELAMQFSKRRAKELGEQTLCKKHQEPLKLFCEYDQTSICVVCDRSKLHKYHSVIPIEEAAQVY

QEKIHHCLQTLKEERDKILSFKLDAEKPSQDLLAETEAERQKLVLECQQLRQLLEKEEQLLLTRLQKLEI

EIEKKKEEHAAKFSEEISHISTLIGELEQKGQEPPGGLMQDIGSIFNRCKRGKFQFPTPPDSSELSTMLQ

EFSKESASLQGKLRKFRETLDEQKWINENVTLDPMTAHPRFVVFDGQKAVAWGSVRQELPYSTGRFDPTR

CVLGCQGFTSGKHHWTVDVSKGTFWAVGVAKESVKRKGHLNIVPEEGIMAVGFNNGQYKTLTSPPTILNP

SKHPQKIQIRLDYEGGTVGFFDAEENKRLVLFSVKFVSKIFPFFRVGDMNTVLRLC

>aeolian_BTN_XP_053234972.1

MEAPIFSCDPEMRSFLFILTIFFVIPPINNVDPATGLAPLISIEGYQDKGIRIVCRSSGWYPKPEILWRG

PNGRHFPSLGQKISMEGNGLFEVQNDIILTGSSNHSLTCVVRNSFLNQEKESTIHIADHFFPKISQWIAG

LCVSLLALLGFILLLLYLFNINRKLSAELSWRNLVMPIKKANVTLDPDTANNELILSADRKTVIQGFTWH

SLPDSPLRFSLERCVLGSEGFSSGRHYWEVEVGEEGYWAVGLARGSVKRKECLSLDPSEGIWAVEKCRVQ

YQALTVPETPLSLRKRPRKLGIYLDYETGQVAFHDVNHKTPIFTFPSAAFDGEVVFPFLHVGVGCWLRLC

PRISLCEA

>aeolian_leftOfBTN_psOnly_XP_053235017.1

MSFSLVSWLSFRNFPMRSGVQPTLKGFYCILWLMLCFLAEQKGGGKVLASLPRGCYRVAAKVTFNPDTAH

PALVVSTDRKTVTSEGVEHPVPPNPKRFTKSPAVLGSPGFKSGKHCWEVVYGNQREWAVGVARESVKRDV

YLSLTPEEGIVQEGLWWLRRRQSDPQPPPQGSGTIGVLLDCDQDTVTFYMAGKVIKKDVPRNGEVVYPFF

YVGGDVSLRLNDLKE

>sand_hTRIM_XP_032997387.1

MAAAGDPSITLQDEATCSICLDYFRDPVMIIDCGHSFCRACITQCSEGPGRDLSSCPQCRIAFPRGDLRP

NRHLANMAEAIKRLRLQRLSLAEGEPKKEAPKLCGKHYEALRLFCQEDRAFICLVCRESRAHKAHVALPI

DEAAQDAKDQLQTCLRTLKKERDEILKEDWCSERTIQKMKDELQSCLTILKKHRDGIMKQKTVQKSKKGA

ENTRNQIARNQELLDGVEEIKKSIDKTAREFSKRSSHLAKLIKEIEEKCLLSAFELLQDMDPLLSRCNIE

ISGKPKADLNEKVELFNEKLQLVKTKLREVSDRYHSNTVKSKWIKENVVLDPDTAHPRCIVSSDGKSVRW

GKVRQDYPYSPNRFEYARCVLGMQGFSSGKHYWTVDVEDVDHWAVGVAKESVERDREIDFEPEEGIWAVG

FSNGQLKALTSPPTPLEPEEDPREIQVSLNYEAGTVAFYDAEDKTRLFIFRSIDFEGEEVFPFFRIVNSS

TCLQLGS

>sand_right1_XP_032997408.1

MAAAGDPSIALQDEATCSICLDYFRDPVMIIDCGHSFCRACITQCSEGPGRDLFSCPQCRIAFPRGNLRP

NRHLANMAEAIKRLRLQRLSLAEGEPKKEAPKLCGKHYEALRLFCQEDRAFICLVCRESRAHKAHVALPI

DEAAQDAKDQLQSCLITLKKERDEIMKRKWCSEGGIQEIKKDVENIRNQVAGRWELLDACEEIMKSIDEI

ASEFSKRSSHLANLIKEVEEKCLKSAFELLQAMDPLLSRCNIEISGKPKADLNEKVELFNEKLQLVTTKL

REVSGNRLCFN

>sand_left1_XP_032997388.1

MATADPAKHLQDEVTCSICLDYFKDPVMITECQHDFCRACVTMFWKRPGASICCPDCRRKASWQSLKPNR

RLANMVEAAKQLSLQLEQSPGREKMCKEHKKPLSLFCKTENTLICMFCERSKAHRNHSVISAETAAADYK

GILLRHLKELKKERSKILSVKSNGEKPCQELLKEAEAKREEIVSELQKQRQFLEEQEQSQLARLEDIKRE

IEKRRNDYANKFEDEISYISDLIGEIEKKDNQSADGFLEGIGSILDRCETEKFQFPATPASTDMKERLRK

FSQESASLTSSLRRVRDCSEPNCSQSNTDKSKWIRENVVLDPDTAHPRCIVSSDGKSVRWGKVRQDYPYS

PNRFEYARCVLGMQGFSSGKHYWTVDVEDVDHWAVGVARASVERDREIDFEPDEGIWAMGFSYDQYKALT

SPPTILEPEEDPVEIQISLNYEAGTVAFYDAEDKTRLFIFRSIDFEEEEVFPFFRIVDSSTCLQLGS

>sand_left2_XP_032997424.1

MAAPNPEKSLQDEATCSICLDYFQNPMMVIDCGHNFCRDCIAQCCEGSVANVFSCPQCRKPFPWQNLRPN

RHLWNIVELAMQFSMRRAKESGEQQLCKKHQEPLKLFCEYDQTSICVVCDRSKLHKYHSIIPIEEAAQVY

QEKIHHCLQTLKEEKDKILSFKLEAEKPSQDLLAETDSERQKLMLEFQQLRQFLEKEEQILLTRLQKLEI

EIEKKKEEHAAKFSEEISRISTLIGELEQKGQEPPGGLMQDIGSVFNRCKTGKFQFPTPPDSLELSTMLQ

EFSKESASLQGELRKFRETLDEQKWINENVTLDPMTAHPRFVVFDGQKAVAWGSVRQELPYSTRRFDPSR

CVLGSQGFTSGKHHWTVDVSKGTFWALGVAKESVKRKGQLNIVPEEGIMAVGFSNGQYKTLTSPPTFLNP

SKDPRKIQIRLDYEGGTLGFFDAEENKRLVLFTVKFVSKLFPFFRVGDMNTVLRLC

>sand_BTN_XP_032997396.1

MEAPIFSCDSEMCSFLFILTIFFVIPTVNNVDPATGPAPLISIQGYQDRGIRIVCRSSGWYPKPEILWRD

PNGRHLPSLGQKISMEGNGLFEVQNDIILTGSSNHSLTCVVRNSFLNQEKESTIHIADHFFPKISQWIAG

LCVSLLAFLGILLLLLYLFNINRKLSAELSWRNLVMPIKKANVTLDPDTANYEIILSADRKNVIQGFTWH

SLPDSPLRFSVERCILGSEGFSSGRHYWEVEVGEEGYWAVGMARGSVRRKECLSLDPSEGIWAVEKCRVQ

YQALTVPETPLSLRKRPRKLGIYLDYETGIVAFHDVNHKTHIFTFSFAAFNGEVVFPFLQVGVGCWLRMC

P

>sand_leftofBTN_psOnly_XP_032997411.1

MSFSLVSMRSNVQPMLKGFFCILWLMLCFLAEQKGGGKVLASLPKECYRVAAKVTFDPATAHPALVVSKD

QKTVTSEGVDHVVPPNPKRFTKSPAVLGSPGFKSGKHCWEVVYRNQREWAVGVALESVKRNVYLSLTPEE

GIVQYGVWWLRRRESDPQPPPQGSGTIGVLLDCDQDTVTFYIDGKVIKKNVPRNGEVVYPFFYVGAEVSL

RLNDLRE

>sand_trim41_XP_032997386.1

MAAAAAAAAPAGSGSRAAPNPVATLHEEAVCAICLDYFIDPVSIGCGHNFCRVCITQLWGSGSEEGDDAG

ASGSDGGYGPGSDAGADDGGVDDLGPEEDADDDEDDLEDNEYLDDGGDMEDEEEEVVSREEEFDDEEDDD

GWDDEDERAYWAQGPGSDDMWDDDMGNDLLFDNYDDEEVMDEEEEPEDEYTVPATPPTPQMRPAFSCPQC

RKVFPRRSFRPNLQLANMVQIIRQMHPQPFNSVVASTEAVELAAAGDSSGAPSVLAAGRAPSAERTLCEK

HQEPLKLFCEADEEPICVVCRESRAHKHHSVMPLEEVVQEYKAKLQGHLDPLKKKLETVMKQKLVEQEKI

ADLRNKMKLDMKEFEADFERLHQFLIGEQALMLHQLEDRYEVLLASQTNNVSHLEEQGAALSRLIAEAED

KSKQDGLQMIKDFKGTLVRCENIKFQDPEMVPVDAGKKYRNYFLIDVLMKKMEKIFCQVPQADLILDPET

AHPRLTLSSDCLGVRLGERWRDLPDSPKRFDSDYCVLALQGFTYGRHYWEVEVGGRRGWAVGVARESARR

KEKSGGGSHQKREIWCVGTNGKKYQALTTTEQTSLSPSEKLRRFCVYLDYERGQLSFYNAESMGHIHTFN

ASFRERIFPFFRILSKGTRIKICN

>fence_LeftOfHarbi1_XP_042310940.1

MDVWKAVKYIQEEATCPICLELFKEPVILDCGHNFCRACISRCQQEPKRKVSCPECRQSFVCANLRPNRQ

LGNILGLLKQFRVRDVESVEETVRLCESHQKPVWLFCSNDRVLLCLGCTESNPHRGHLLMPLEEAAQAYK

DEIHTLLEGKKKKREEVLKYFSSLGEQRYNLEKLMEHGTQKMVLEYMQLGEVMKSLQAEVRDLAKLMERD

FDSRLSKLLEEASHIETQVVEMERVCKQPAYEFLQGIKETLNKWEAEALGAPECVSPELKEKLWDLYVRS

LFIEDTLRKCTETLPLTPKVEKAKVTLDPATAHPLLALSEDGKKVKHGLSASRPTASSQGFDSVPCVLGC

EAITSGRHYWDVEITDEGGNWSIGVIKGSVERKGEIDINMDGGIWALEPESSHFRYRRGATITIRVFVDY

EGGRVAFFDVSSGDFVSTLVPADFGGESIFPFFMLSKVGTQAKVCP

>fence_LeftOfHarbi2_fragment_XP_042310996.1

MAALNPEKSLQDEATCPICLDYFQSPMMIIDCGHNFCQGCITQYCEGSPGDALSCPQCRNPFLRKNLQPN

RHLLNIVALAKGFGMQRAKEGEQKLCEKHQEPLKLFCEEDQVSICVVCDRSRVHKNHSVIPMEEAAQLYK

EKIECCLQILKKERGKILSYKSDSAKSNEDLLSKIKNERKDLVSEFQQLRQVLEKEEQLLLVQLEELEAE

AEKKKAEANELSAEISHLDILIGELEKKYQESTVEMLQSPMISLTSCNGVLSNFSWTSS

>fence_LeftOfHarbi3_fragment_XP_042311000.1

MKKLQKLTKESESLQKELKKFRETLKEPKWIKEDVTLDPVTAHPRFIVSEDLKSVRWRSVREELPYDARR

FDTVRCVLGSQGFSSGKHHWIVDVDQGNFWAVGAARESVKRKGQFNIVPEEGIWAVGLLSGQYKVLTSPP

TIINQSKHLKKIQICLDWDGKTLAFFDFEEKKRLVLFSSLTFKSEKIYPFFRVGDINTVLCLC

>fence_leftofHarbi4_XP_042310941.1

MAAAGGRMDPLEEAICSICLDPFQDPVMITSCGHNFCRACITQCQCLCPQCRTRFPPSSLRPNRELGNVV

QALLQLKLRSLQDSNPDPRRMCKKHQEVAKLFCLEDKALLCLVCRESRAHKTHSALPIEEAAQEYQYEIE

NFLKILTDIKEKTLKAKSCHEKKIQQLKRDVGYAWQRIAFRFPKQEAKTLVPLLDTCETKMDQIVDFAER

GPCLTKLIGEIEDKCGQTSFAFLQDIGDLLSRCEKEIEKPRPNLNEIIRFVTEQMKLVHTKLAQFPEKPS

KPTWIKENVMLDPETAHHRYIVSQDLKSVHWAKSRQDLPYSSKRFEGTRCVLGRKGFTSGKHYWIVDVES

EGYWAVGVAQGSVDREEKLDIEPDSGIWALARYNDQYKGLTSPPTLLDLNYYPVEIRVYLNYDDGTVDFY

DEDMTRLFIFQSIDFQGDEVFPFFRIFKPSSSLRLC

>fence_BTN_XP_042310934.1

MKPSAFPQDPGMNSYLFLFILFFVISPVNNAGPVQLTVNGPLHPVTASLSEDIVLRCHLSPRTSAENMEI

TWFRSQNSSYVHLYHSGKDHLEKQQPEYQGRTEILTDGIGDGKIGLRILNVSFFDEGQYHCSVKNRSFHQ

EAILDIKVAASGSSPLISIESYQEKGIYLVCRSWGWYPKPEVWWRDPSGRHLPSLSKEISQKNNGLFEVQ

KGILLTGSSNQPLTCVVRNILLNQEKGSTIHMADNLFTMISSWTAGLCVSLLISFGLVIAILYLINRNRK

LITELSWRKMVVPIEKANVTLDPDTANHDLILSADQKNVIQGFMWNSLPDNPQRFDTERCVLGKEGFCSG

RHYWEVEVGEDGYWAVGVARYSVKRKGRLSLDPDEGIWAVEKYWVQTVQYQALTIPQTPLPLRKRPRKLG

IYLDYELEQVAFHDVNHKMHIFTFSSADFNEEKIFPFLHVGVGCWLRLRP

>fence_leftofBTN_1_psOnly_XP_042311001.1

MKTGVLQRFKGLCCILWLSLCFLGELEGGRKASIRKKKYKINVSFDPQTAHPNLLVSPDRKTVTWVPKPQ

DVPDNPERFNSTRCLLGFPRFSSGKYYWEVEYANQKEWAVGVARESVERKRYLRLTAEEGIWQKGLWWLW

RRGTHSRRRPNHSRKIGVFLDYERGKVTFYMKNKAMVINTSFNREIVRPFFYVGATVSLTL

>fence_leftofBTN_2_psOnly_XP_042303361.1

MHLAIKQCFGLFCFVCLLLCILADQEGGGKVLAVKKEKYKSDVSFDPDTAYPRLIVTPDKKTVKSSSTVQ

DVADNPGRFSKNPCVLGSPGFTSGKHYWEVEYGNSRGWAVGVARNSVKRKNGLSLTQEEGIWQQGLWWAQ

QLDLGSHGLPTGCGKIGVLLDYEGDNMTFYNGNQVTEIKASFNGEKVFPFFYVGADVSLHLVP

>greenAnole_hTRIM_XP_008103837.1

MGKPQGKRTEFPPSLPACFSALAPPFRLVFFGFAFVSEQPLFLHSPPTALPSRLQQRGAMAVGGHPSARL

QDDATCSICLDYFQDPVMIIDCGHNYCRACISQCQGERSLCPRCRIPFPSDNLLPNRDLRNLVEAIRQLS

LQPLEKAARPKPAGGGPKMCEKHQEAAKLFCQEDKAFLCLICRESRAHKAHTALPIEEAAQEYRDQIQTF

LQKLAKERDELQRAVVYNRHHYQQLKSIVESAKQRIGPNFPKKESQTLMPLLNTCEANLQIVTEASNDLS

CLNTLITDIEEKSKQSSFGLLQGIGDLLNRCEQETRRPRRELKTILESIAENMDLVNTRLDQYAENTPKK

KWIKENVTLDAKTAHPRYFVSDDRKSVHWAKSRQDLPYSSKRFEFARCVLGKKGFTSGKHYWIVDVEDGD

YWAVGVAQESVDRDGELDFEPDEGIWAMARYDDQYKALTSPPTLLDLDYDPKEIQVSLNYDAGIVHFYDE

DKKPLYSFRSIDFEGEKVFPFFRIGDSSTSLYLL

>greenAnole_BTN_XP_008103839.1

MKSPVFPYASEMSCFWFIFMLFFVISPINDVSPAQLTVNGPLHPVTASLSEDIVLHCHLSPRTSAENMEI

KWFRSQNSSYVHLYHNGKEHLEKQQPEYQGRTELLTDGIEDGKIGLRILNVSFYDAGQYHCSVKNKSFHQ

EATLDIKVAALGSTPLISIERYQERGIYLVCRSRGWYPQPEVLWRNPSGKHLSFLAKETSQENNGLFEVQ

KDILLTEGSNQPLTCVVRNRLLNQEKGSTIHVADNFFPKIDPWAVGLCVSILVPVGLLIAILYLINRNRK

LITELSWRNMVVPIEKANVTLDPSTANKELILSADQKNVIQGFMWHNLPDHPWRFDVERCILGKEGFSSG

RHYWEVEVGEEGYWAVGVARESVRRKGRLSLDPDEGIWAVEKARVQTVQYQALTIPQTPLTLRKRPRKVG

IYLDYELGKVAFHDVNHKMHIFTFFAADFNGETIFPFLQVGIGCWLTVCP

>greenAnole_leftofBTN_psOnly_XP_062825617.1

MKVRGVFQTFGGLCCILWLSLCFLAELEGGGKVLASTPADVLFDPNTAHPNLHVSPNKKKVTWLPEPQNV

PDNPERFNATYCLLGTPRFTKGKHYWEVEYGGQREWVAGVARESVQRKGFVRMTPKEGIWQMGLWWLRRG

EMDSHPPPKHPAKIRISLDYEQGTVTFYLENKVNVQKVSFKGESVRPFFYVGPTVSLGL

**TRIMs for recombination analysis (ungapped, for GARD)**

>iguana_hTRIM

AATGTGATGCTGGATCCAGAAACAGCCCATCCTCGATACATTGTGTCTAAAGATCGGAAGAGTGTACACTGG---GGGGACAGCCGTAGGGACCTGCCTTACAACCCCAAAAGATTTGAGTATGCTCGCTGTGTGCTTGGAAAGAAGGGATTCACCTCAGGAAAGCATTATTGGACTGTGGATGTAGAGGATGGAGAATACTGGGCAGTGGGTGTGGCCTTAGAATCTGTGGACAGGGATGGGGAACTTGATTTTGAGCCTGATGAAGGGATCTGGGCTCTGGCACGTTAT---GATGATCAGTACAAAGCTCTTACTTCACCTCCAACTCTTCTTGACCCAGACAATGATCCGGTAGAGATCCAGATTTCTCTGAACTATGATGCTGGAACAGTGGCTTTTTATGAT---GAAGACAAGACCCGTCTCTTTATTTTCCGGTCCATTGATTTTGAAGGGGAGAAAGTGTTCCCTTTTTTCCGGATTTCTGATTCATCGACTTCTCTACGGCTGTGT---

>XM_067463364.1

AATGTGACGTTGGATGCAAAAACAGCCCATCCTCGGTTCTTTGTGTCTGATGATCGGAAAAGCGTACACTGG---GCGAAGACCCGCCAGGATCTGCCATACAGCTCCAAGAGATTTGAGTTTGCCCGCTGTGTGCTTGGGAAGAAAGGATTCACCTCTGGAAAACATTATTGGATTGTGGATGTAGAGGATGGTGACAACTGGGCGGTGGGTGTAGCCCAAGAGTCTGTGGACAGGGGAGGGAAACTTGATTTCGAGCCTGATGAGGGGATCTGGGCTATGGCACGTTAC---GATGATCAGTATAAGGCTCTTACTTCACCCCCTACCCTTCTGGATCTAGACTATGATCCAGTAGAGATCCAGGTTTCTTTGAACTATGATGCTGGAATAGTGCATTTCTATGAT---GAAGATAAGAAACCCCTCTTTACTTTCCGATCATTCGATTTTGAAGATGAGAAGGTCTTTCCTTTTTTCCGAATTGGTAATTCATCCACATCTTTGTACCTGCTC---

>XM_060270853.1

AACGTGGTGTTTGATCCGGAAACAGCCCATCCCCGATATATTGTGTCTTCTGATGGAAAATCAGTATGGTGG---GGAAAGGTCCGCCAGGACTACCCCTACAGTTCAATTAGGTTTGAATACGCCCGCTGTCTACTTGGAATGTGGGGATTCAGCTCAGGGAAACATTATTGGACGGTGGATGTTGAGGGTGTCAACCATTGGGCTGTGGGCGTAGCCAGAGAGTCTGTGGAGAGGGATAGAGAAATTAATTTTGAACCTGATGAAGGGATCTGGGCAGTGGGCTTTAGCCGCAATGATCAGTTCAAAGCTCTGACTTCACCACCTACCTATTTGGACCCAGAAGAGGGCCCAACACAAATCCAAGTTTCGTTGAACTACGAAGCAGGGACTGTGGCTTTTTATGATGCTGAAGATAAGACCCGCCTCTTTATTTTCGAGTCAATTGATTTTGAAGGGGAGGAAGTTTTCCCCTTTTTCCGGATTGTAGATCCATCAGTTTGCCTCCAGCTGTGCTAC

>XM_035105027.2

AATGTGGTGTTGGATCTGGAAACAGCCCATCCCCGGTATGTTGTGTCTTCTGATGGAAAATCAGTATGGTGG---GGAGATGTCCGTCAGGACTACCCCTACAATCCAAAGAGATTTCAATACGCTCGCTGTGTGCTTGGAACGCAGGGATTCAGCTCTGGGAAACATTATTGGACGGTGGATGTCGAAGATGTTGACAATTGGGCTGTAGGCGTAGCCAGAGAGTCTGTGGAGAGGGATAGAGAAATTGATTTTGAACCTGATGAAGGGATCTGGGCGTTGGGCTTTAGT---TATGATCAGTACAAAGCTCTGACTTCACCTCCTACCATTTTGGAACCAGAAGAGGATCCAATAGAGATCCAAGTTTCTTTGAACTATGAAGCAGGGACTGTGGCTTTTTATGATGCTGAAGATAAGACCCGCCTCTTTATTTTCAAGTCAATTGATTTTGAAGGGGAGGAAGTTTTCCCCTTTTTCCGGATTGTAGATTCATCAACTCCCCTCCAGCTGTGCTCT

>XM_035106984.2

AACGTGAAACTGGACCCCATGACGGCTCACCCTCGATTCATTGTGTTTGATGGCCAGAAAGCTGTTGCATGG---GGATCTGTCCGCCAGGAGCTGCCCTACAGCACCGGGCGATTTGACCCCACCCGCTGCGTGCTTGGCTGTCAGGGATTCACGTCCGGGAAGCACCATTGGACAGTAGATGTGAGCAAGGGAACTTTCTGGGCTGTGGGTGTAGCCAAAGAATCTGTGAAGAGGAAAGGACAGCTGAATATGGTCCCTGAGGAGGGAATCATGGCTGTGGGTTTTAAC---AACGGCCAGTACAAAACCTTAACGTCGCCCCCGACTGTTCTGAACCCAAGTAAACACCCCCAAAAGATTCAAATCCGTTTGGACTATGAAGGAGGAACGGTGGGGTTTTTTGATGCGGAAGAAAAGAAGCGCCTTGTTCTCTTC---AGTGTCAAATTTGTGTCC---AAAATATTTCCTTTCTTCCGAGTTGGGGACATGAACACTGTGCTCCGCCTGTGT---

>XM_060270855.1

AACGTGGTGTTTGATCCGGAAACAGCCCATCCCCGATATATTGTGTCTTCAGATAGAAAATCAGTATGGTGG---GGAAAGGTCCGTCAGGACTACCCCTACAGTTCAAATAGGTTTGAATACACTCGCTGTGTACTAGGAATGCGAGGATTCTACTCTGGGAAACATTATTGGACGGTGGATGTTGAGGATGTCGACCATTGGGCTGTGGGCGTAGCCAAAGAGTCTGTGGAGAGGGATAGAAAAATTGATCTTGAACCTGATGAGGGGATCTGGGCAGTGGGGTTTAGC---AATGATCAGTTGAAGGCTCTGACTTCACCTCCTACCCCTTTGGAAACAGAAGATGTCCCAACACAGATCCAAGTTAAGTTGAACTATGAAGCAGGGTCTGTGGCTTTTTATGATGCTGAAGATAAGACCCGCCTCTTTATTTTCCGGTCAATTGATTTTGAAGGGGAGGTAGTTTTTCCTTTTTTCCGGATCGTCAATTCATCAACTTGCCTCCAGCTGTACTCT

>XM_035106987.2

AACGTGGTGTTTGATCCGGAAACAGCCCATCCCCGATATATTGTGTCTTCAGATGGAAAATCAGTATGGTGG---GGAAATGTACGCCAGGACTACCCCTACAGTTCAATTAGGTTTGAATACGCCCGCTGTCTACTTGGAATGCGGGGATTCAGCTCAGGGAAACATTATTGGACAGTGGATGTTGAGGGTGTCAATCATTGGGCTGTGGGCGTAGCCAGAGAGTCTGTGGAGAGGGATAGAGAAATTAATTTTGAACCTGATGAAGGGATCTGGGCAGTGGGCTTTCGCCGCAATGATCAGTTCAAAGCTCTGACTTCACCACCTACCTATTTGGACCCAGAAGAGGGCCCAACACAAATCCAAGTTTCGTTGAACTACGAAGCAGGGACTGTGGCTTTTTATGATGCTGAAGATAAGACCCGCCTCTTTATTTTCGAGTCAATTGATTTTGAAGAGGAGGAAGTTTTCCCCTTTTTCCGGATTGTAGATCCATCAGCTTGCCTCCAGCTGTGCTAC

>XM_060239254.1

AATGTGTTGCTAGATCGAGAAACAGCCCATCCCCGATATGTTGTGTCTGAAGATTGGAAAAGTGTAAGATGG---GGGGACATCCGTCAGGAATTCCCCTACAACCCAAAGAGATTTCAATATATTCGTGGTGTTCTTGGATGTGAAGGATTCACTTCTGGAAAACATTATTGGACTGTGGATGTAGGAAATGGAGGTTGTTGGGCTGTGGGAGTGGCCAGAGAATCTGTGGAGCGGGAGGTGCAAATTGATTTTCAACCAGATGAGGGAATCTGGGCGATTGCACATTCC---AGTGATCAGTACAAAGCTCTAACCTCACCTCCTACTCTGCTGGATGTAGAAGATGAGCCAACACAGATCAAGGTTTGTTTGAACTATGAAAAGGGGACAGTTGCTTTTTATGATGTTTACGATGATACCCGCCTCTTCAAATTCTTTTCAGTTAATTTTGAAGAGGAGGAAGTTTTTCCTTTTTTCCGGATTGTTGATTCATCCACAGTTCTGGAATTGTGTTCT

>XM_060238766.1

TATGTGGTGCTGGACCAAGGTACCGCTCATCCTCGATTTGTTGTGTCTGATAATCGGAGAATGGTAACATGG---GGACCTTTTCGGCAGGAAGTGCCCTACAGCCCTGAGAGATTTGACCCAGCCCGCTGCATGCTTGGTTCTCATGGATTCTCATCAGGGAAACATCACTGGACAGTGAATGTGGAGGATGGAACTTTCTGGGCTGTAGGTGTGGCCCAAGAGTCTGTAAAAAGAAAAGGACAGTTCCATTTTGTGCCTCAGGAAGGGATTTGGGCTGTGGGCCTTCAT---AATGGTCAGTACAGAGCTCTGACTTCACCTCCTGACCTTTTGCTTCTAAATAGCCCCCTTAACAAGATCCAGGTTTGTCTGGATTATGAAGGAGGGACTGTAGCTTTCTGTGATGCTGAACAACAGCGTCAGTTCTATTGCTTT---CCAGCAAACTTCGAGGGAGAAAAACTCTTTCCTTTCTTCCGTGTTGGGGACATGAAGACTACCCTTTTCCTTTGC---

>XM_060239255.1

AATGTGTTGCTGGATCCAGAAACATCCCATCCCCGATATATTGTGTCTGCTGATCGGAAGACTGTAAAATGG---GGGAGTGCCCGTCAGCAGTTTCCCTACAACAAAAAGAGGTTTCAGTGTGTTCGCTGTGTTCTTGGCTCTGAGGGGTTCACTTCAGGAAAACATTATTGGACTGTGGATGTAGGAGATGGAGATTATTGGGCTGTGGGAGTGGTCAGAGAATCTGTGGAGCGGGAGGAGGAAATTCCTCTTGAACCTGATGAGGGAATCTGGGCAATTGGGCTTTAT---GACAATCAGTACAAAGCTCTGACCTCACCTCCTACTGTGCTAGAGGTAGAGGATGAGCCAACAGAGATCCAGGTATCTTTGAATTATGAAGCAGGGACAGTTAATTTTTATGATGCAGAAGATAGGACCCGTCTGTTCACTTTTCAGTCAGTTGATTTTGAAGGGGAGGAAGTTTTCCCTTTTTTCCGGATTGTGAATTCATCAACTGTCCTGACGATGTCCTAT

>XM_060239253.1

AATGTGTTGCTGGATCCAGAAACAGCCCATCCCCGATATGTTGTGTCTGAAGATCAGAAGACTGTAAGATGG---GGGAGTTTCCGACAGCAGTTTCCCTACGACCCAAAGAGATTTGAATGTATTCGCTGTGTTCTTGGCTCTGAGGGGTTCACTTCAGGAAAACATTATTGGACTGTGGATGTTGGAGATGGAGATTATTGGGCTGTTGGAGTGGCCAGAGAAAATGTGGAGCGGGAGGAGGAAATTCCTCTTGAACCTGATGAGGGAATCTGGGCAATTGGGCTTTAT---GACAATCAGTACAAAGCTCTGACCTCACCTCCTACTGTGCTAGAGGTAGAGGATGAGCCAACAGAGATCCAGGTATCTTTGAATTATGAAGCAGGGACAGTTAATTTTTATGATGCTGAAGATAAGACCCGTCTGTTCACTTTTCAGTCAGTTGATTTTGAAGGGGAGGAAGTTTTCCCTTTTTTCCGGATTGTGAATTCATCAACTGTCCTGACATTGTCCTAT

>XM_020777528.1

AATGTGCTGCTGGATCCTGAAACAGCCCACCCTCGATACATCATATCTGATGACTGCAAAAGAGTAGATTGG---GGAGATGTCCGTCAGGATTTCTTTTACAACCCAAAGAGGTTTCAGTATGCTCGTTGTGTGCTTGGAACAAGGGGATTCAAGTCAGGAAAACATTATTGGACAATAGATGTACAGGATGGACACCACTGGGCTGTGGGTGTGGCACGAGAGTCTGTGGAGAGGGATGTGGTGCTCTCTTTTGAACCCGATGAAGGGATCTGGGCCATGGCACGTTAT---TGTGATCAGTACAAAGCCATGACTTCACCTCCTACCATTCTGGAACTAGATGATGAGCCAACAGAGATTCAGGTTTCTCTGAACTGTGACGCTGGCACTGTAACTTTTTATGATGCTGAAGACTTAACCCGCCTCTATAAATTTGAGTGTGTTGATTTTGAAGGAGAGAAGGTTTTTCCTTTTTTTCGGATTTCTGATTCGTCTACTTCCCTCTCGATCACTTCC

>XM_020777535.1

GATATGACATTGGACGTCGCAACAGCTCATCCACGATTTGTTGTGTTTGAGGATCGGAAAACTGTCGCATGG---GGCCCTGTTCGTGAGGAGCTGCCTTACGATGCCAGAAGATTTGACCCTTCCCGCTGTGTGCTTGGTTCGCGAGGGTTCACCTCAGGGAAGCATCACTGGACAGTGGATGTGGACCAGGGAACTTTCTGGGCCGTGGGTGTGGCCCGAGAGTCTGTGAAGAGGAAAGGACAATTCACTATAATCCCCGCAGAGGGAATCTGGGCTGTTGGACTGTCC---AGTGGCCACTACAAAGCTCTGACTTCACCTCCTGTCATTCTGAGCCATTGCAAACCCATGAGAAAGATCCGGGTTTGTTTAGACTGTGAAGGAAAAACGGTGATGTTTTTTGATGCTGAAGAAAAGGAGGCCCCTGTTCTGTTCTCATTCCCAATTTCTGGATCAGGAAAATTTTTTCCTTTTTTCCGAGTCGGAGACATGAACACCGTCCTTCGCCTGTGT---

>XM_053295997.1

AATGTGTATCTGGATCCAGAAACAGCTCATCCTCGGTACATTGTGTCTCAAGACGGGAAAAGTGTCGAATGG---GGAGAAATCCGTCAGGATTTTCCCTACAACCCAAAGCGATTTCAGTATGCTCGTTGTGTTCTTGGTTGTGAGGGATTCACCTCAGGGAAACATTATTGGACAGTGAGCGTAGGGGATGGAGACTATTGGGCTGTGGGAGTGGCCAGAGAGTCTGTGGAGAGGGGGGAGGAAATTGATTTTGATCCTGATGAGGGGATCTGGGCTTTGGGACGTTAC---GATGATCAATATAAGGCTCTGACTTCCCCTCCTACTCTTTTGGAGCTAGACTATGAGCCAACAGAGATCCAGGTTTCTCTGAATTATGAAGCAGGAAAAGTGCATTTTTATGATGGAGAAGATAAGACCCGCATCTATACTTTC---GATGCTGATTTTGAAGGGGAGGCAATTTTCCCTTTTTTCCGGATTGTTGATTCATCAACTTGCCTCCAGCTCTGTTCT

>XM_053287828.1

AACGTTCTGTTGGATCCATTGACGGCGCACCCTCGATTTGTGGTGTCCGATAATCAGAAAAGTGTTGCATGG---GGATCTTTCCGCATGGACTTGCCCTACAACGCCAAGAGGTTTGACCCTGCCCGCTGTGTGCTTGGCTCTCAAGGATTCTCCTCGGGAAAACATCATTGGACGGTAGATGTGGCCAATGGAACTTTCTGGGCTGTGGGAGTGGCCAGAGAGTCTGTAACGAGAAAAGGACAACTCAGTTTTATTCCAGAGGAAGGAATCTGGGCTGTGGGTTTTTAC---AATGGTCAGTACAAAGCTCTAACTTCACCACCTACCTTTCTGAATCCCAGGAAACAACTTAAAAAGATCCAGATCTGTTTGGACTATGAAGAAGAGACTGTGACTTTTTGTGATGCTGAAGGAGGAGGGCGCCTTGTTAAATTTAAACCAGCTCATTTTGCAGGAAAGAAAATCTTTCCTTTCTTCCGTGTTGGGGATATAAACACGTGCATCGGGCTGTGT---

>XM_053302966.1

AATGTATTGTTGGCTCCAGAAACAGCCCATCCCCAATATATTGTGTCTGCAGATCGGAAATGTGTAAGATGC---AGAGATATCCTTCAGGAATTTCCCTCCAGCTCAAAGAGAATTGGAAATTTTCACTGTGTACTTGGAATGAAGGGATTCATCTCCGGAAAACATTATTGGAAGGTCAATGTAGGCCATGGGAAAAGTTGGGCTTTAGGAGTA---------TCTGTGGAGAGAAAAGGAGATGTCTGTATTATTCCCAAGGAGGGGATCTTTGCTCTGGGGTGTCAG---AATGGTTCCTATGTAGCATTCAGTTCCCAAGTCACTCCACTATACCTGAACAACAACCCAGAAGAGATCAAGGTTTCTTTGGATTATGAAGAAGCAACAGTGTCATTTTGTGATACATATGACTACCGTGTTCTTTATATGTTCCAATCTGTCAATTTTAAAGGTGAGAAAATCTTCCCTTTTTTCCAAGTTGCAGATATGGGCACAATTCTCAGTTTACAT---

>XM_054978680.1

AATGTGTTGTTTGATCCAGAAACAGCCCATCCCCGATATGTTGTGTCTGCAGATCGTAAAACTGTAAGATGG---GGGAGTATCCGTCAGGAATTCCCCTACAACCCAAAGAGATTTCACTATGTTCGCTGTGTTCTGGGCTGCAAGGGGTTCACTTCAGGGAAACATTACTGGACCGTGGATGTCGGAGATGGAGATTACTGGGCTGTGGGAGTGGCCAGAGAATCTGTGGAGCGGGAGGAGGAAATTGAGTTTGAACCTGATGAAGGAATCTGGGCTCTTGGGCTTTAT---AATGATCAGTACAAAGCTCTGACTTCACCTCCTACCCTACTGGATGTAGAGGATGAACCAACACAGATCCAGATTTCTTTGAACTATGAAGCGGGAACAGTTGCTTTTTATGATACTGAAGATAATACCCGTCTCTTCACTTTCCAGTCAGTTGATTTTGAAGGGGAGGAAATTTTCCCTTTTTTCCGGATTGTTGATTCATCAACTGTCCTGAAGTTGTGTTCT

>XM_054976459.1

TCATTGACATTTGATGTGGAGACAGCTCATCCTCGTCTGGTGGTCTCCAGAAGTGGCAAAAGCGTGAGATGGGGAGGAGATATCCAACATGCCCTGCTTCCTGGGCCCTGGAGATTTGACCATAGTCGCTGCTTGCTGAGTAGCCAAGGATTCACCTCTGGGTATCACTGCTGGATAGTAGAGGTGGTCAAGGAGGGGCCTTGGGCTATTGGGGTTGCTCTAGAGTCTGTGCAGAGGAAGGGGTCAGTCAACTTGATCCCTAGGGAAGGAATTTGGGCCATGGCATTTAAC---AACAGCAAATACTTGGTTCATACATCTCCCCCTACCCCTCTGATTCTGAGCTCTTCACCCAGAATAATACAAGTGTATCTGCATTATGAAAATGGGAAGGTTGTATTTATGGACTTCCATAGTAAGACTGTACTGTATGAATTCCTGTCTGCCTCTTTCTTGGGGCAGAAAGTCTATGCTTTCTTCCGTGTTGGCGGACCATCAGCTCATCTCAGATTGTGCTGT

>XM_054976458.1_cds1

AATGTGATGCTGGACCAGGAGACGGCTCATCCTAGATTTCTTGTGTCTGATGATCGGAGAAGCGTGTCATGG---GGATCGTTCCGGCAGGAGGTGCCCTACAGCCCCAAGAGGTTTGAGCCGGCCCGCTGTGTGCTTGGCTCTCATGGATTCATGTCAGGGAAACATCACTGGACGGTGGATGTGGAGGATGGAACTTTCTGGGCTGTGGGTGTGGCCCAGGAGTCTGTGCAAAGAAAAGGAAAGTTTAATTTTGTACCTGAGGAGGGGGTTTGGGCTGTGGGCTTTTCA---AACGGTCAGTACAAAGCTCTAACTTCACCTCCTGACTTTCTAATGTTACTTACCCACCCTAATAAGATCCAGGTTTATTTAGACTATGAAAACGAGGCTGTCACTTTCTGGGATCCAGAAGAGCATGCTGAGTTGTATCGCTTCGAATCAGCAAACTTTAAGGGGAAAACACTTTTTCCTTTCTTCCGGATTGGGGACATGAAGACTAGCCTTTTGCTATGT---

>XM_053379005.1

AACGTGGTGTTTGATCCGGAAACAGCCCATCCCCGATATATTGTGTCTCCAGATGGGAAAGCGTTATGGTGG---GGAAATGTCCGTCAGGACTACCCCTACAGTTCAATTCGGTTTGAATACGCCCGCTGTGTACTTGGAATGTGGGGATTCAGCTCAGGGAAACATTATTGGATGGTGGATGTCCGAAATGTCAACCATTGGGCTGTGGGCGTAGCCAGAGAGTCTGTGGAGAGGGATAGGGAAATTAATTTTGAACCCTATGAAGGGATCTGGGCAGTGGGCTTTAGCCGCCATGATCAGTTCAAAGCTCTGACTTCACCTCCTATCTATTTGGACCCAGAAGAGGGCCCAGCACAAATCCAAGTTTCTTTGAACTATGACGCAGGGAAAGTGTCTTTTTATGATGCTGAAGATAACACCTGCCTCTTTATTTTCGACTCAATTGATTTTGATGGGGAGGAAGTTTTTCCCTTTTTCCGGATTGTCGATTCATCAGCTTGCCTCCAGCTGTGCTAT

>XM_053378956.1

AATGTGGTGTTGGATCCGGAAACAGCCCATCCCCGGTATATTGTGTCTTCAGATGGAAAATCAGTACGGTGG---GGAGATGTCCGCCAGGACTACCCCTACAACCCAAAGAGGTTTGAATACGCCCGCTGTGTGCTTGGAACGCAGGGATTCAGCTCTGGGAAACATTATTGGACAGTGGATGTTGAGGATATTGACCACTGGGCTGTGGGCGTAGCCAGTGAGTCTGTGGAAAGGGATAGAGAAATTGATTTTGAACCTGATGAAGGGATCTGGGCGATGGGCTTTAGT---TATGATCAGTACAAAGCTCTGACTTCACCTCCTACCGTTTTGGAGCCAGAAGAGGATCCAGTAGAGATCCAAGTTTCTTTGAACTATGAAGCAGGAACTGTGGCTTTTTATGATGCTGAAGATAAGACCCGCCTCTTTATTTTCCAGTCAATTGATTTTGAAGGGGAGGAAGTTTTCCCCTTTTTCCGGATTGGAAATTCATCAGCTCCCCTCCAGCTGTGCTCT

>XM_053378966.1

AACGTGACACTGGACCCCATGACGGCTCACCCTCGATTTGTTGTGTTTGATGGTCAGAAAGCTGTCGCATGG---GGATCTGTCCGCCAGGAGCTTCCCTACAGCACCGGACGATTTGACCCCACCCGCTGCGTGCTTGGCTGTCAAGGATTCACCTCAGGGAAGCATCATTGGACAGTGGATGTGAGCAAAGGAACTTTCTGGGCTGTGGGTGTAGCCAAAGAATCTGTGAAGAGGAAAGGACATCTGAATATTGTCCCTGAGGAGGGAATCATGGCTGTGGGTTTTAAC---AACGGTCAGTACAAAACCTTAACGTCGCCCCCGACCATTCTGAACCCAAGTAAACACCCCCAAAAGATTCAAATCCGTTTGGACTATGAAGGAGGAACTGTGGGGTTTTTTGATGCGGAAGAAAACAAGCGCCTTGTTCTCTTT---AGTGTCAAATTTGTGTCA---AAAATATTTCCTTTCTTCCGAGTTGGGGACATGAACACTGTGCTCCGCCTGTGT---

>XM_033141496.1

AATGTGGTGTTGGATCCGGATACAGCCCATCCCAGATGTATTGTGTCTTCTGATGGAAAATCAGTACGGTGG---GGAAAGGTCCGTCAGGACTACCCCTACAGTCCAAATAGGTTTGAATATGCTCGCTGTGTACTAGGAATGCAAGGATTCAGCTCCGGGAAGCATTATTGGACGGTGGATGTTGAGGATGTTGACCATTGGGCTGTGGGTGTAGCCAAAGAGTCTGTGGAGAGGGATAGAGAAATTGATTTTGAACCTGAAGAAGGGATCTGGGCAGTGGGGTTTAGC---AATGGTCAGTTGAAAGCTCTGACTTCACCTCCTACCCCTTTGGAACCAGAAGAGGATCCAAGAGAGATCCAAGTTTCTTTGAACTATGAAGCAGGGACTGTGGCTTTTTATGATGCTGAAGATAAGACCCGCCTCTTTATTTTCCGCTCAATTGATTTTGAAGGGGAGGAAGTTTTTCCCTTTTTCCGGATCGTCAATTCGTCAACTTGCCTCCAGCTGGGCTCT

>XM_033141497.1

AATGTGGTGTTGGATCCGGATACAGCCCATCCCAGATGTATTGTGTCTTCTGATGGAAAATCAGTACGGTGG---GGAAAGGTCCGTCAGGACTACCCCTACAGTCCAAATAGGTTTGAATATGCTCGCTGTGTGCTTGGAATGCAAGGATTCAGCTCCGGGAAACATTATTGGACGGTGGATGTTGAGGATGTTGACCATTGGGCTGTGGGCGTAGCCAGAGCATCTGTGGAGAGGGATAGAGAAATTGACTTTGAACCTGATGAAGGGATCTGGGCGATGGGCTTTAGT---TATGATCAGTACAAAGCTCTGACTTCACCTCCTACCATTTTGGAACCAGAAGAGGATCCAGTAGAGATCCAAATTTCTTTGAACTATGAAGCAGGGACTGTGGCCTTTTATGATGCTGAAGATAAGACCCGTCTCTTTATTTTCCGGTCAATTGATTTTGAAGAGGAGGAAGTTTTTCCTTTTTTCCGGATCGTCGATTCGTCAACTTGCCTCCAGTTGGGCTCT

>XM_033141533.1

AACGTGACACTGGACCCCATGACGGCTCACCCTCGATTTGTTGTGTTTGATGGTCAGAAAGCTGTCGCATGG---GGATCTGTCCGCCAGGAGCTTCCCTACAGCACCAGACGATTTGACCCCTCCCGCTGCGTGCTTGGCTCTCAAGGATTCACCTCGGGGAAGCATCATTGGACAGTGGATGTGAGCAAGGGAACTTTCTGGGCTTTGGGTGTAGCCAAAGAATCCGTGAAGAGGAAAGGGCAGCTGAATATCGTCCCTGAGGAGGGAATCATGGCTGTGGGTTTTAGC---AACGGTCAGTACAAAACCTTAACGTCGCCCCCGACTTTTCTGAACCCAAGCAAAGACCCCCGAAAGATTCAGATCCGTTTGGACTATGAAGGAGGAACATTGGGGTTTTTCGATGCAGAAGAAAATAAGCGCCTCGTTCTCTTT---ACTGTCAAATTTGTGTCA---AAATTATTTCCTTTCTTCCGAGTTGGGGACATGAACACTGTGCTTCGACTGTGT---

>XM_042455007.1

AACGTGATGCTGGATCCAGAAACAGCCCATCATCGGTACATCGTGTCTCAAGATCTGAAAAGTGTACATTGG---GCAAAAAGCCGTCAGGATCTGCCCTACAGTTCCAAAAGATTTGAGGGTACTCGCTGTGTGCTTGGAAGGAAGGGCTTCACCTCAGGAAAACATTATTGGATAGTGGATGTAGAGAGTGAAGGATACTGGGCAGTAGGTGTAGCCCAAGGATCTGTGGACAGGGAGGAGAAACTTGATATTGAACCAGATTCGGGAATCTGGGCTCTGGCACGTTAT---AATGATCAGTACAAAGGTCTTACATCACCACCTACCCTTCTGGATCTAAACTATTATCCAGTAGAGATCCGGGTGTATCTGAACTATGATGATGGAACAGTGGATTTTTATGAT---GAAGATATGACCCGTCTCTTTATTTTCCAGTCAATTGATTTTCAAGGGGACGAAGTCTTCCCTTTTTTCCGGATTTTTAAGCCATCGTCTTCTCTACGCCTATGT---

>XM_008105630.3-trunc

AATGTGACGCTGGATGCAAAAACAGCCCATCCTCGGTACTTTGTGTCTGACGATCGGAAAAGCGTACACTGG---GCGAAGAGCCGCCAGGATCTGCCATACAGCTCCAAAAGATTTGAGTTTGCCCGCTGTGTGCTTGGGAAGAAGGGATTCACCTCTGGAAAACATTATTGGATCGTGGATGTAGAAGATGGTGACTATTGGGCGGTGGGCGTAGCCCAGGAATCTGTGGACAGGGATGGGGAGCTTGATTTTGAGCCTGATGAAGGAATCTGGGCTATGGCACGTTAC---GACGATCAGTACAAAGCTCTTACTTCACCCCCTACTCTTCTAGATCTAGACTATGATCCAAAAGAGATCCAGGTCTCTTTGAACTATGATGCTGGAATAGTGCATTTCTATGAT---GAAGATAAGAAACCCCTCTATTCCTTCCGATCAATCGATTTTGAAGGGGAGAAAGTCTTCCCTTTTTTCCGCATTGGTGATTCATCCACATCTTTGTACCTGCTC---
